# Defining neuronal responses to the neurotropic parasite *Toxoplasma gondii*

**DOI:** 10.1101/2025.03.31.645603

**Authors:** Hannah J. Johnson, Joshua A. Kochanowsky, Sambamurthy Chandrasekaran, Christopher A. Hunter, Daniel P. Beiting, Anita A. Koshy

**Affiliations:** Neuroscience Graduate Interdisciplinary Program, University of Arizona, Tucson, Arizona, USA; BIO5 Institute, University of Arizona, Tucson, Arizona, USA; Department of Immunobiology, University of Arizona, Tucson, Arizona, USA; Department of Pathobiology, School of Veterinary Medicine, University of Pennsylvania, Philadelphia, PA, USA; Department of Neurology, University of Arizona, Tucson, Arizona, USA

## Abstract

A select group of pathogens infect neurons in the brain. Prior dogma held that neurons were “defenseless” against infecting microbes, but many studies suggest that neurons can mount anti-microbial defenses. However, a knowledge gap in understanding how neurons respond *in vitro* and *in vivo* to different classes of micro-organisms remains. To address this gap, we compared a transcriptional dataset derived from primary neuron cultures (PNCs) infected with the neurotropic intracellular parasite *Toxoplasma gondii* with a dataset derived from neurons injected with *T. gondii* protein *in vivo*. These curated responses were then compared to the transcriptional responses of PNCs infected with the single stranded RNA viruses West Nile Virus (WNV) or Zika Virus (ZKV). These analyses highlighted a conserved response to infection associated with chemokines (*Cxcl10, Ccl2*) and cytokines (interferon signaling). However, *T. gondii* had diminished IFN-α signaling *in vitro* compared to the viral datasets and was uniquely associated with a decrease in neuron-specific genes (*Snap25*, *Slc17a7*, *Prkcg*). These data underscore that neurons participate in infection-induced neuroinflammation and illustrate that neurons possess both pathogen-specific and pathogen-conserved responses.

**Importance:** Though neurons are commonly the target of pathogens that infect the CNS, few datasets assess the neuronal response to infection. This paucity of data is likely because neurons are perceived to have diminished immune capabilities. However, to understand the role of neurons in neuroinflammation and their immune capabilities, their responses must be investigated. Here we analyzed publicly accessible, neuron-specific datasets to compare neuron responses to a eukaryotic pathogen versus two Orthoflaviviruses. A better understanding of neuron responses to different infections will allow us to develop methods for inhibiting pathways that lead to neuron dysfunction, enhancing those that limit pathogen survival, and mitigating infection-induced damage to the CNS.

## Introduction

A select number of microbes (e.g. measles virus, *Toxoplasma gondii*) infect the central nervous system (CNS). For many of these infections, neurons are the CNS cell that is primarily infected^1–3^. Until relatively recently, dogma suggested this neuronal predominance arose from neurons lacking cell intrinsic immune responses. Over several decades, work focusing on viral-neuron interactions established that neurons have cell intrinsic responses, though these responses can differ from other cell types and even between neuron subtypes^4–7^. These studies raise the question of whether the cellular immunity of neurons varies by context and/or pathogen. The eukaryotic intracellular parasite *Toxoplasma gondii* is a non-viral microbe with a tropism for neurons^8^ and broad natural host range, including rodents and humans^9^. During infection, parasites invade the CNS where they can infect multiple cells types, but, in neurons, a portion of parasites switch to a slow growing stage that forms tissue cysts^1,10,11^. These tissue cysts cause a persistent, asymptomatic infection, potentially for the lifetime for the host^10–12^. Recent studies suggest that the immune system can recognize infected neurons which contributes to local control of *T. gondii in vivo*^13–16^. *In vitro* studies have shown that neurons can be activated by IFN-γ to limit parasite growth^17^. Like most intracellular microbes, all of which depend upon the host cell for survival, *T. gondii* highly manipulates its host cell through the secretion of effectors proteins. Most of the studies that define how these effector proteins manipulate cells were done *in vitro* in fibroblasts and immune cells such as macrophages^18,19^. While such studies have revealed fundamental aspects of *T. gondii*-host cell biology, they will have missed neuron-specific effects or effects only triggered during *in vivo* infection. The importance of understanding these nuances is highlighted by studies showing that outcomes of *T. gondii*-host cell interactions can vary by *T. gondii* strain and host cell^20–23^.

We previously tried to address this gap by using laser capture microdissection (LCM) in combination with our *T. gondii*-Cre system^24^. In this system, we use parasites that express a *T. gondii*::Cre recombinase fusion protein (ROP::Cre) to infect Cre reporter mice that express a green fluorescent protein (GFP) only after Cre mediated recombination. Because the ROP::Cre protein is introduced into the host cell concomitantly with other early effectors (ROPs) and before full invasion, neurons injected with the ROPs will express GFP even if they cleared the parasite or were never invaded (i.e., aborted invasion)^25–27^. We then used LCM and RNA-seq to isolate, pool, and transcriptionally profile the somas of *T. gondii*-injected (GFP^+^) neurons (TINs). Though a small area centered on TINs’ somas was captured, these transcriptional data still contained immune cells transcripts^24^, making it difficult to distinguish which differentially expressed genes (DEGs) or pathways were derived from neurons, immune cells, or both.

In this study, we sought to identify neuron-specific responses to *T. gondii* by comparing RNA-seq datasets from our *in vivo* data with a newly generated *in vitro* dataset from *T. gondii*-infected primary neuron cultures (PNCs). This analysis revealed a set of conserved pathways driven by chemokines such as *Ccl2* and *Cxcl10.* The comparison to previously published transcriptomes of West Nile Virus (WNV)-infected and Zika Virus (ZKV)- infected PNCs^7^ revealed pathways that were conserved between these datasets and others that were pathogen dependent. For example, *T. gondii* datasets revealed a decrease in neuron-specific genes (e.g., *Snap25*, *Slc17a7*, *Prkcg*) that were unchanged in virally infected neurons. Conversely the type I IFN (IFN-α) response pathway was upregulated by WNV and ZKV and to some extent by *T. gondii in vivo*, but not by *T. gondii in vitro*. In summary, the ability to compare the *in vitro* and *in vivo* response of neurons to infection highlights that neurons have intrinsic, microbe-specific responses that are modulated *in vivo*.

## Results

### Conserved neuron response genes and pathways in *T. gondii* infection models

Transcriptomic data from neurons in two *T. gondii*-infection conditions— neurons in tissue sections isolated with laser capture microdissection (LCM)^24^ and cortical primary neuron cultures (PNCs) (**Fig. 1**)—were compared to find common neuron response pathways. Briefly, in a previous report, we combined transgenic parasites that secrete Cre recombinase with a mouse strain that expresses a Cre-sensitive GFP reporter which allowed us to isolate *T. gondii*-injected neurons (GFP^+^NeuN^+^) by LCM ^27^. The RNA from these neurons was isolated, sequenced, and compared with neurons from uninfected mice (**Fig 1A**).

**Figure 1.**
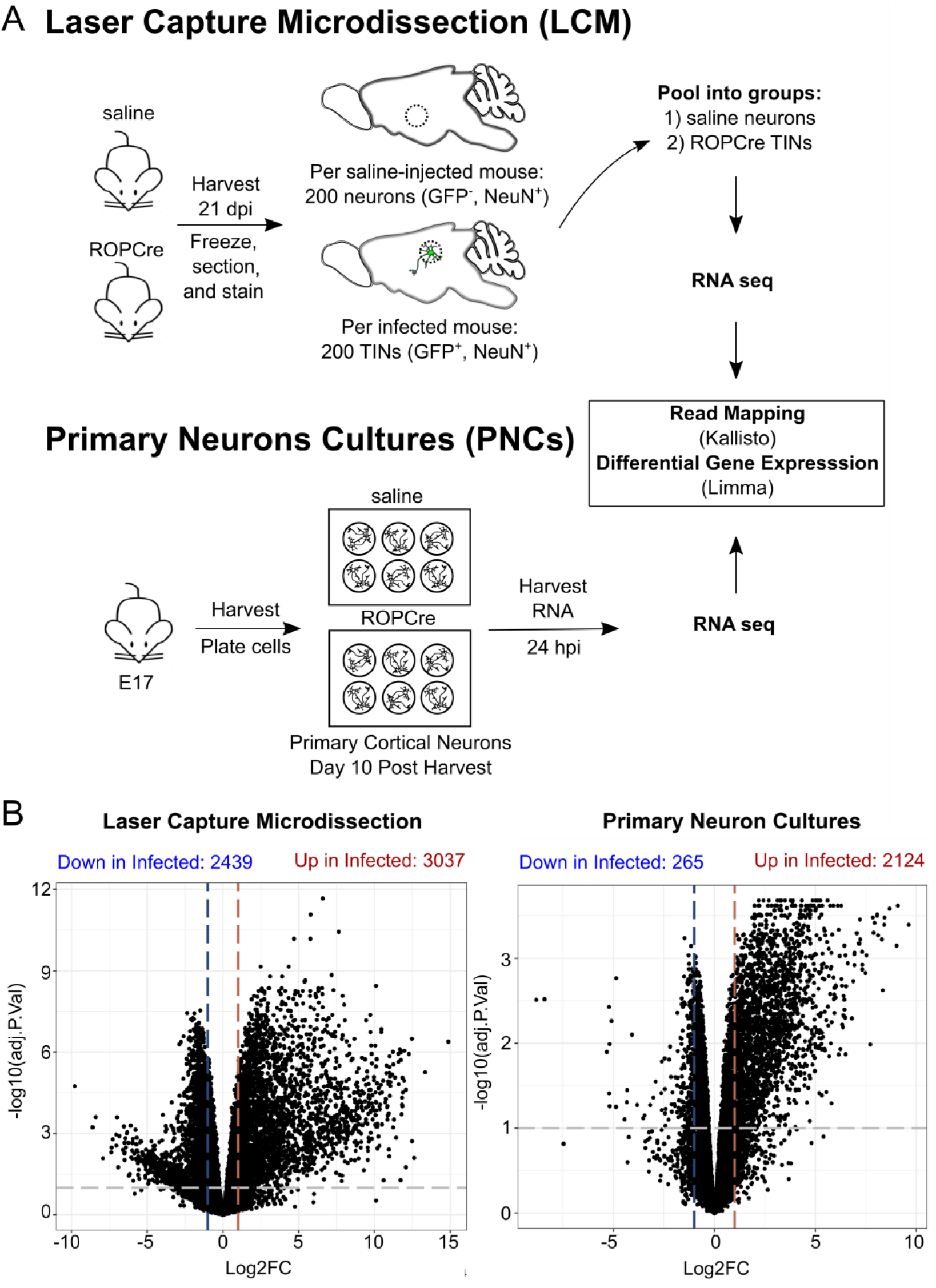
*T. gondii*-infected neurons from *in vitro* (PNC) and *in vivo* (LCM) systems were captured and analyzed for differentially expressed genes. (**A**) Experimental schematic of neurons captured by laser capture microdissection and infected primary murine neuronal cultures. (**B**) Volcano plots of differentially expressed genes in both datasets. Horizontal bars indicate adjusted p. values ≤ 0.1, vertical bars indicate Log2FC ≥ 1 for up and downregulated genes.

In a separate study, PNCs infected with *T. gondii* for 24 hours were used to assess how infection altered the neuronal transcriptome. Reads were filtered, normalized, and represented as counts per million (CPM), shown in **Fig. S1A-C**. As expected, Principal Component Analysis of the LCM and PNCs datasets revealed marked differences between infected and uninfected controls (**Fig. S1D, E**), with over 2100 differentially expressed genes (DEGs) compared to uninfected controls (FDR ≤ 0.1, Log_2_ Fold Change (FC) ≥ 1) (**Fig. 1B**, **Table 1, 2**).

**Table 1.**
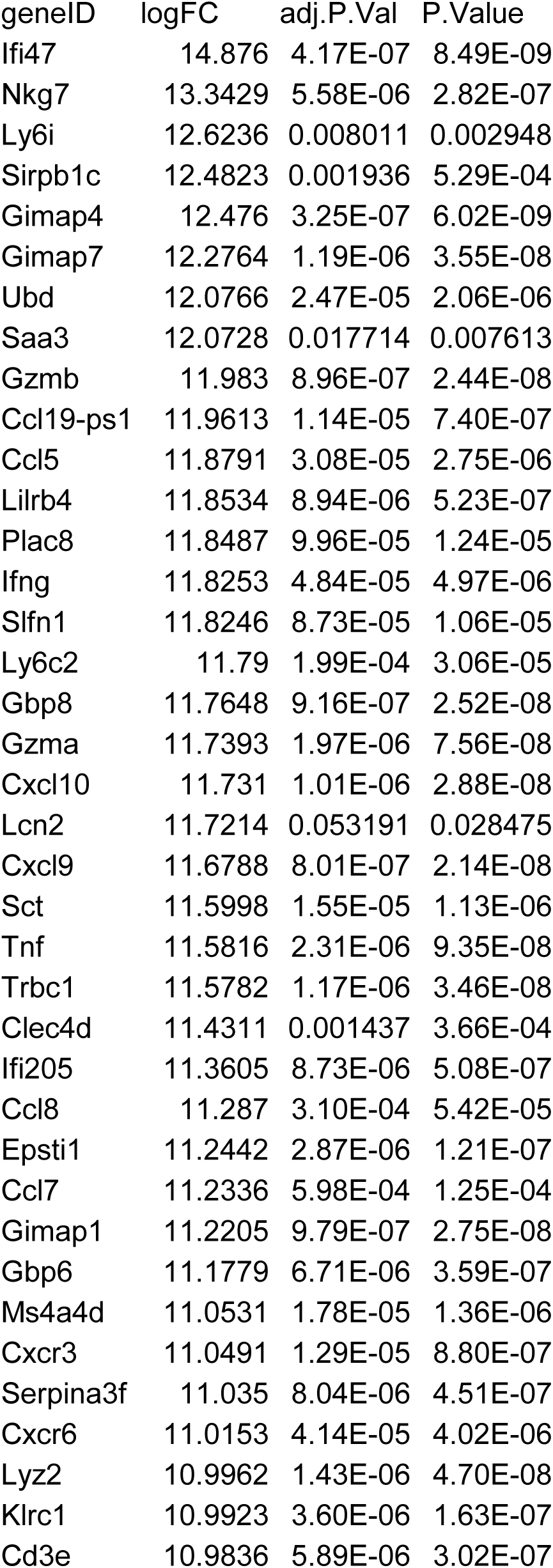
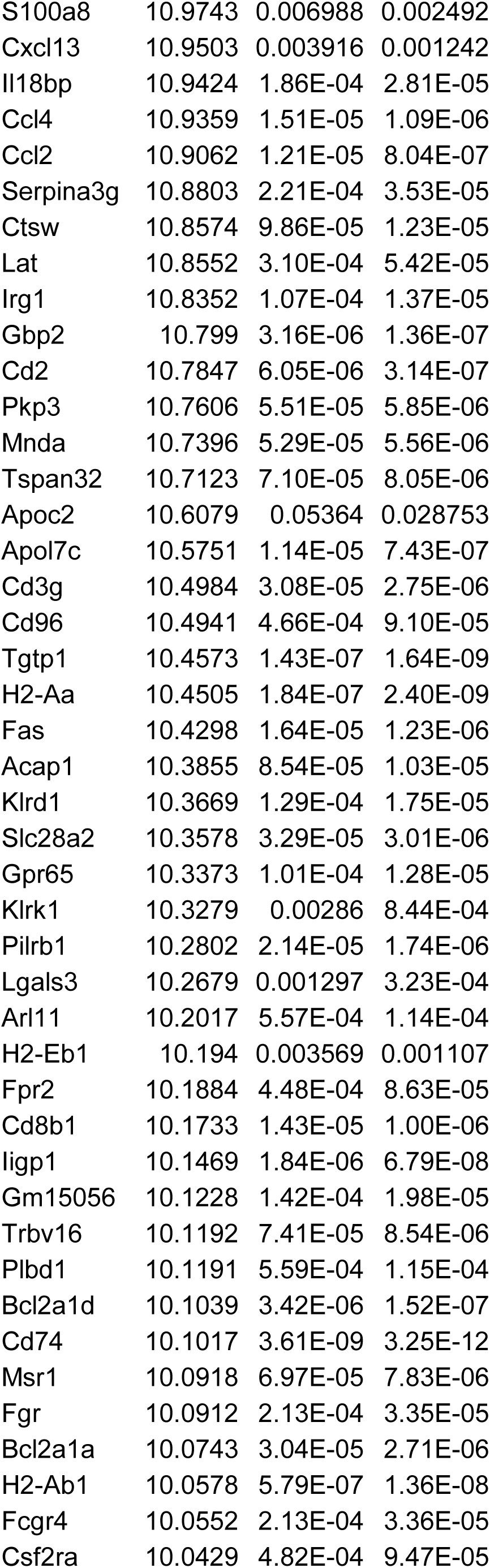
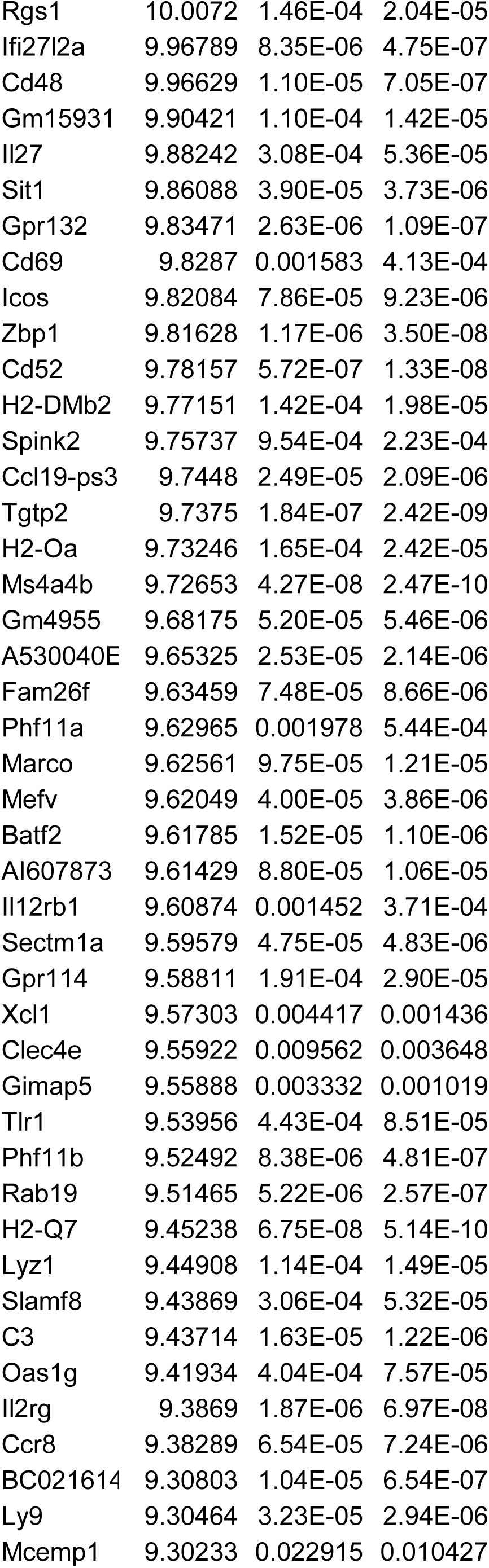
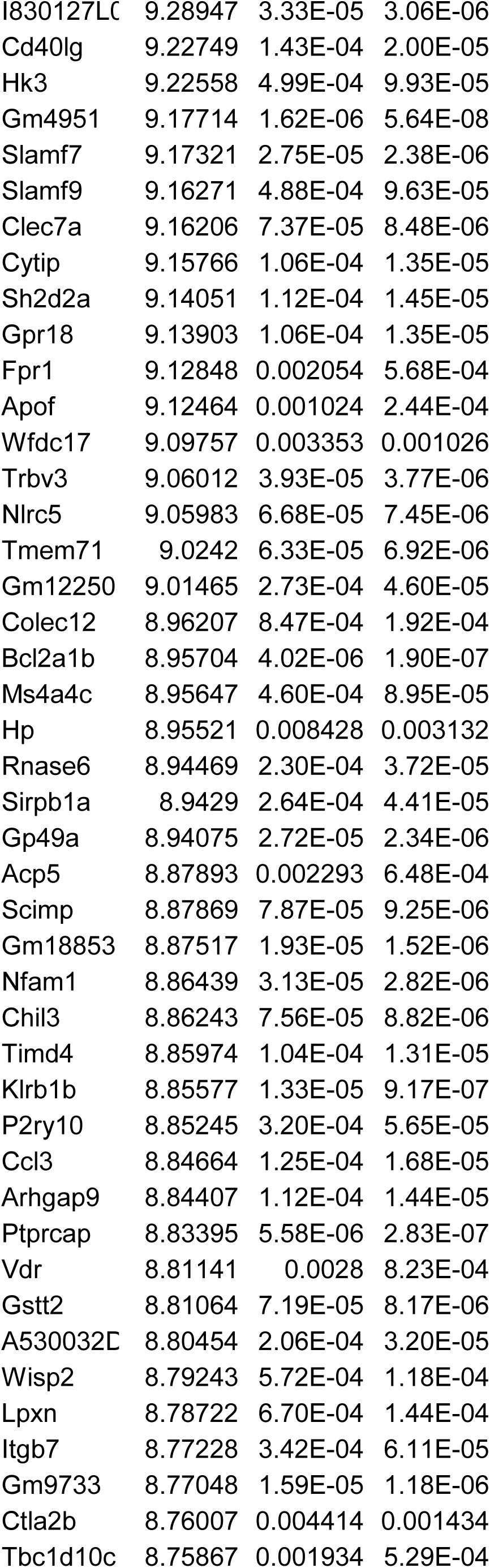
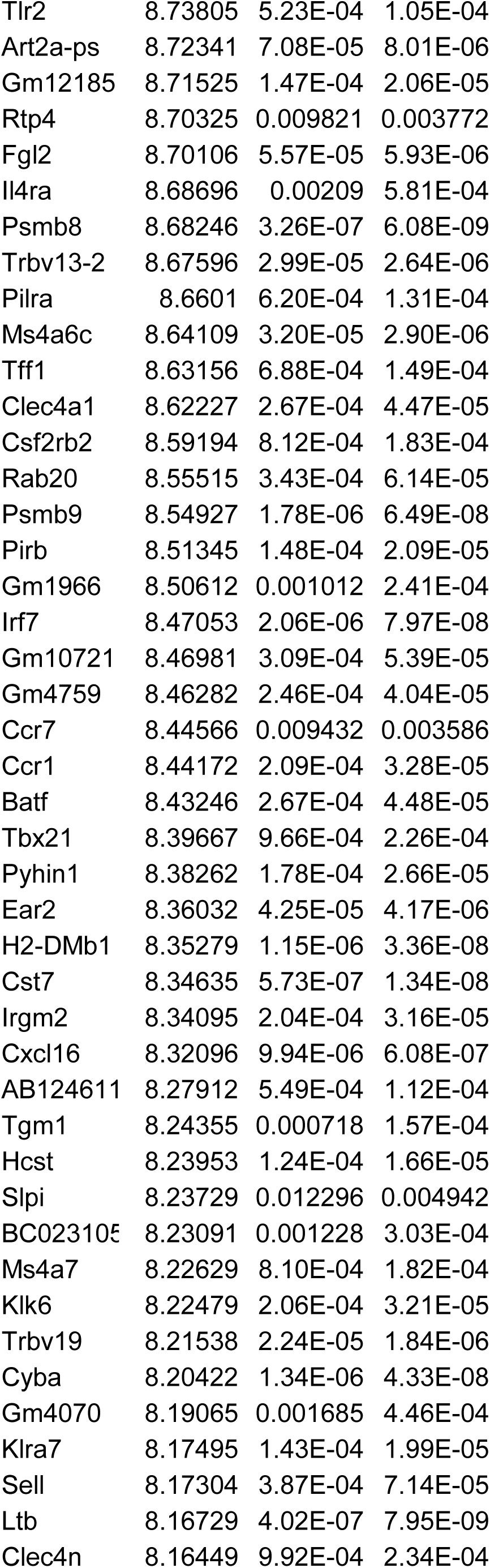
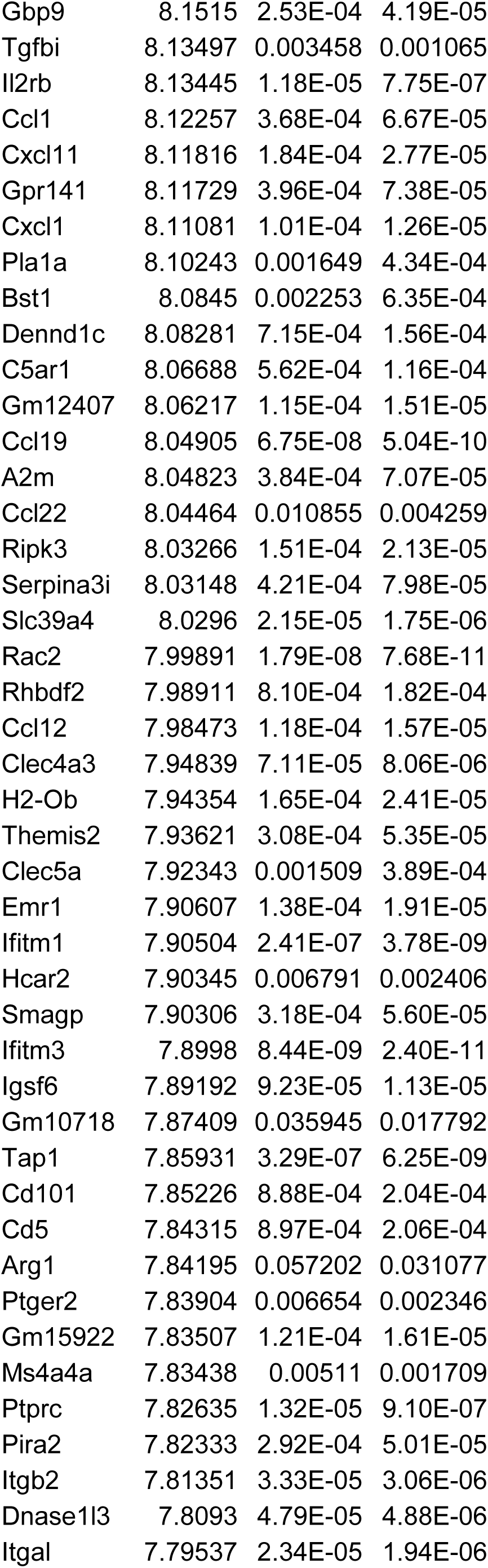
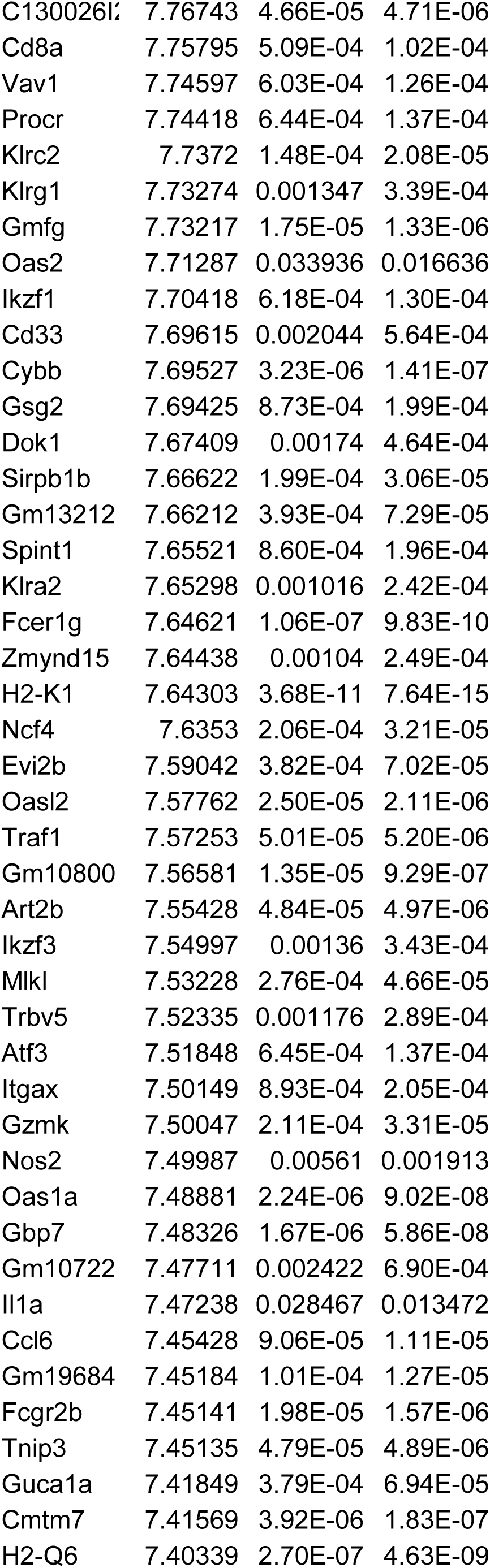
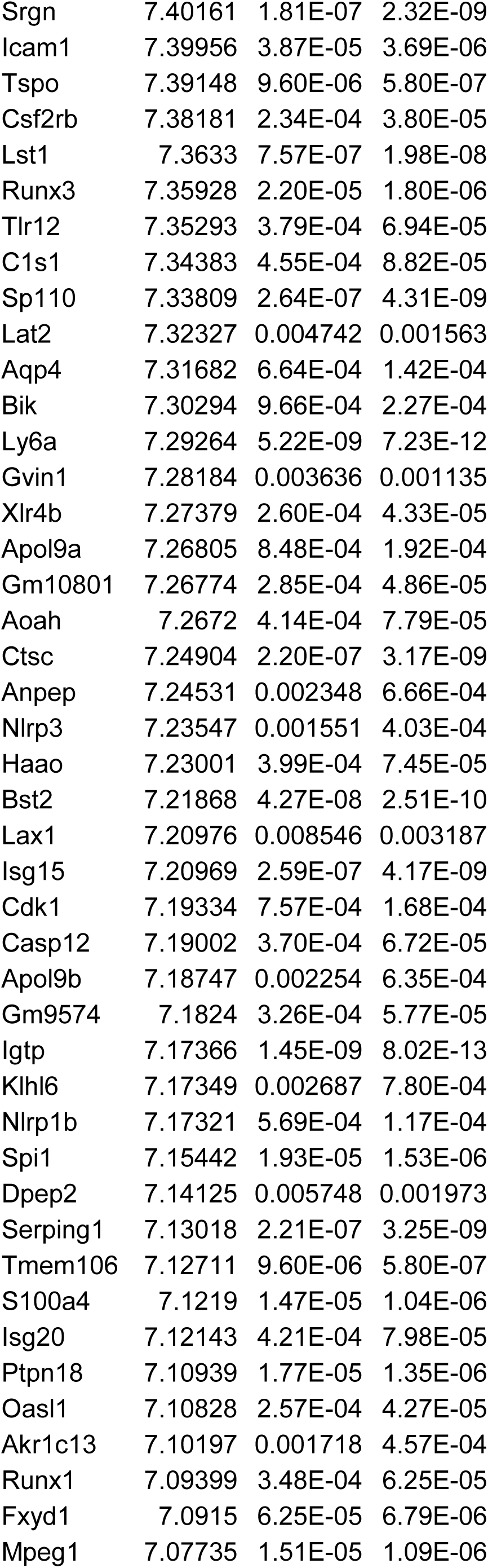
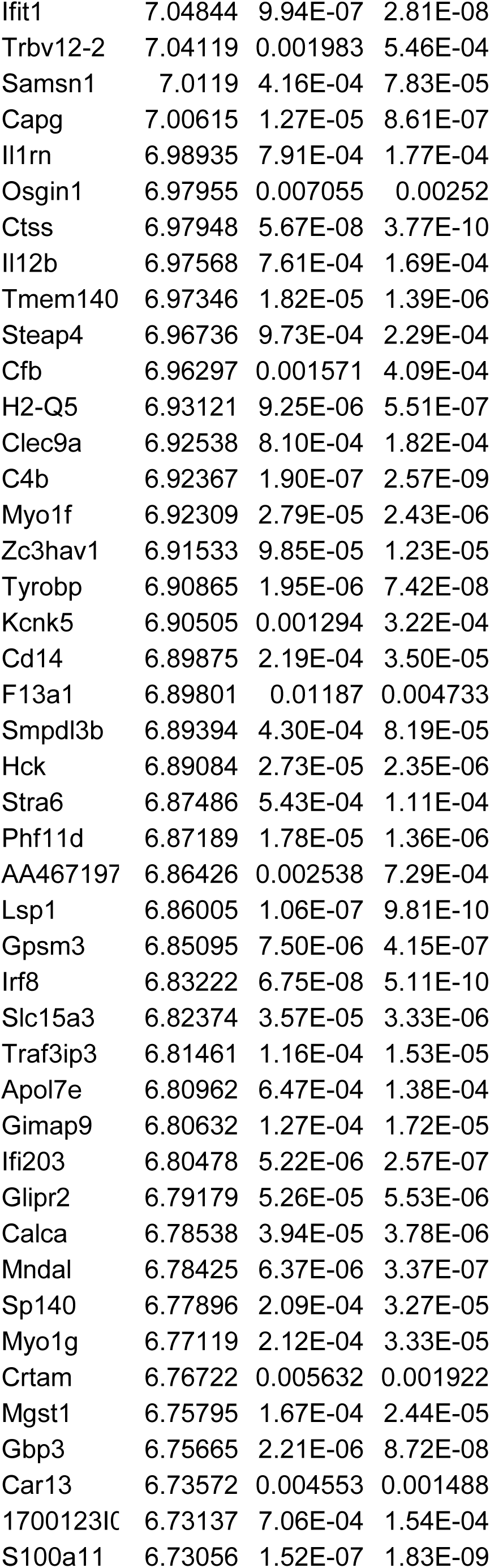
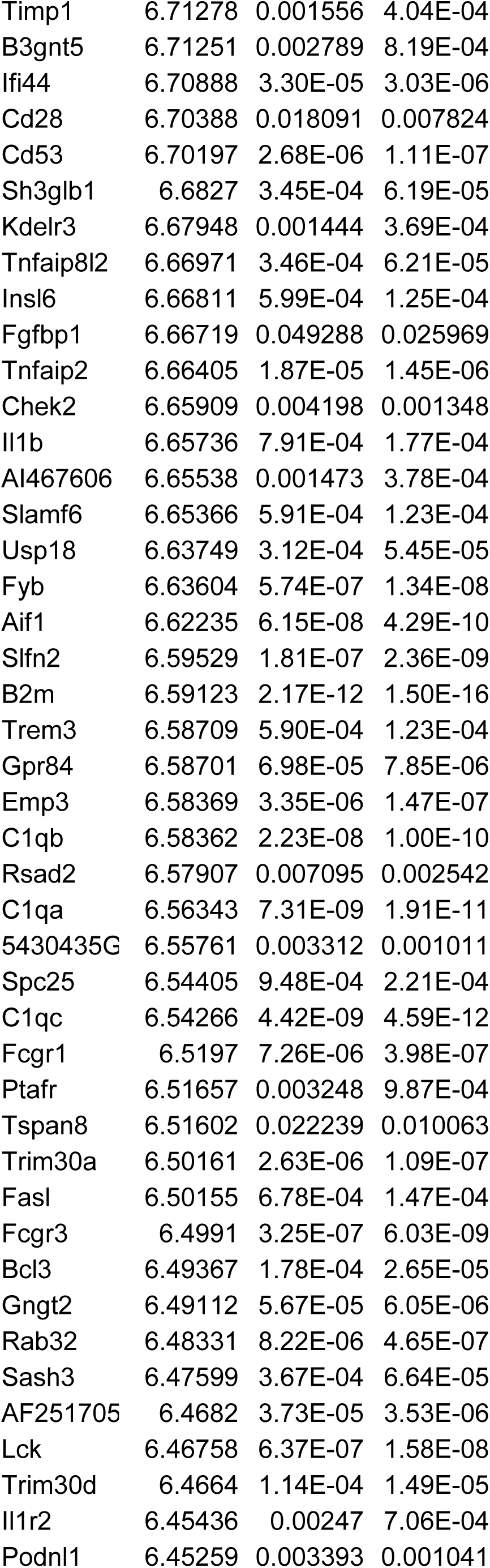
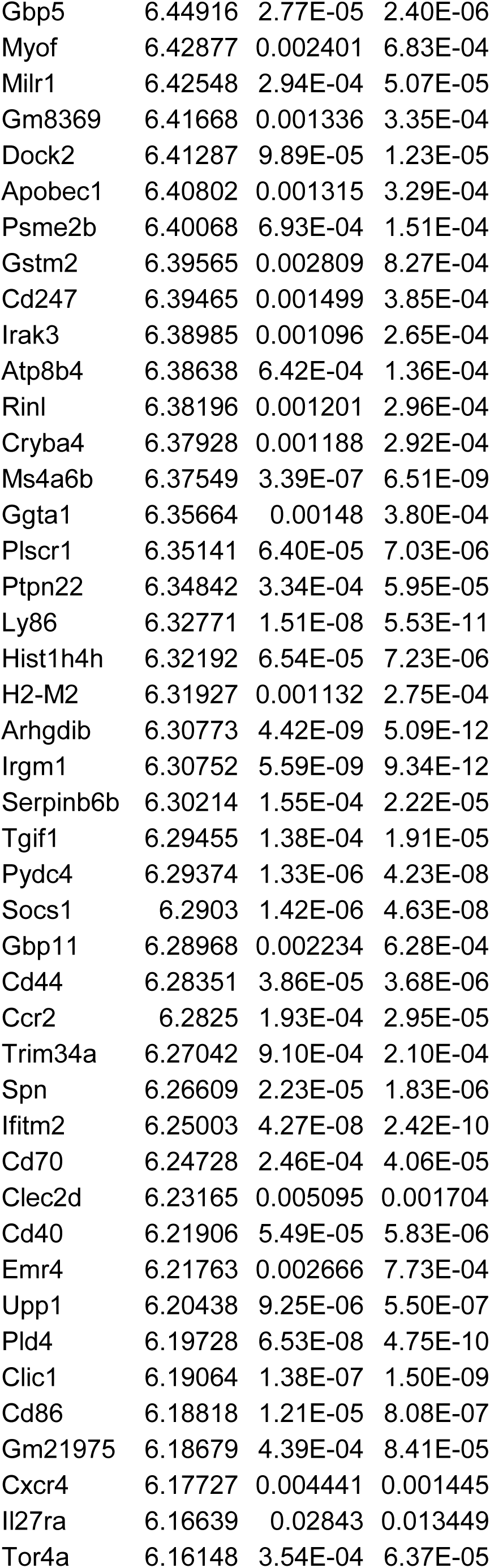
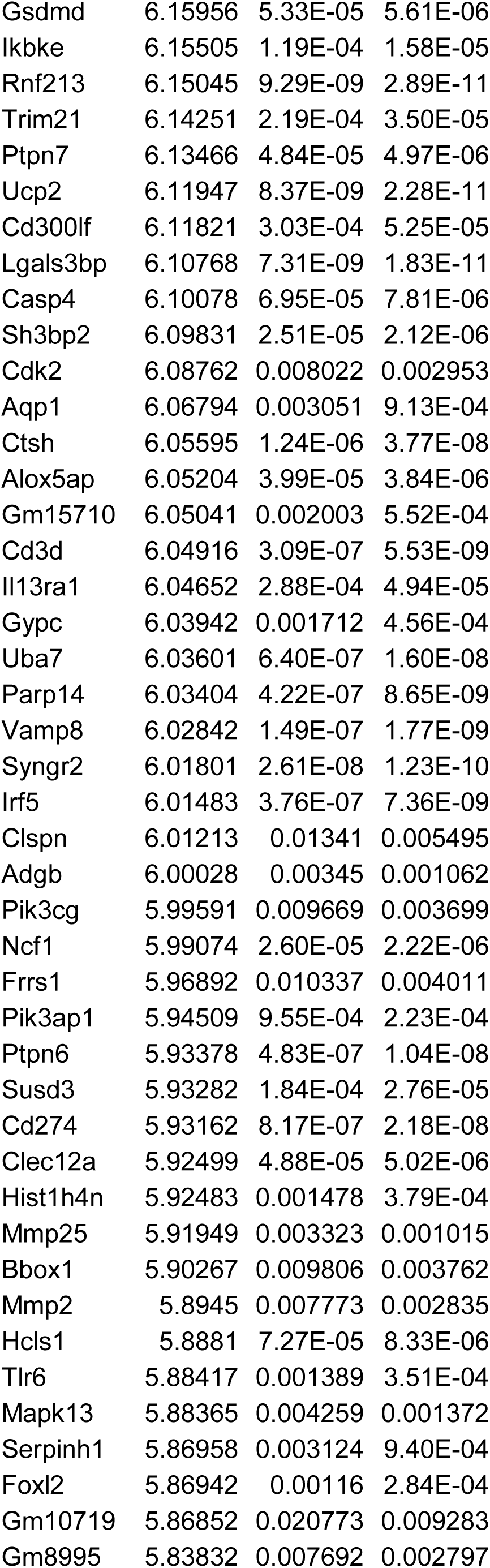
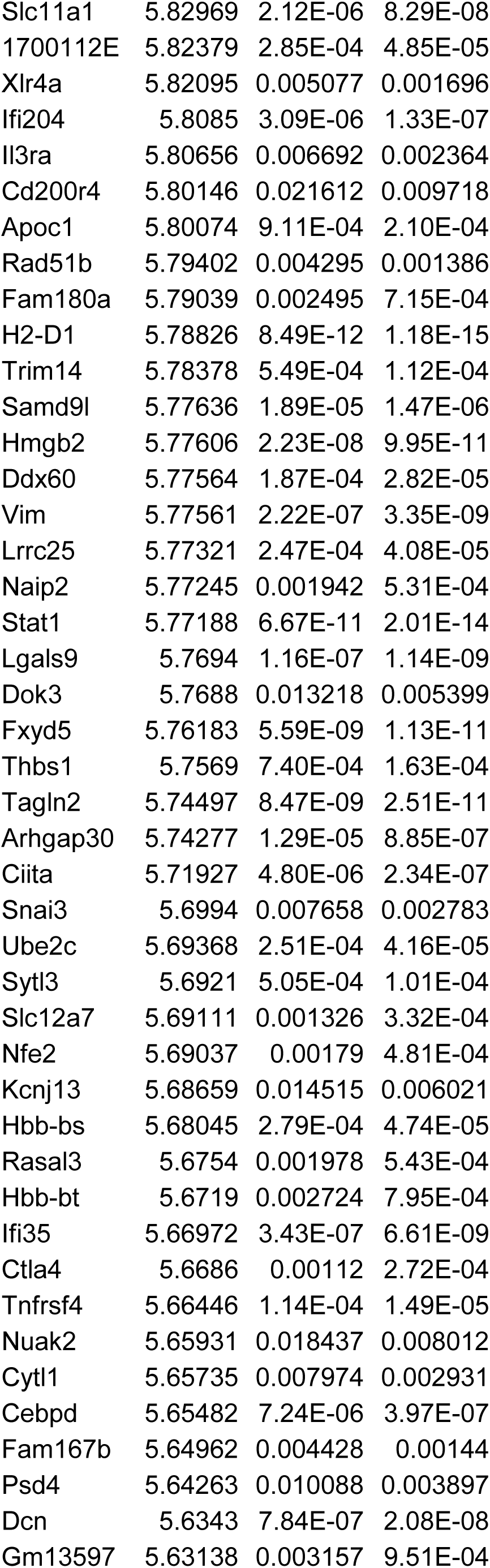
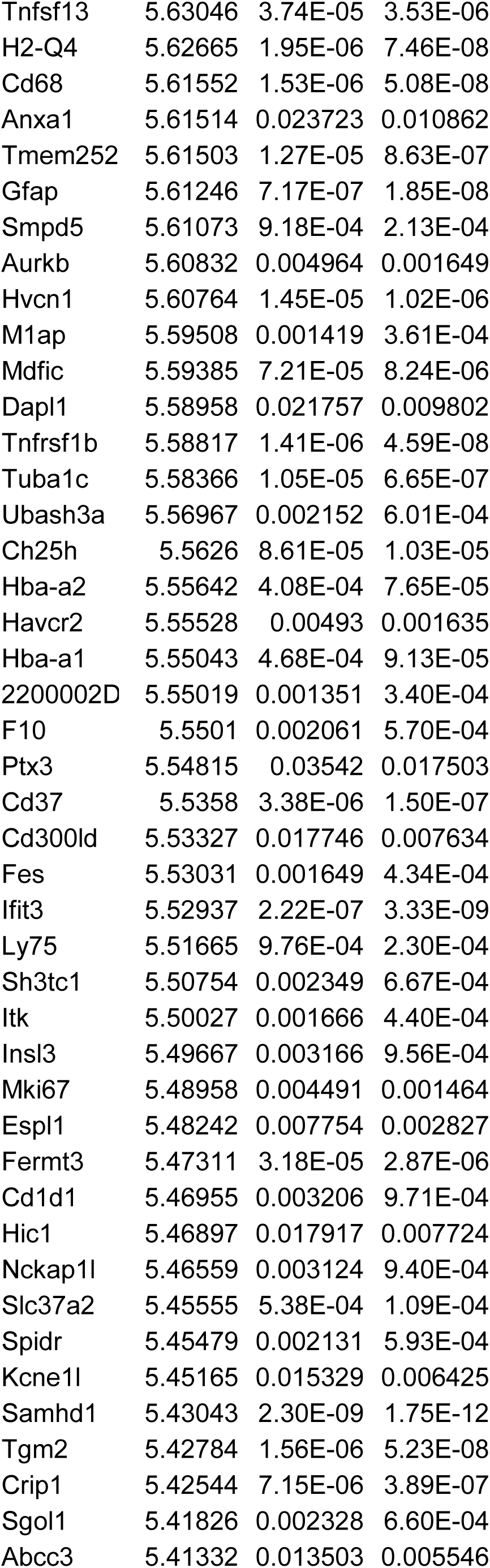
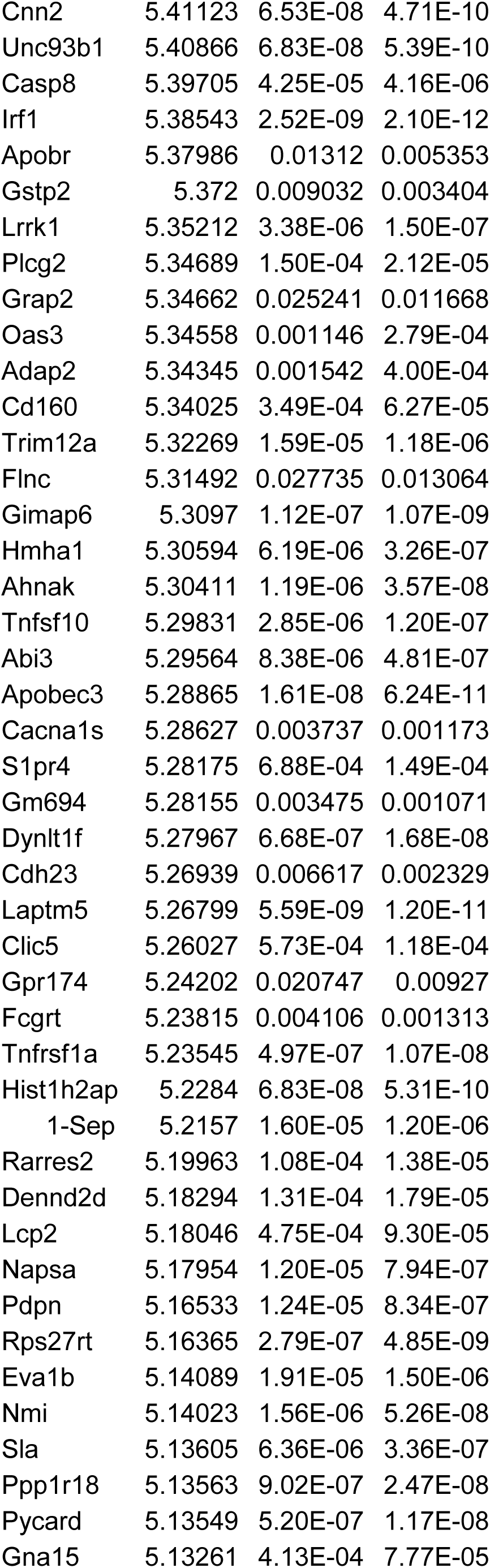
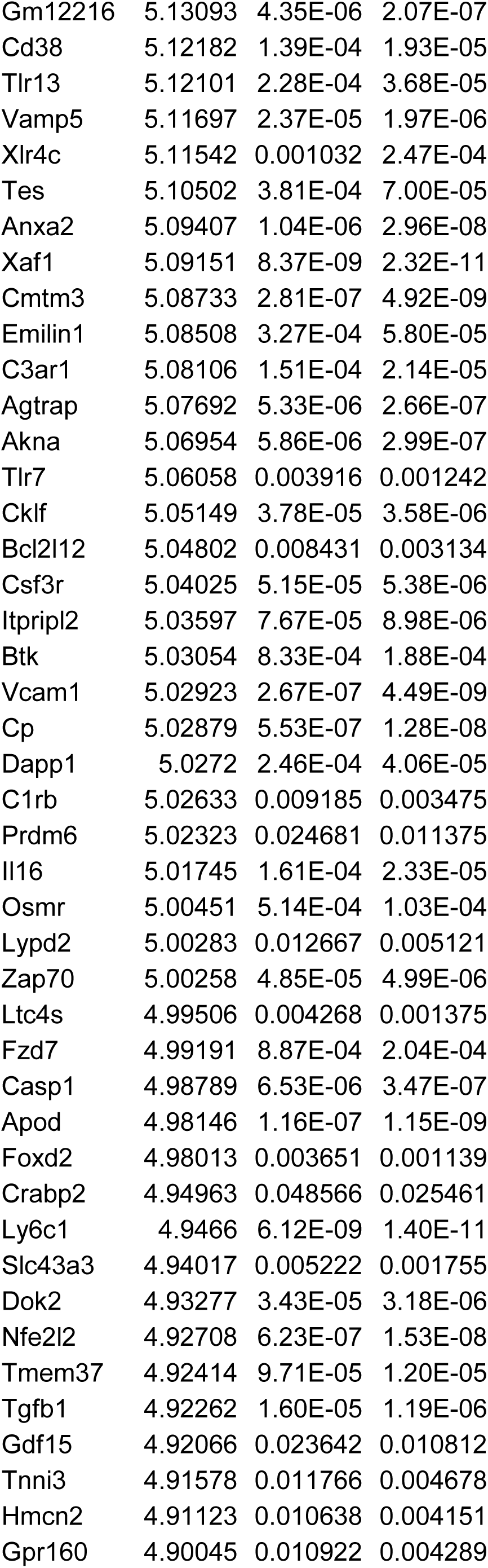
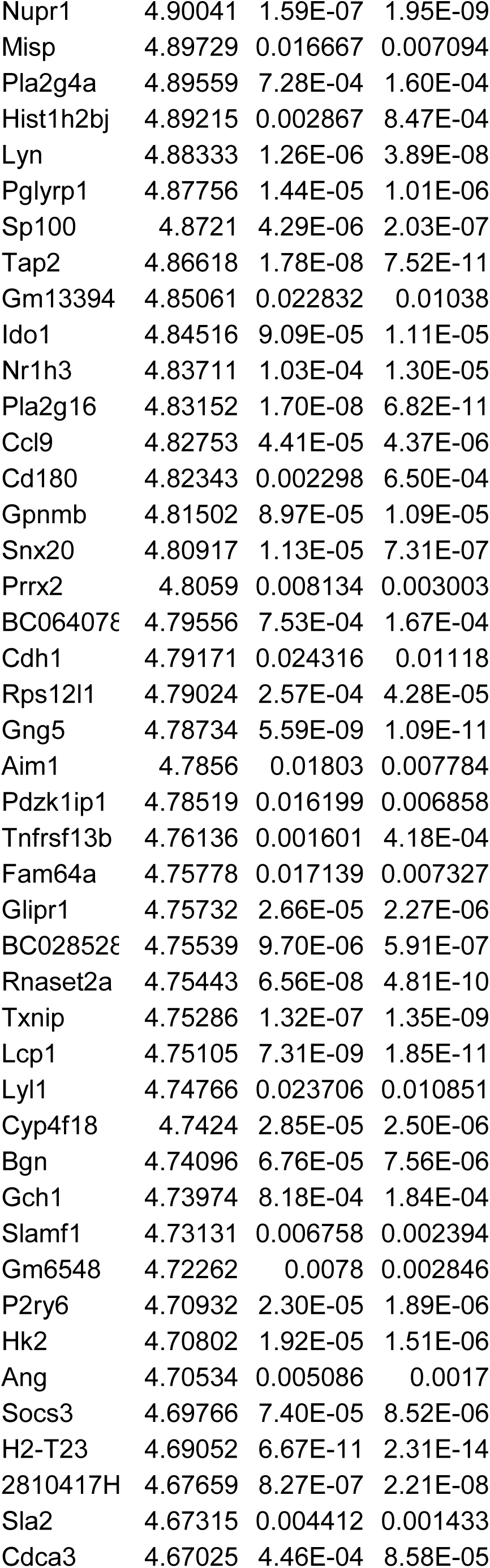
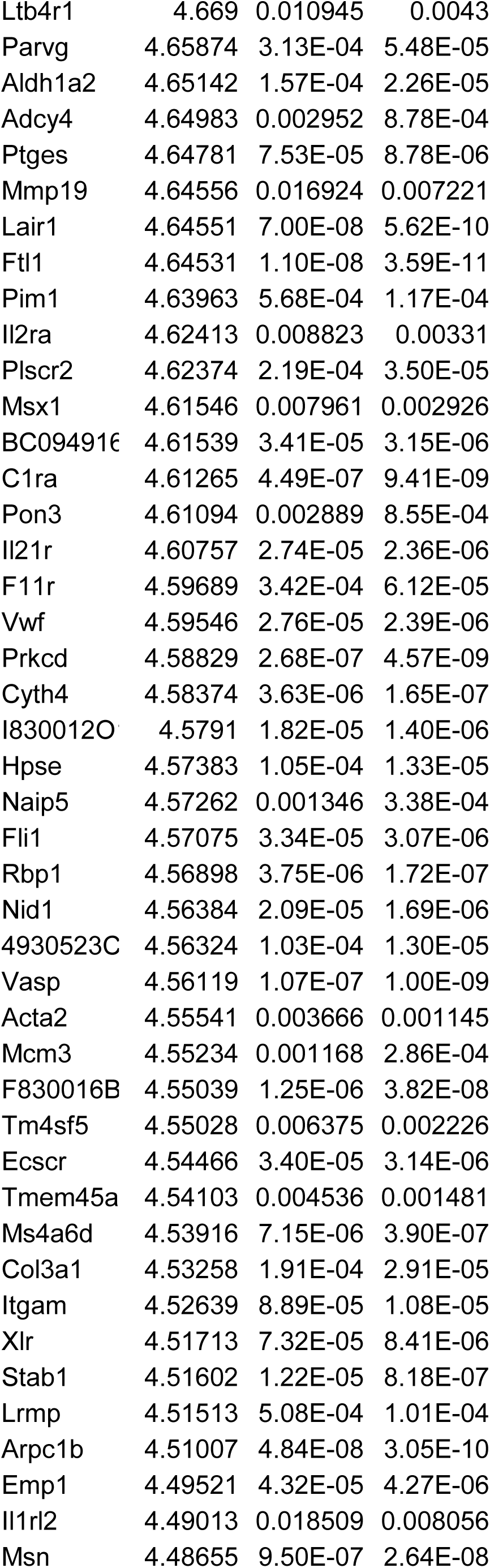
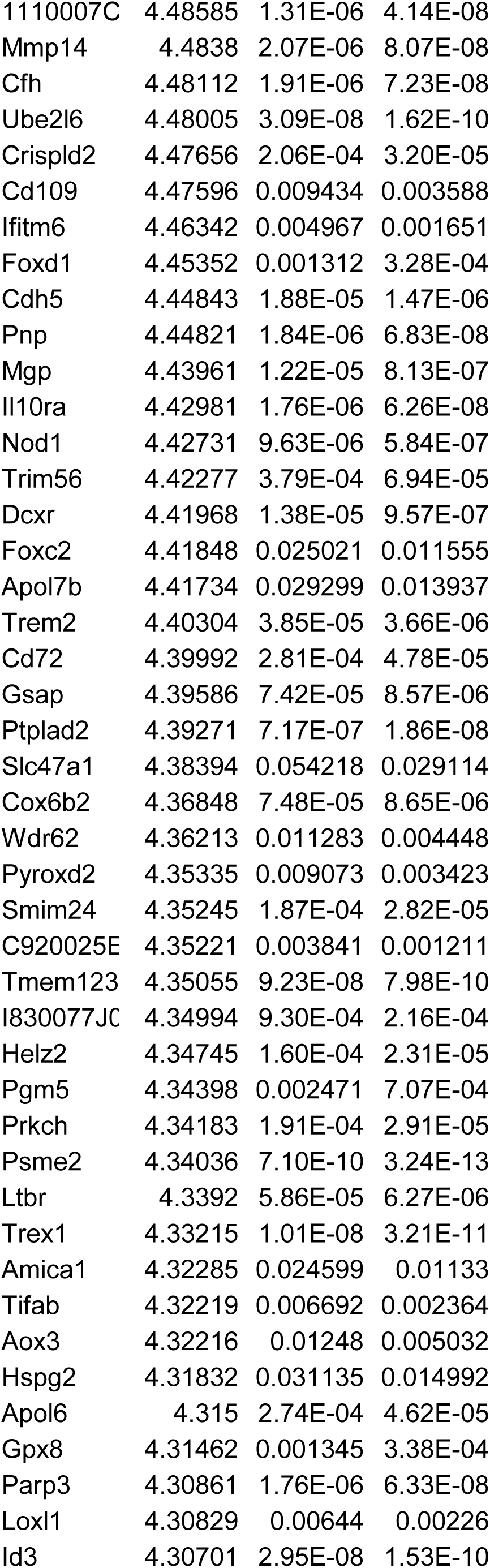
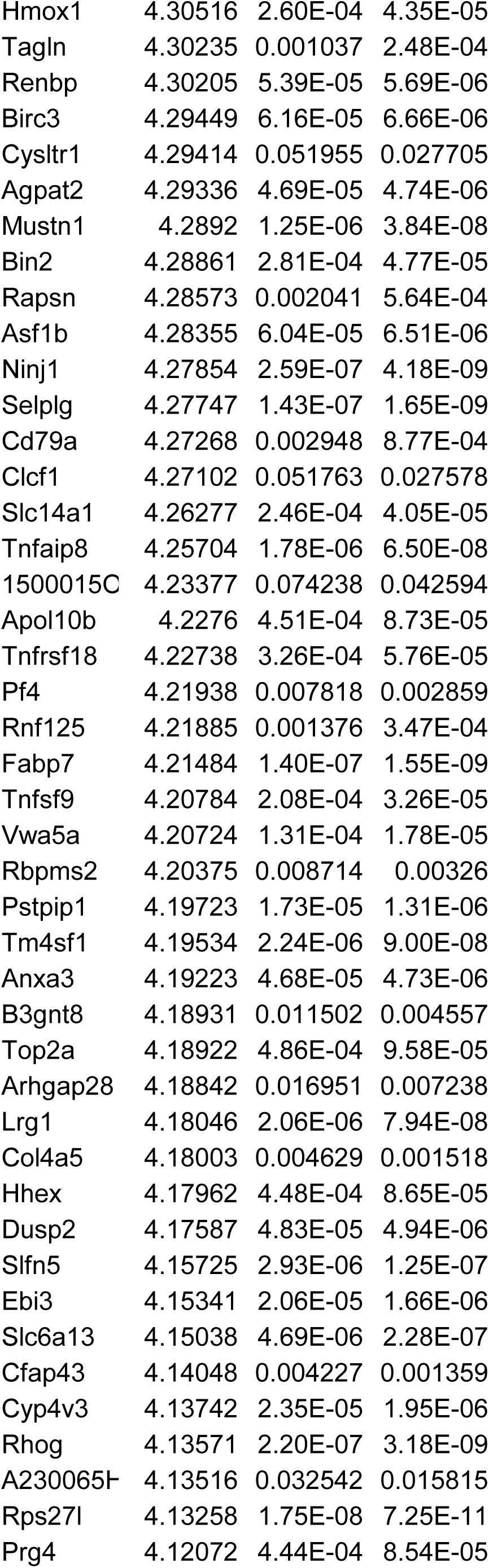
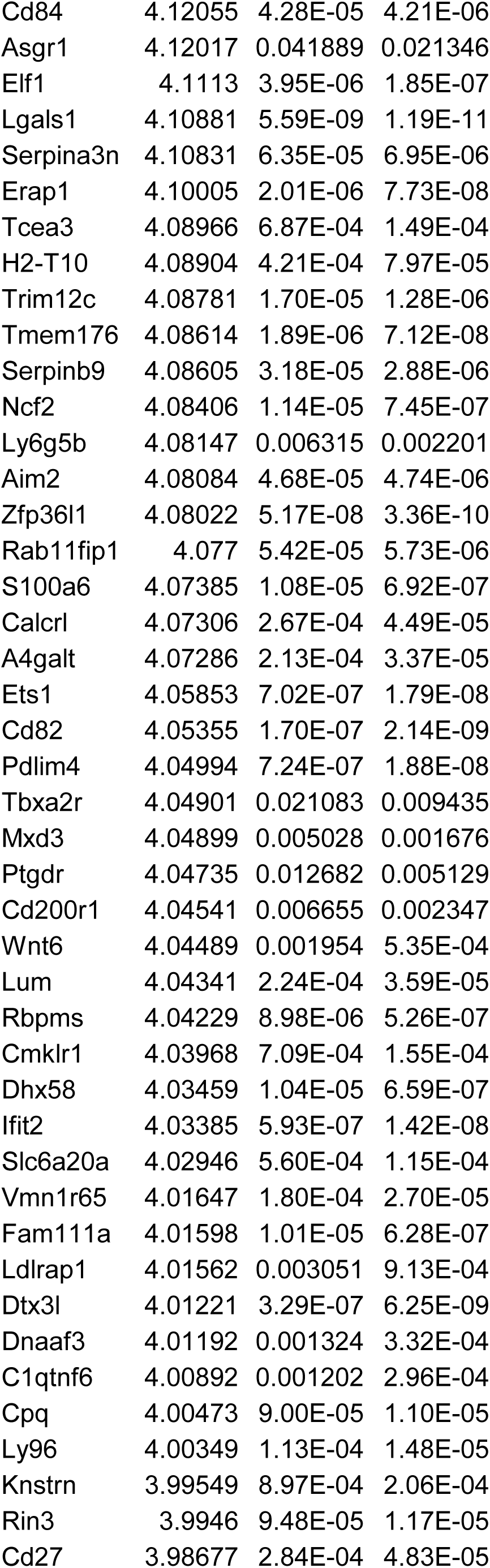
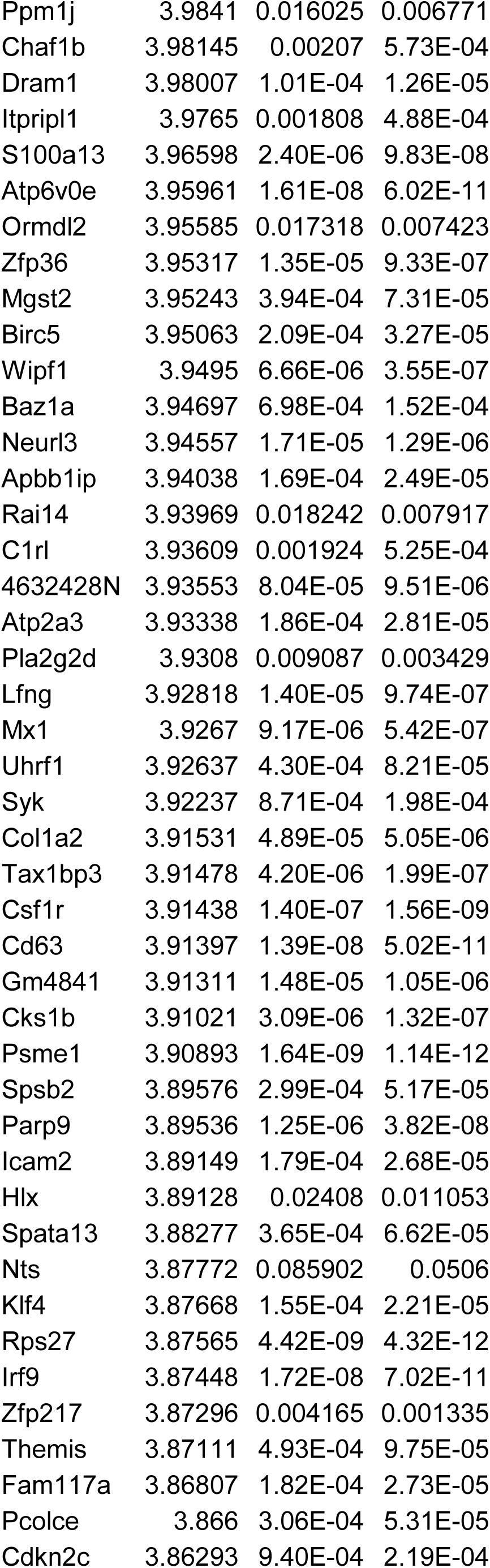
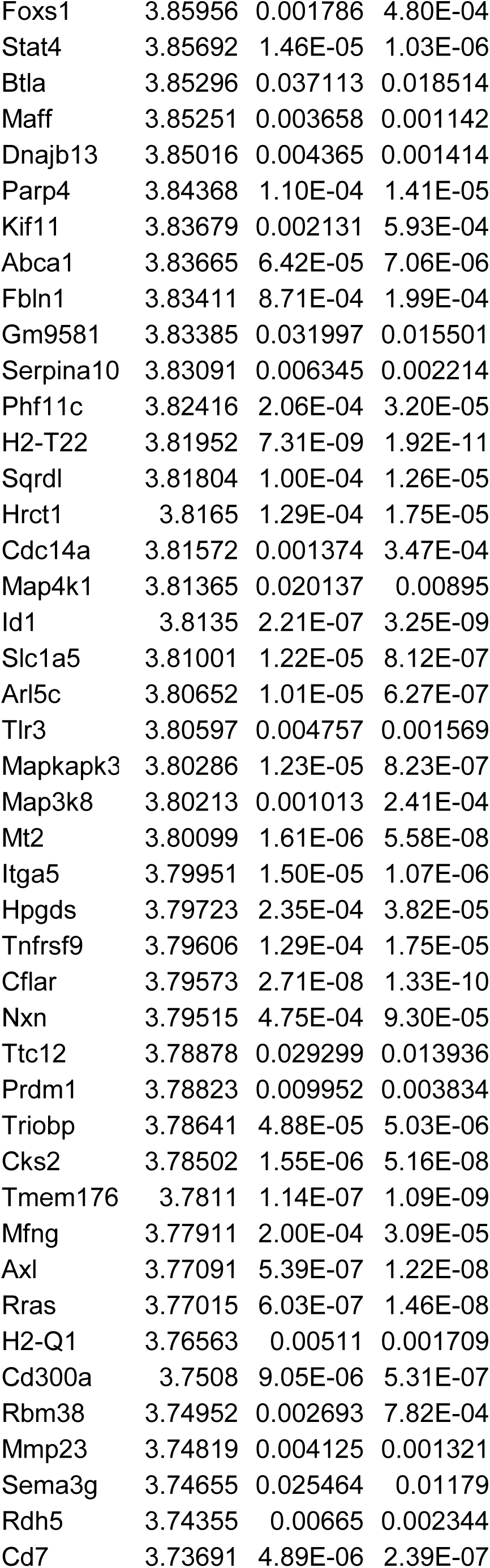
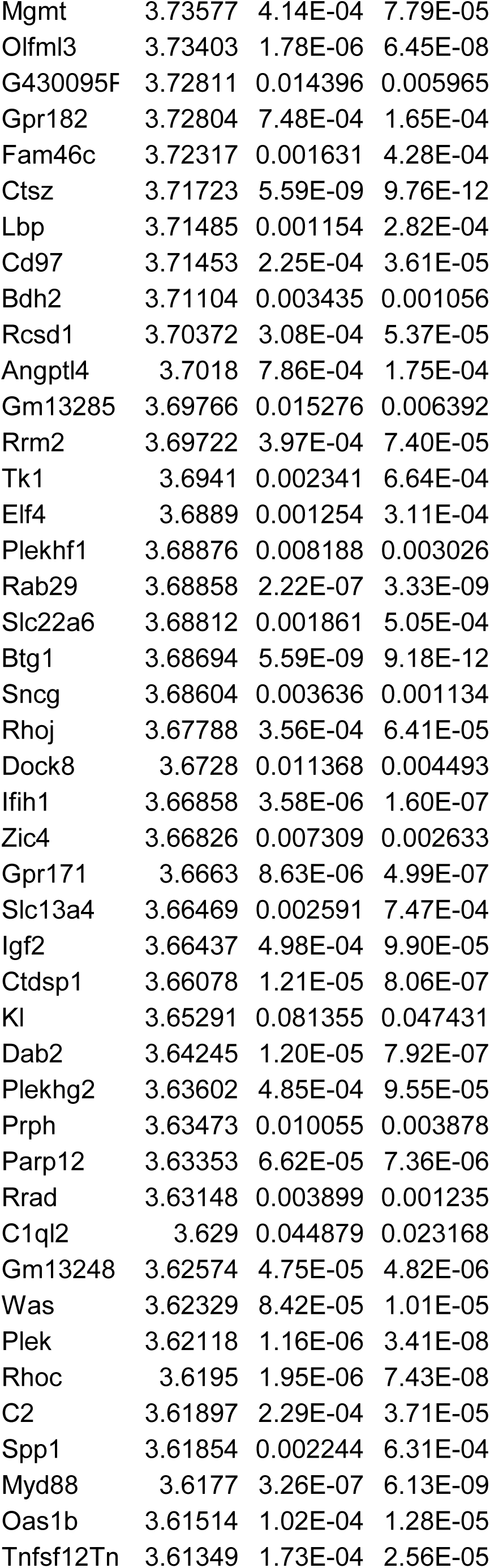
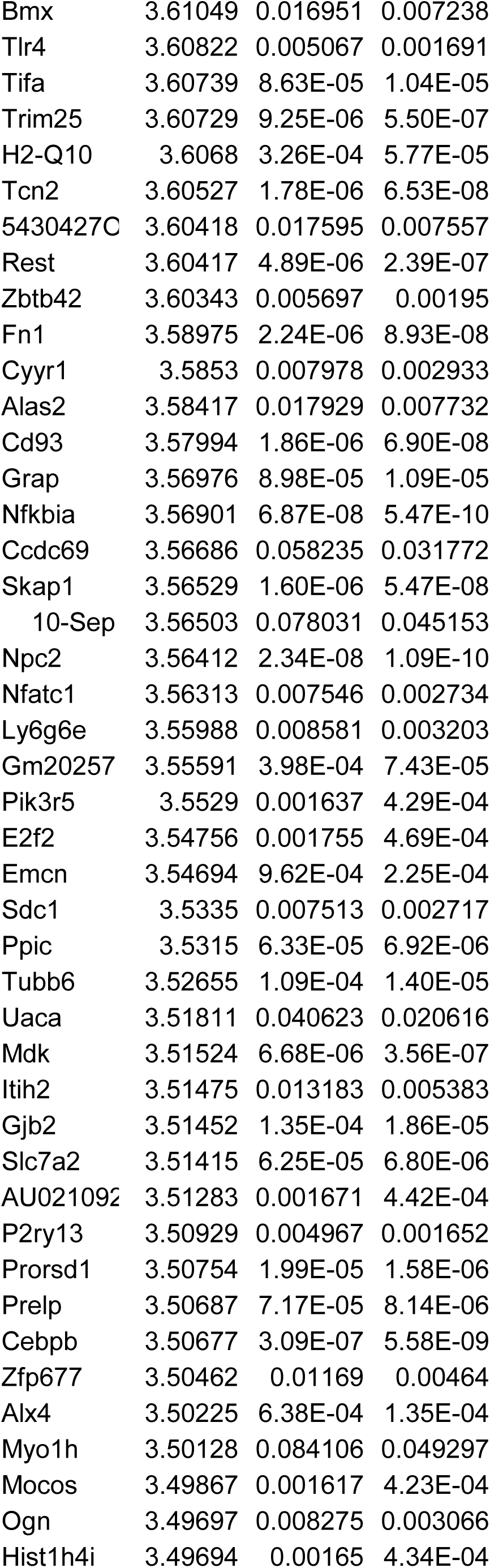
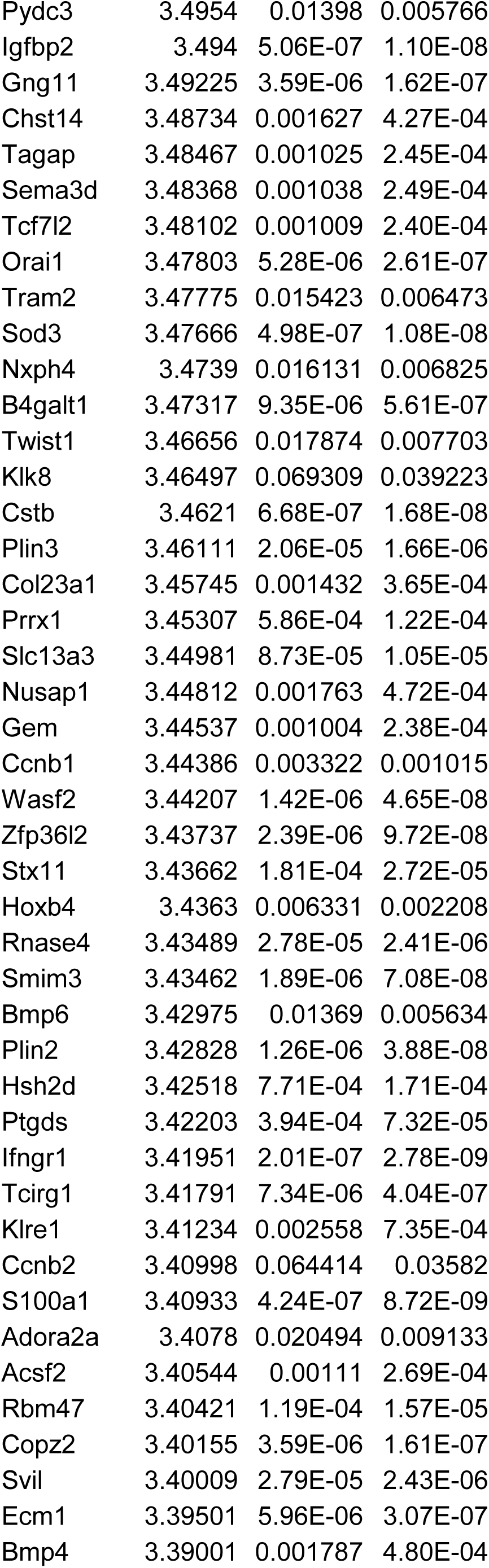
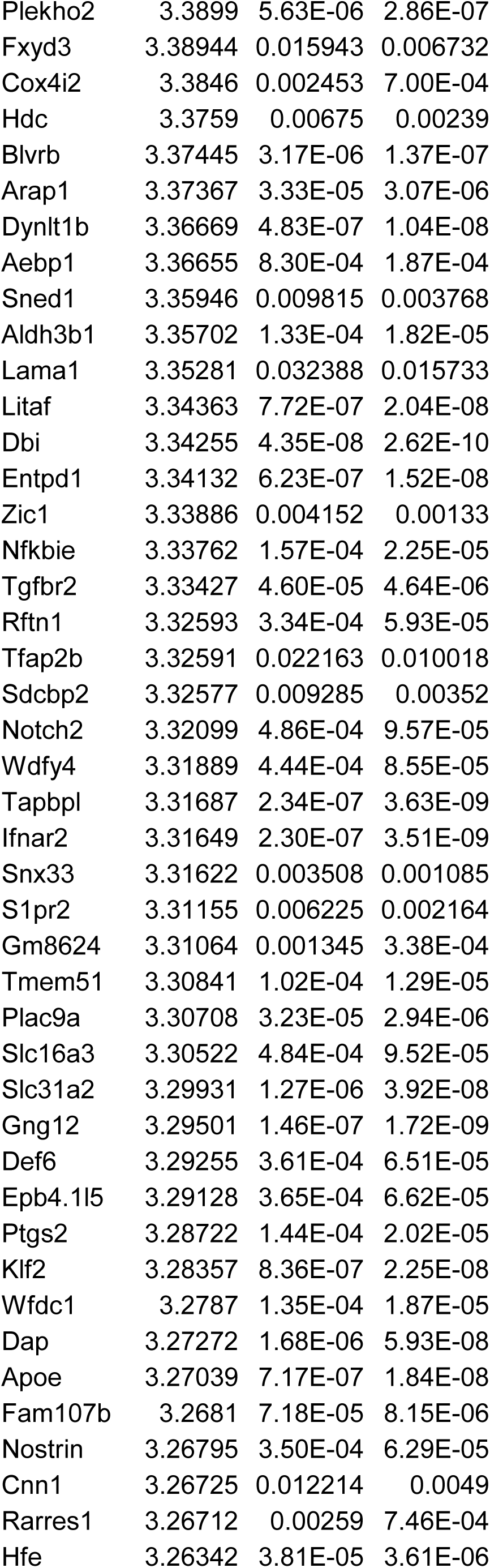
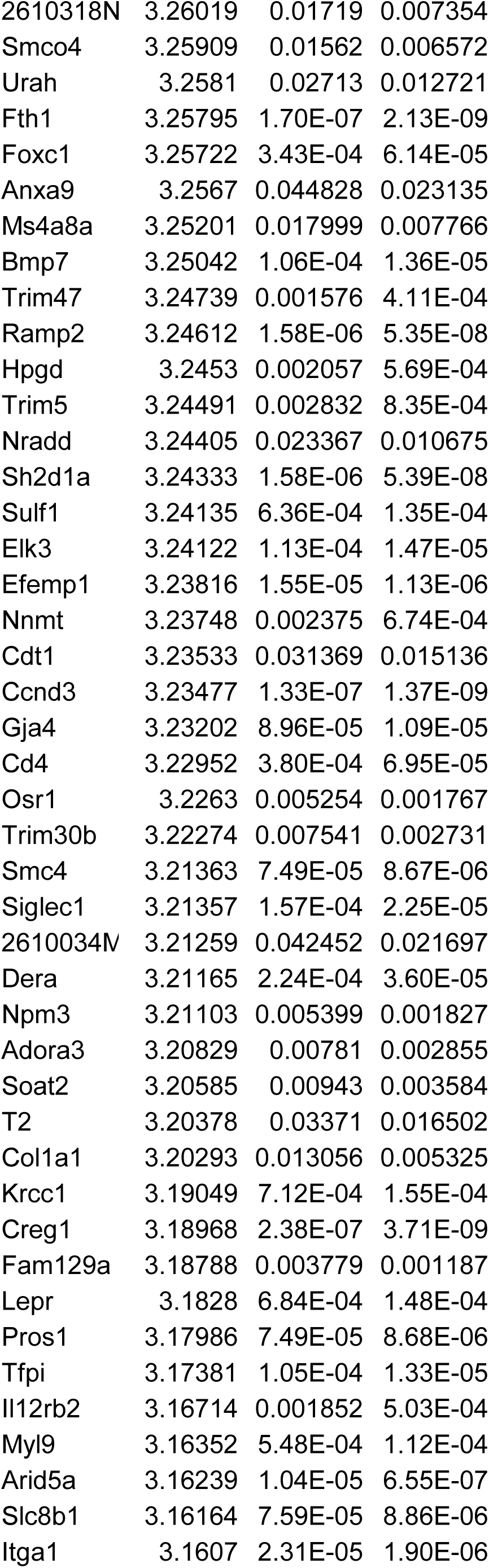
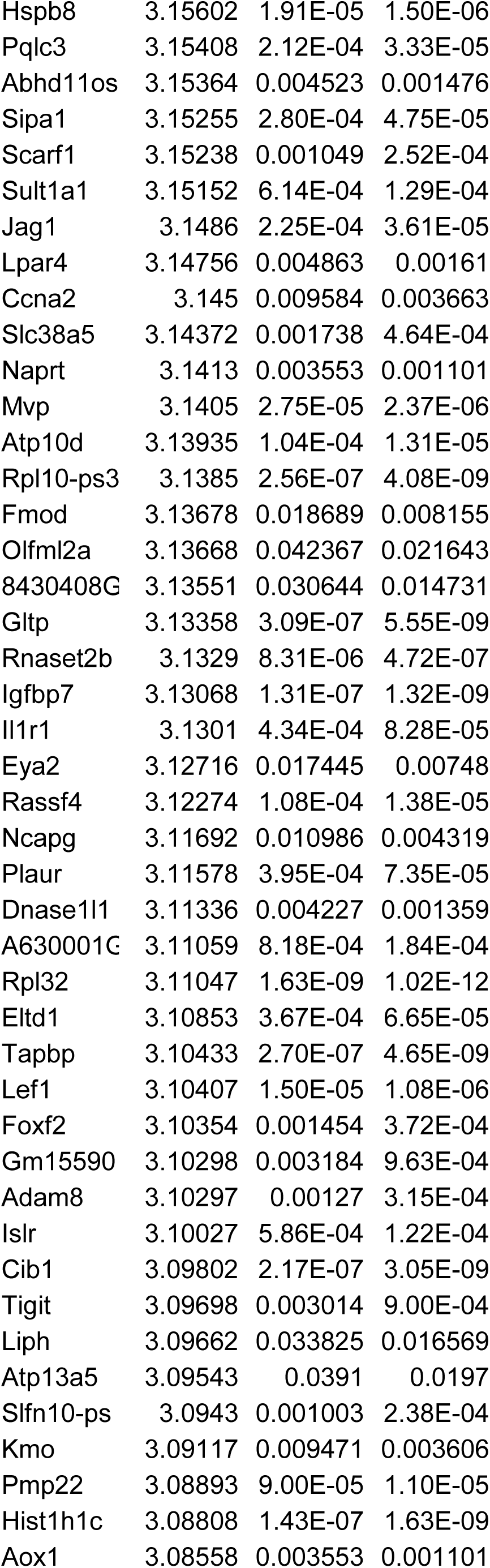
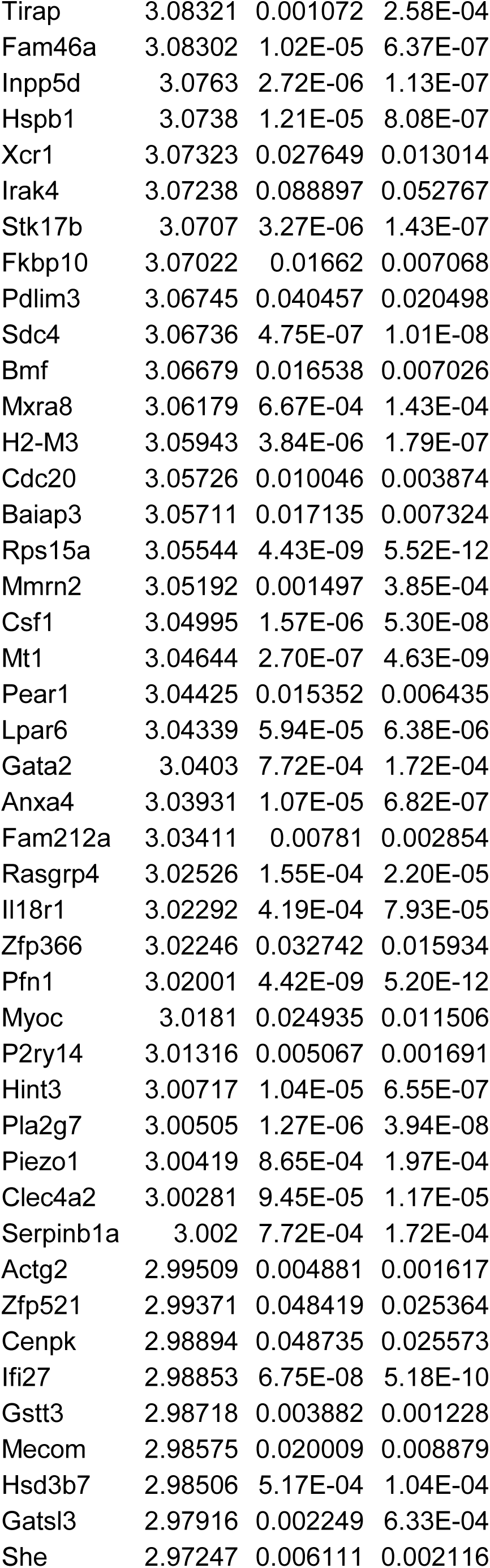
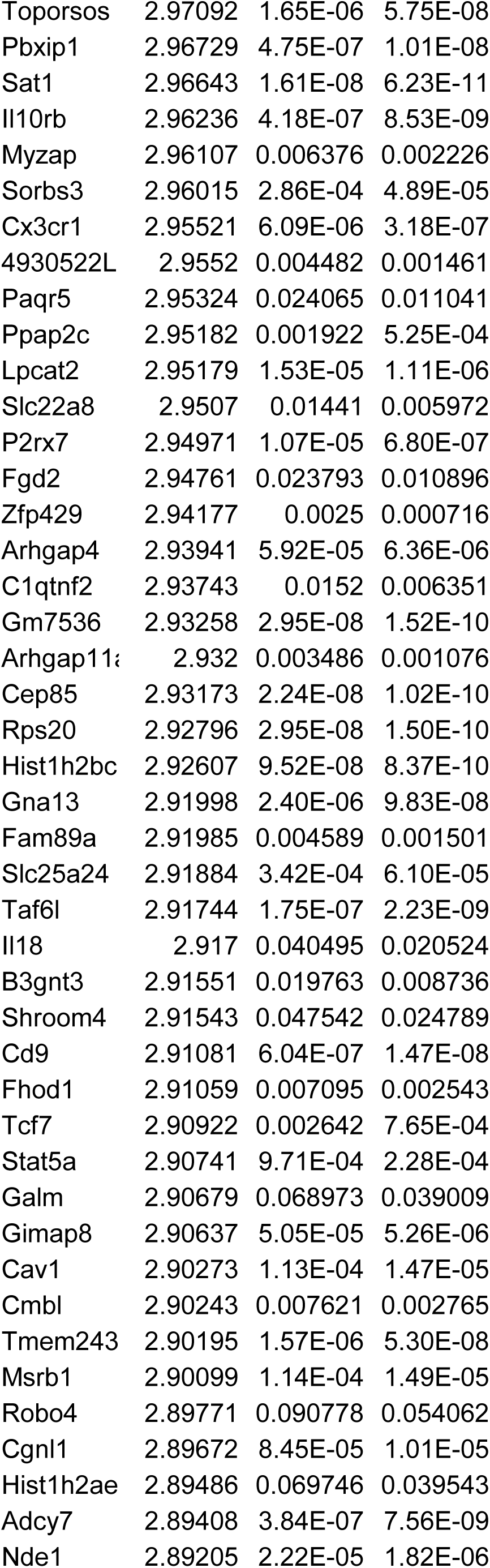
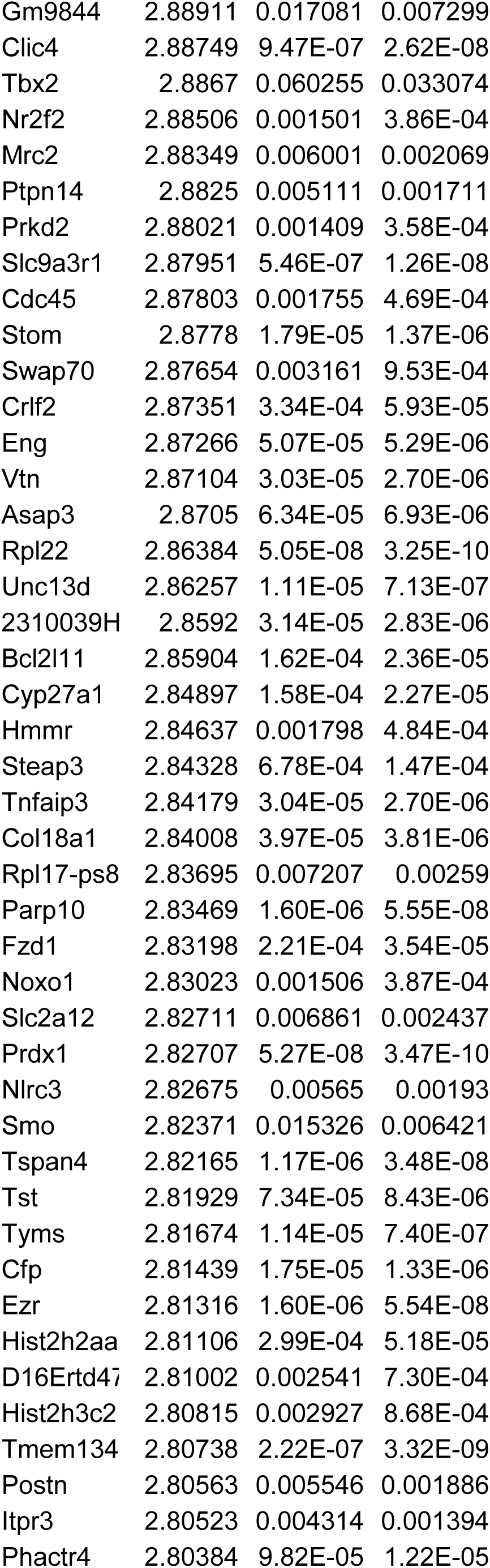
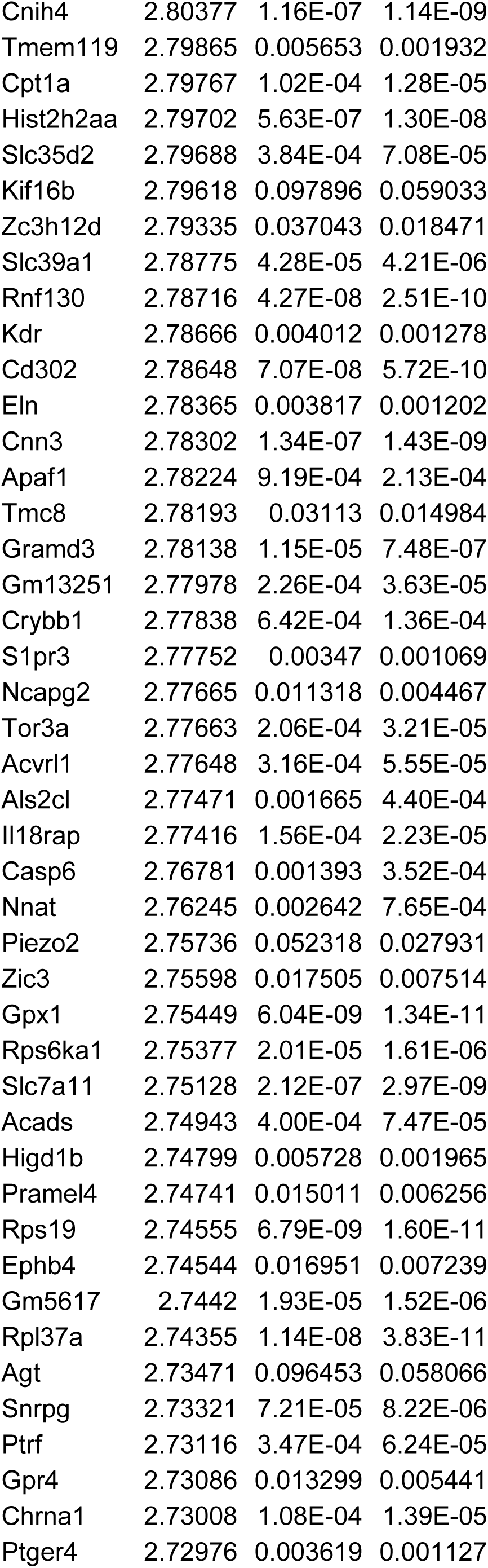
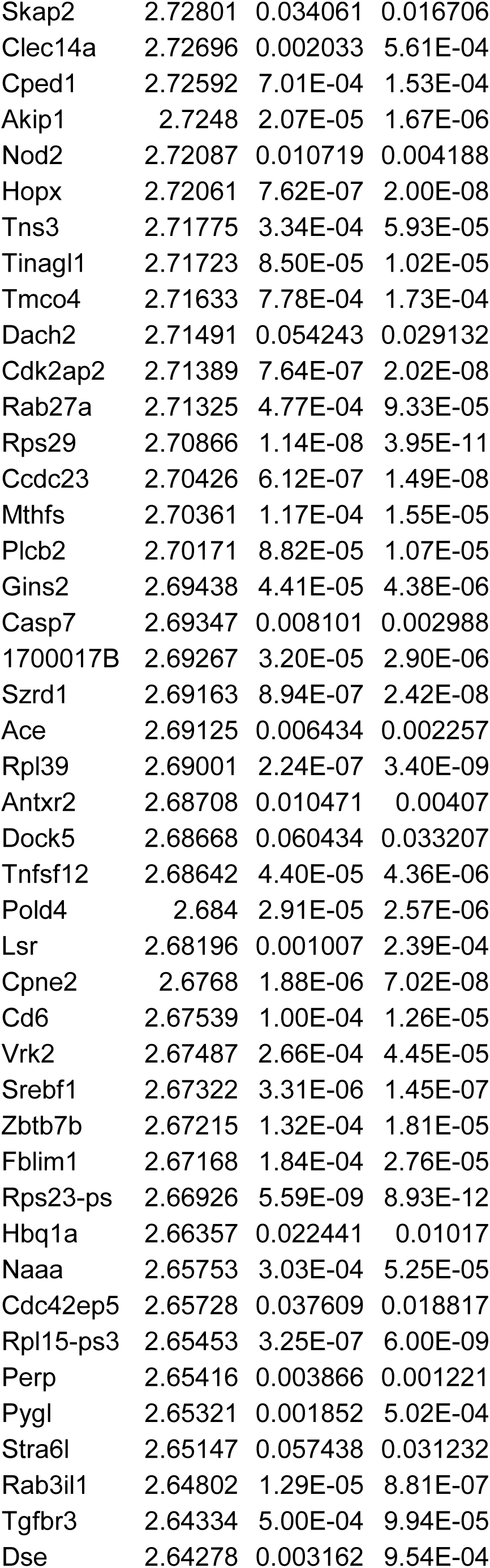
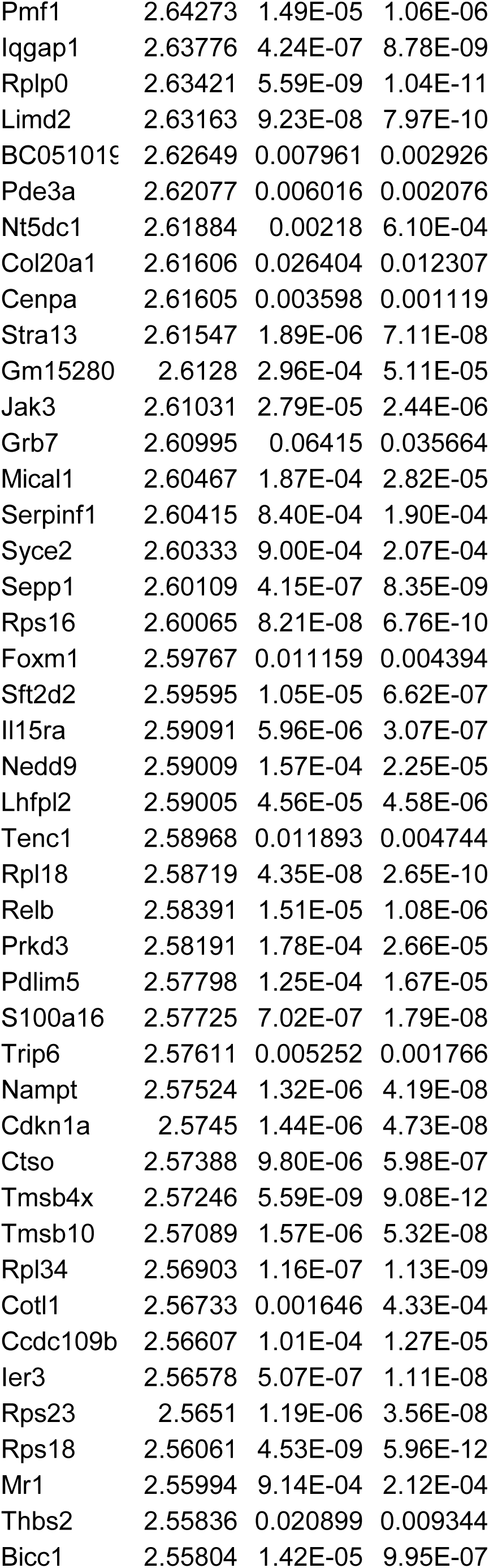
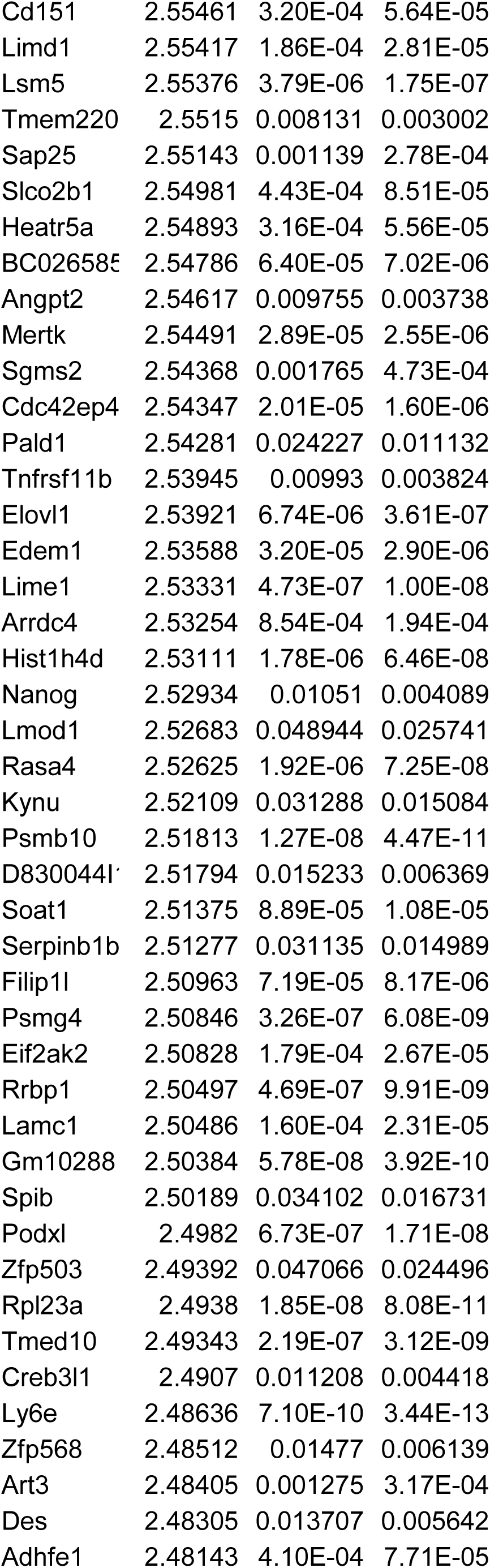
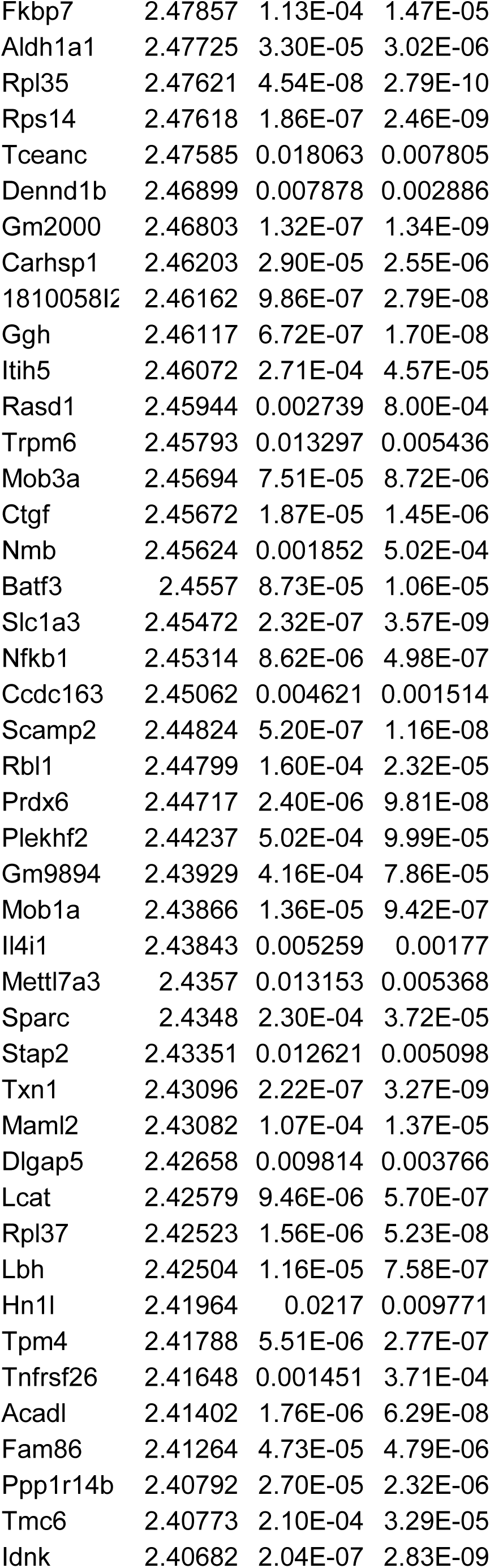
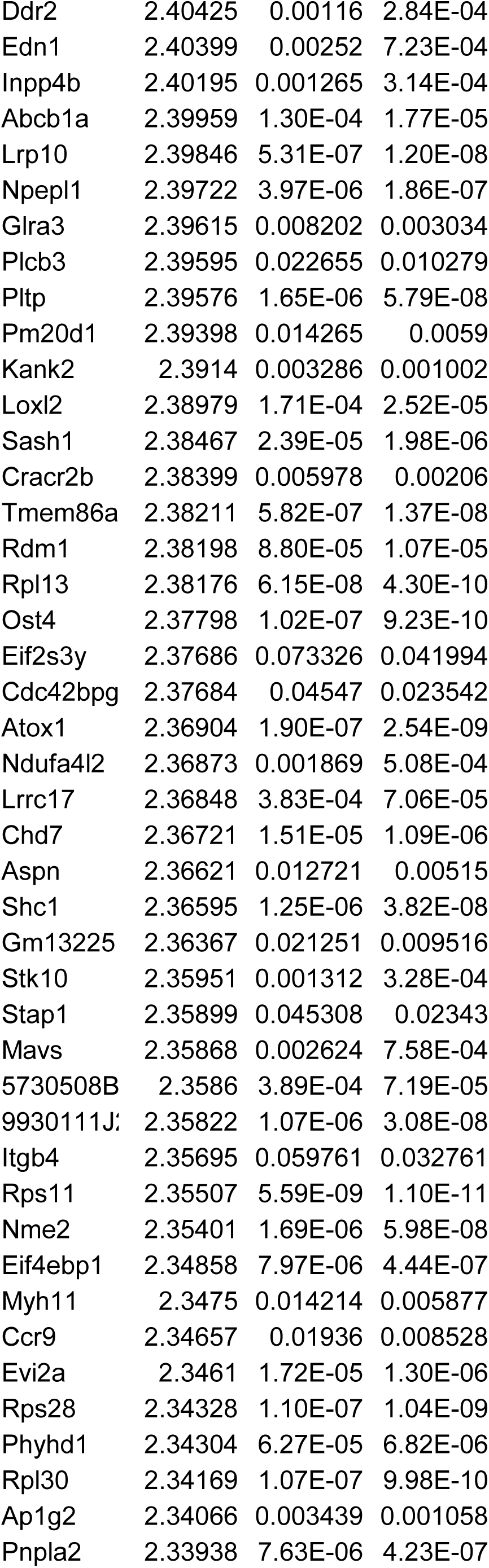
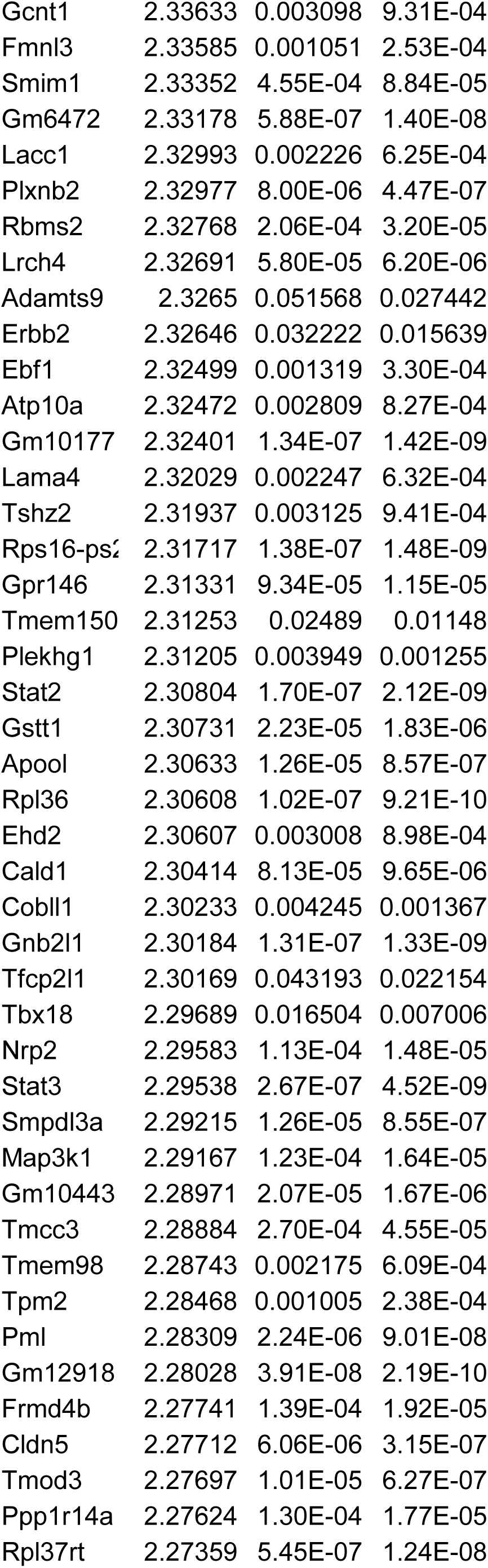
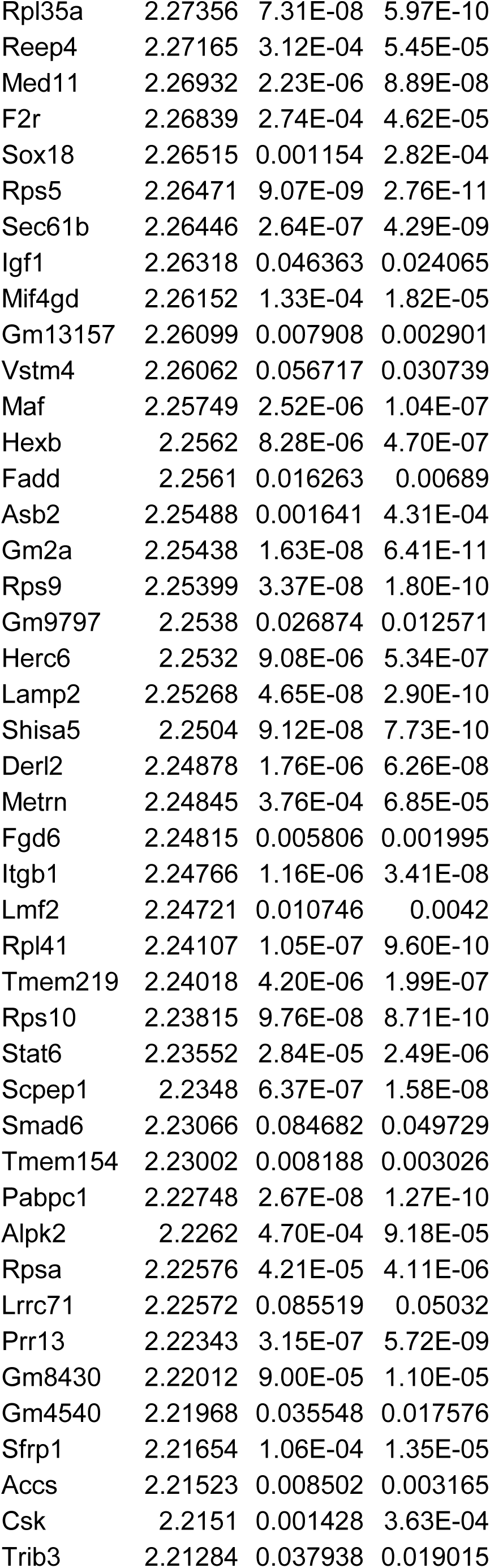
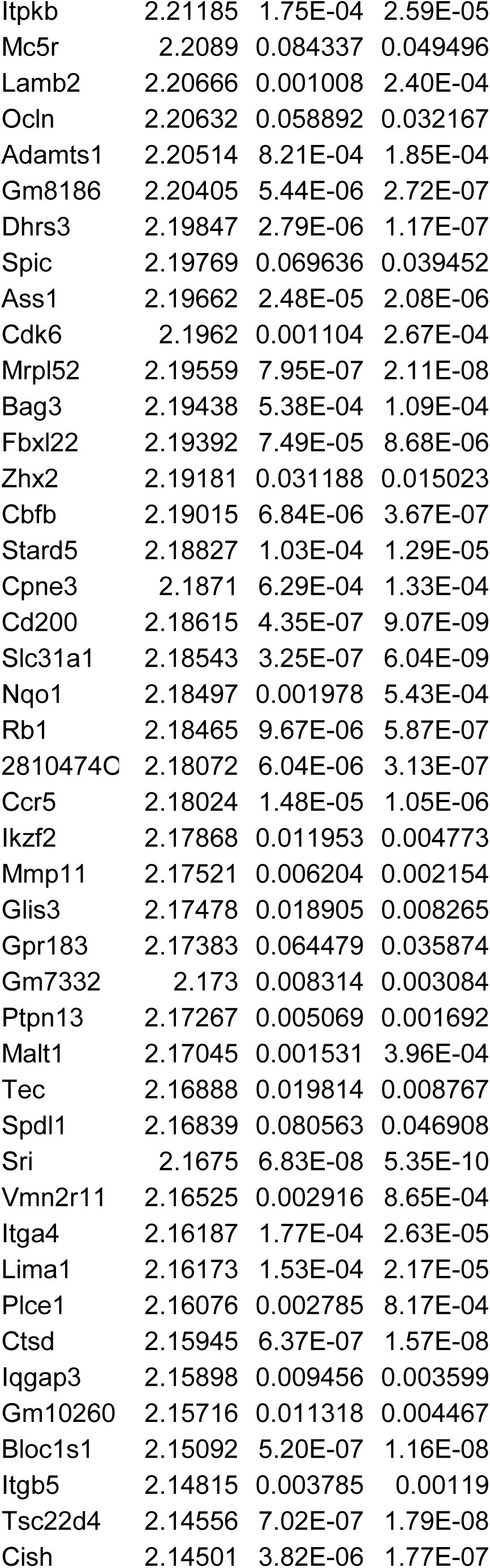
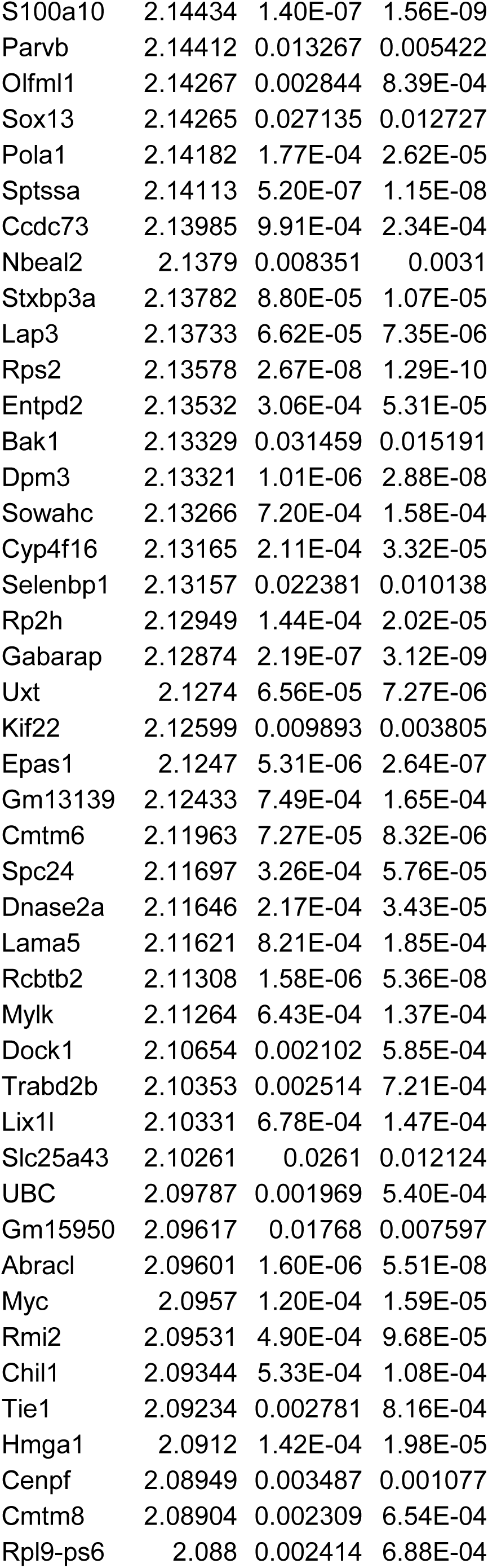
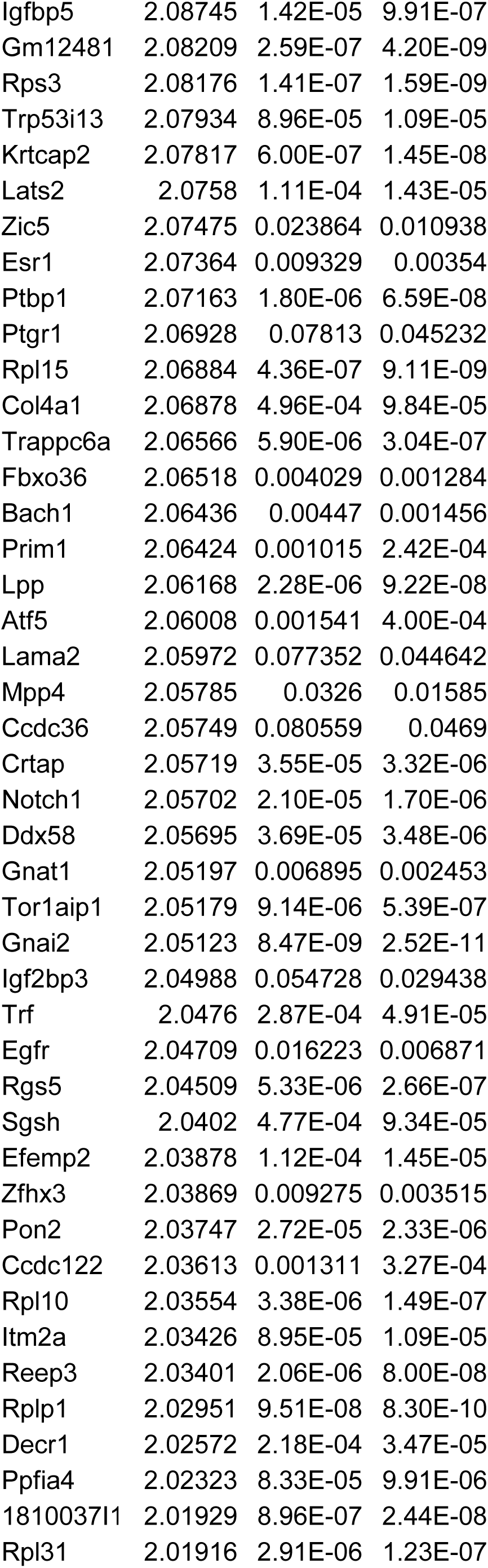
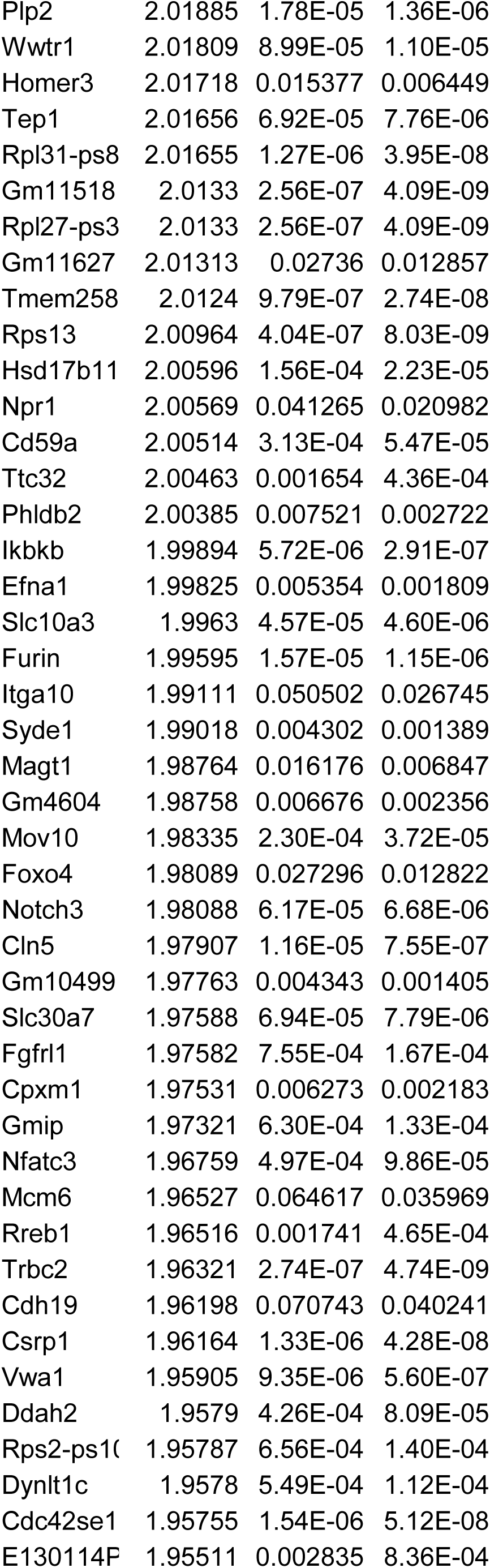
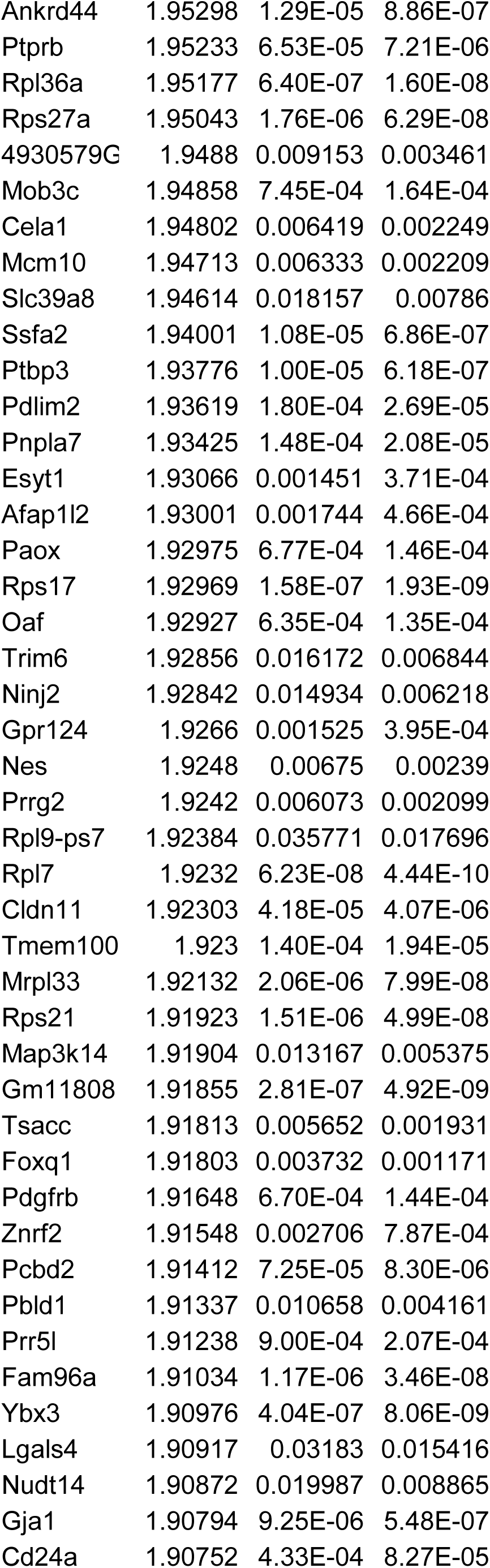
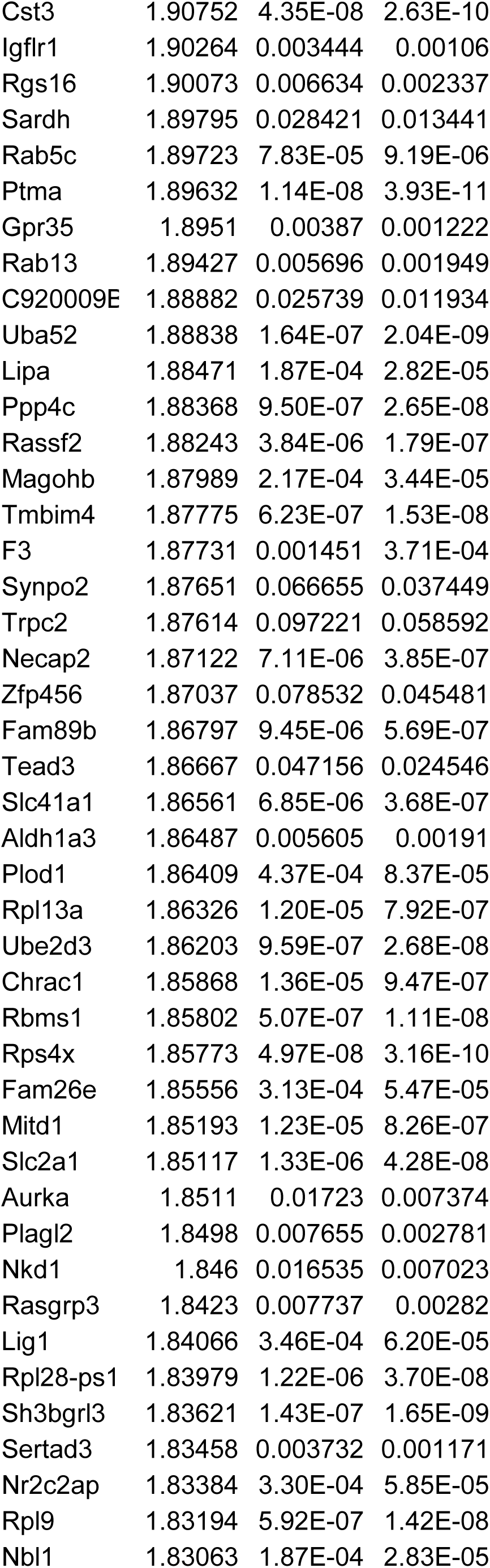
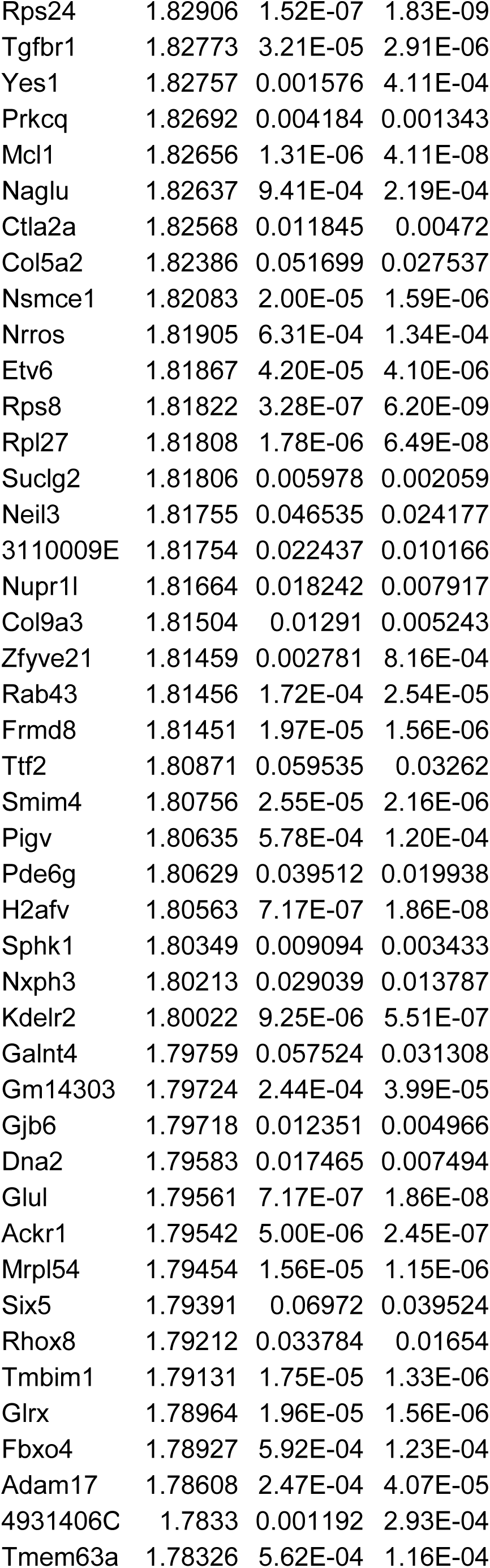
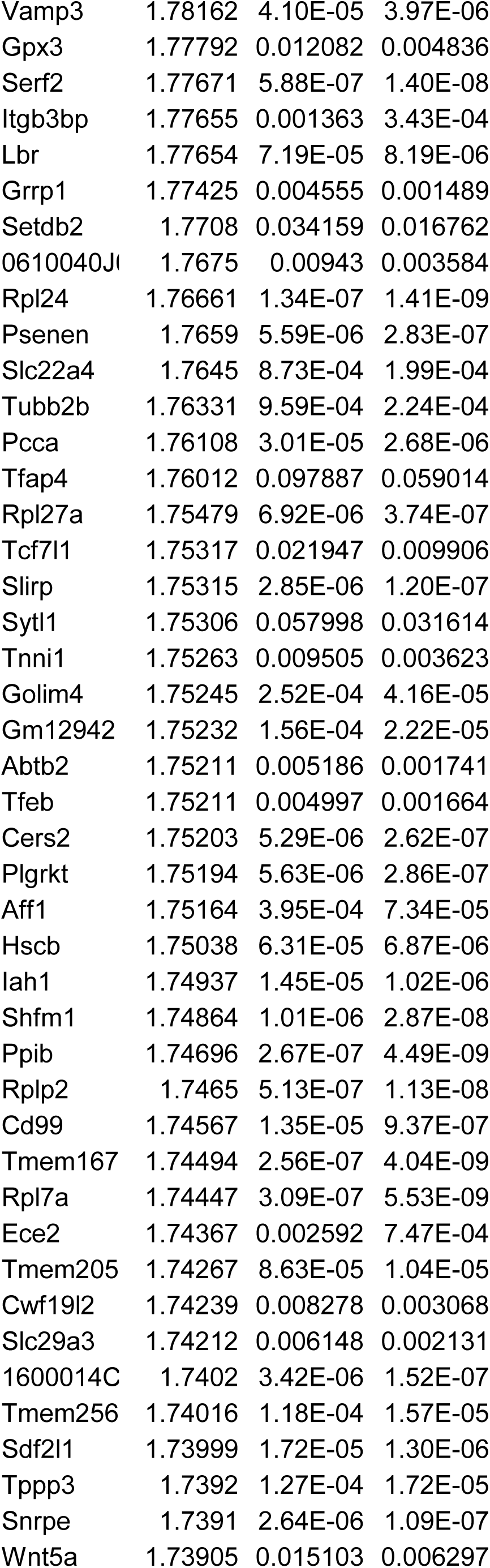
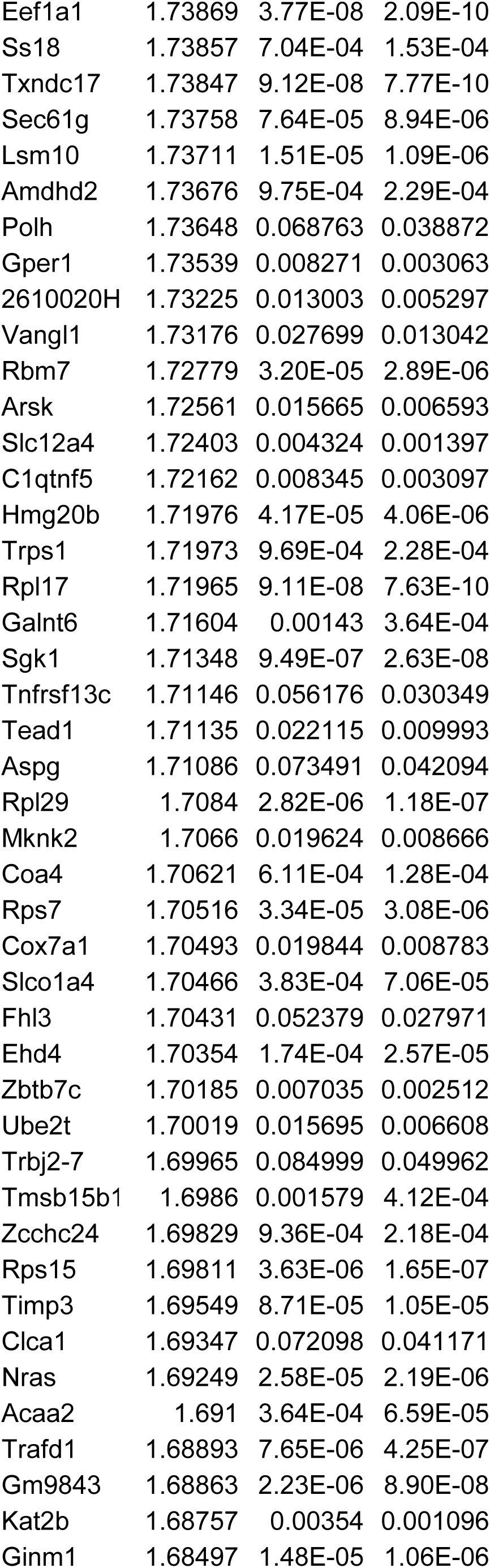
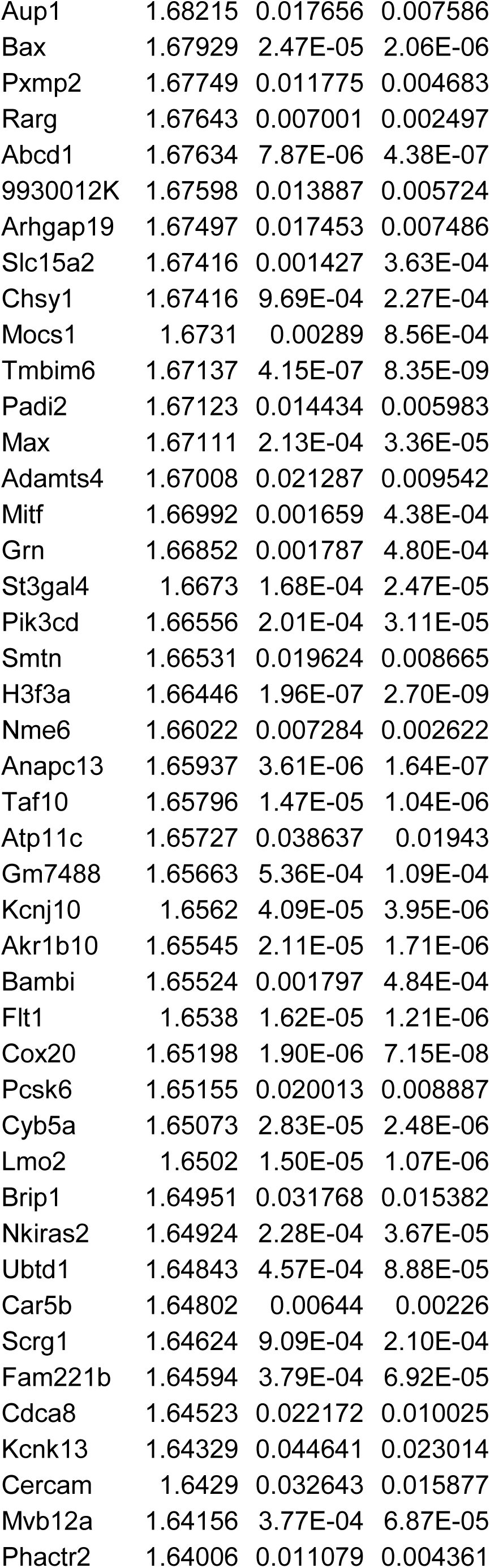
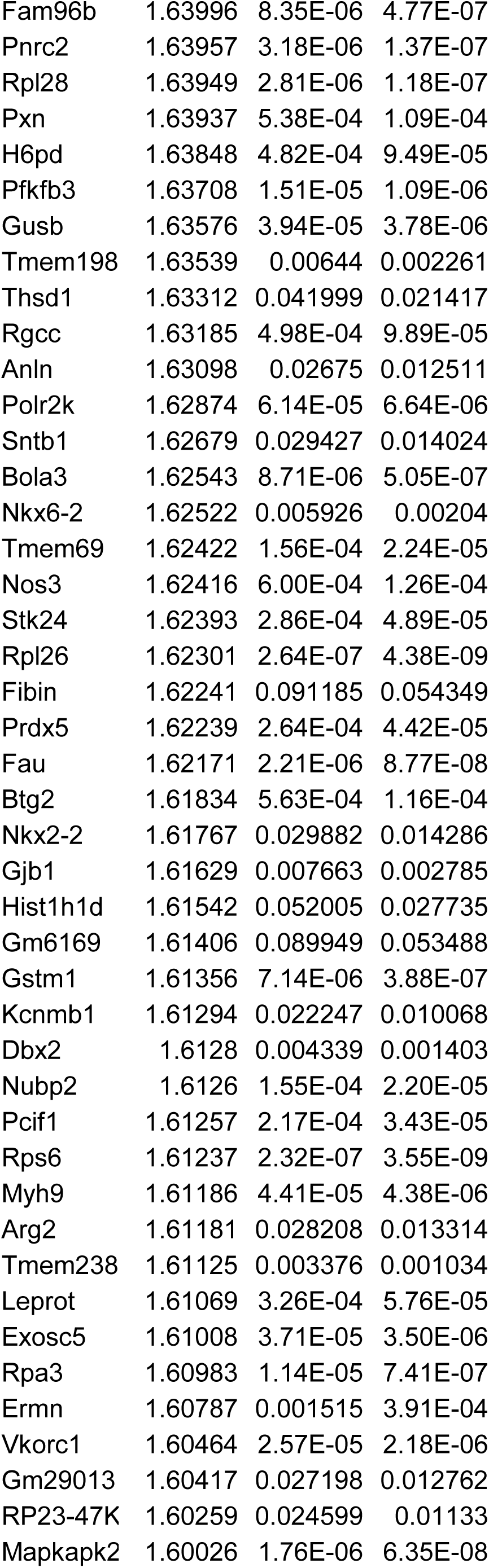
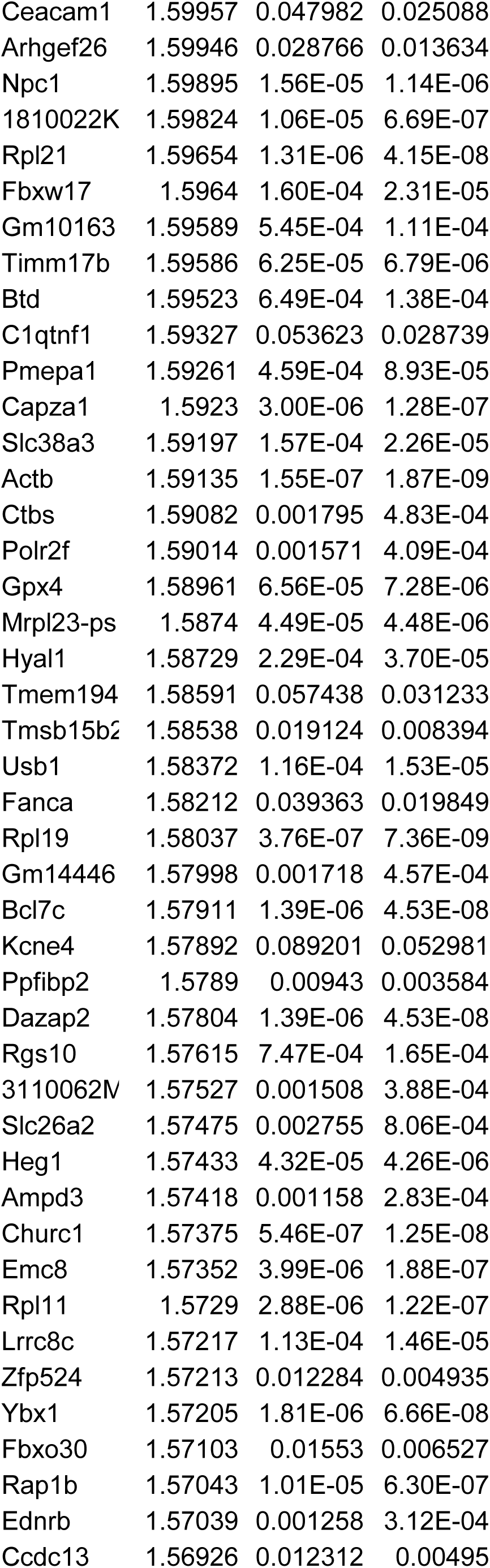
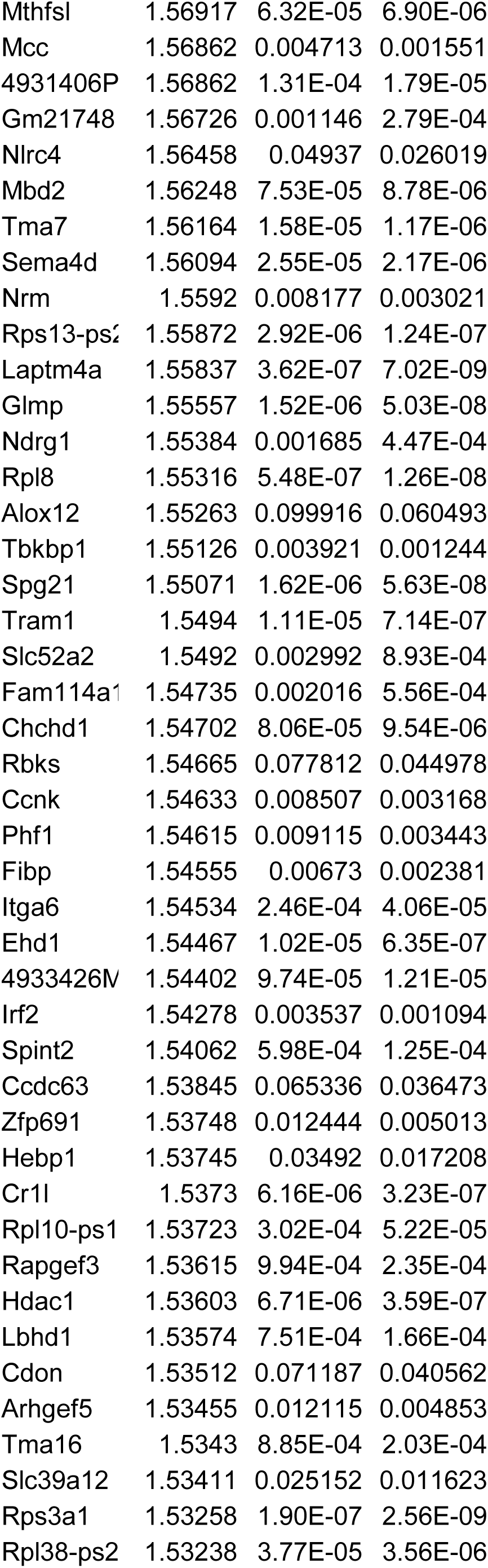
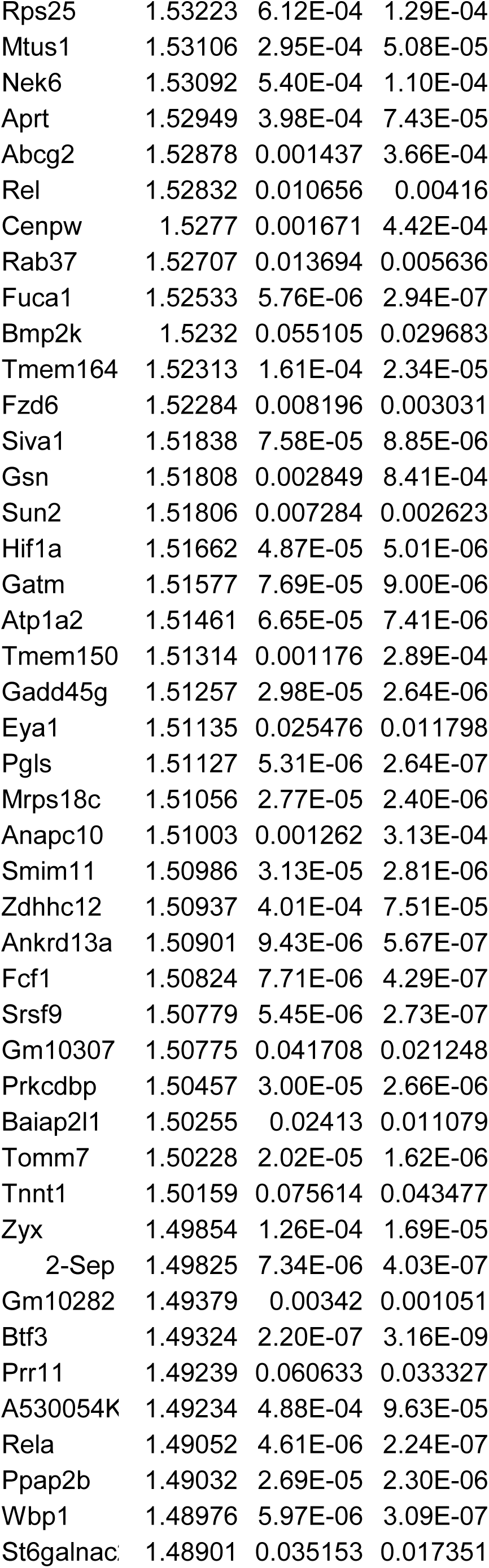
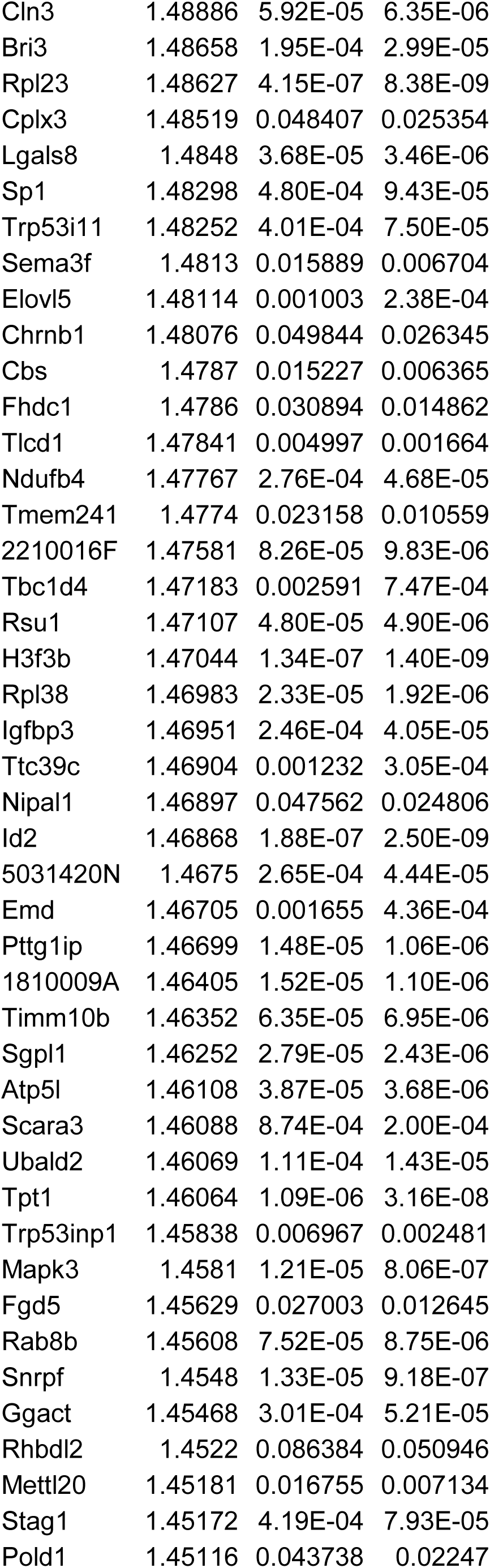
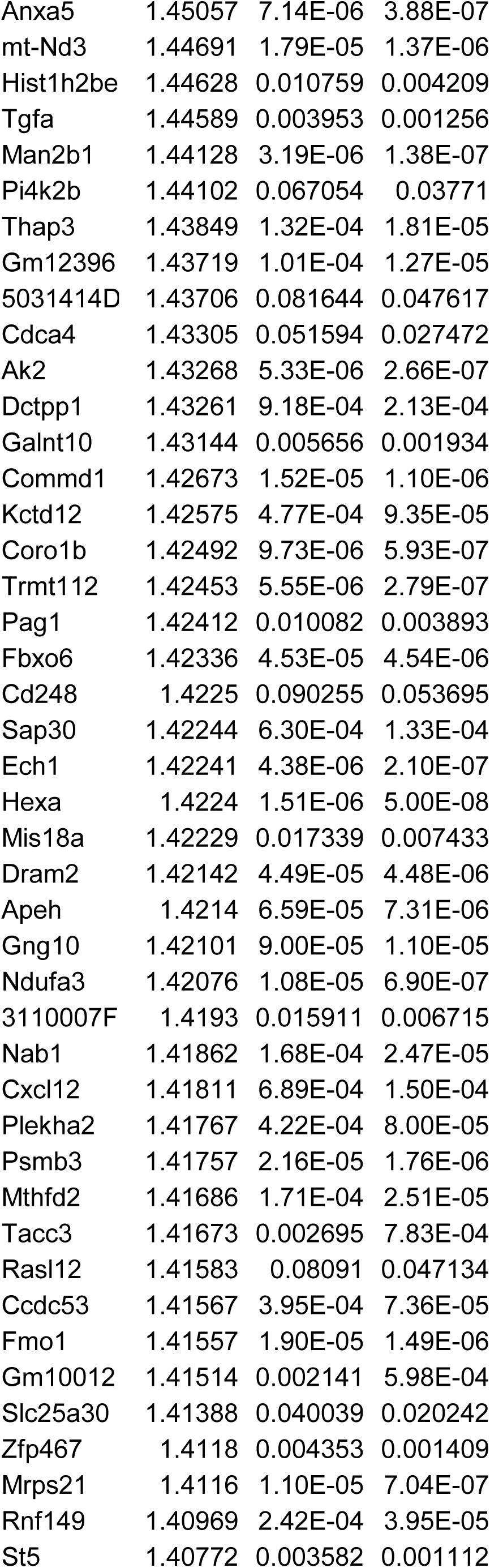
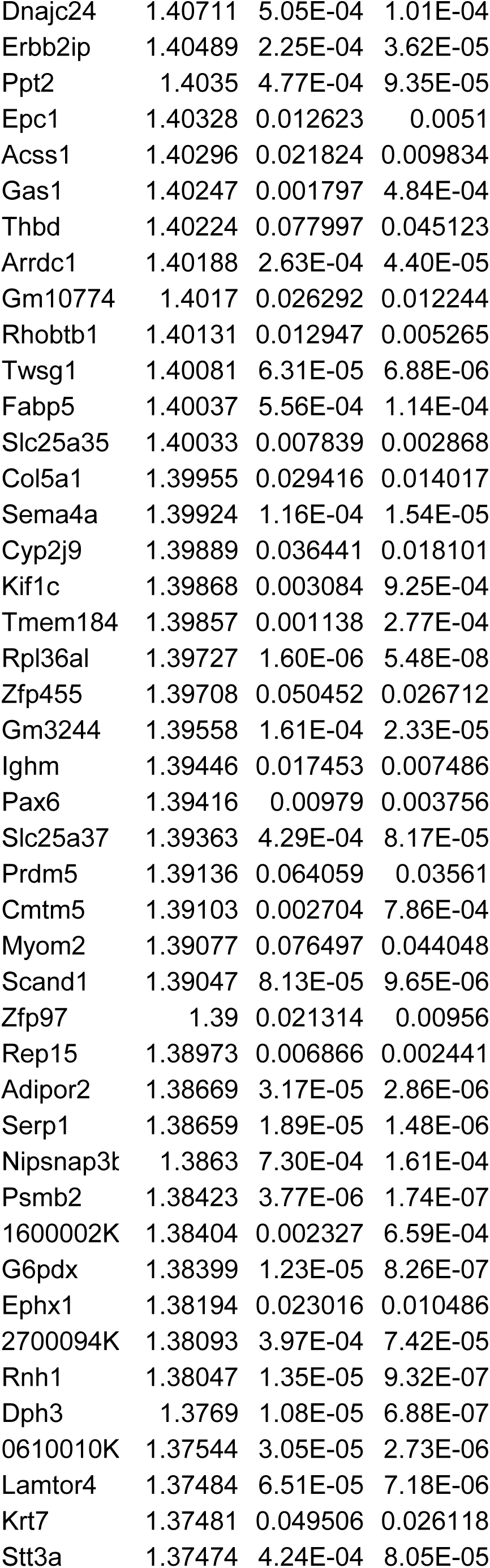
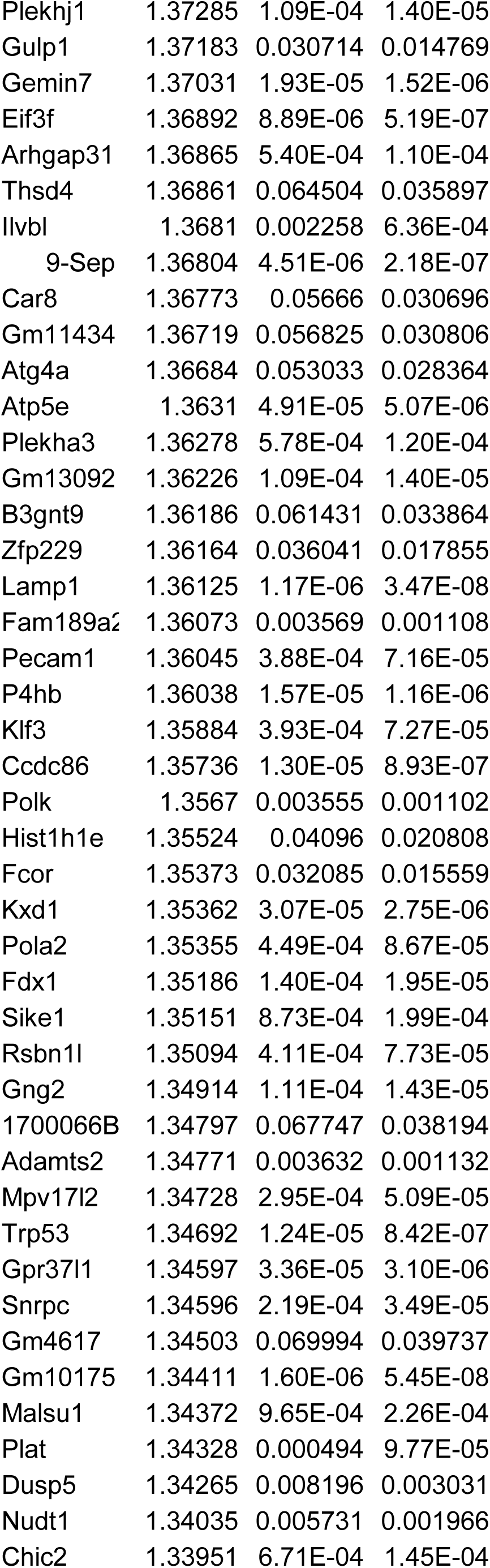
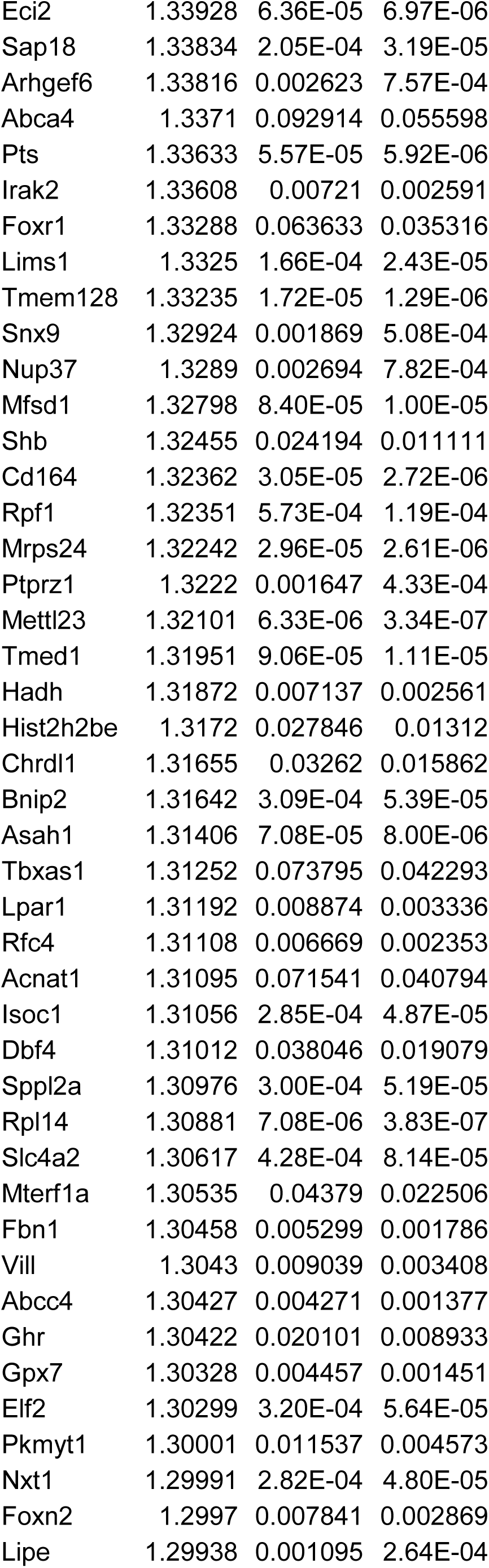
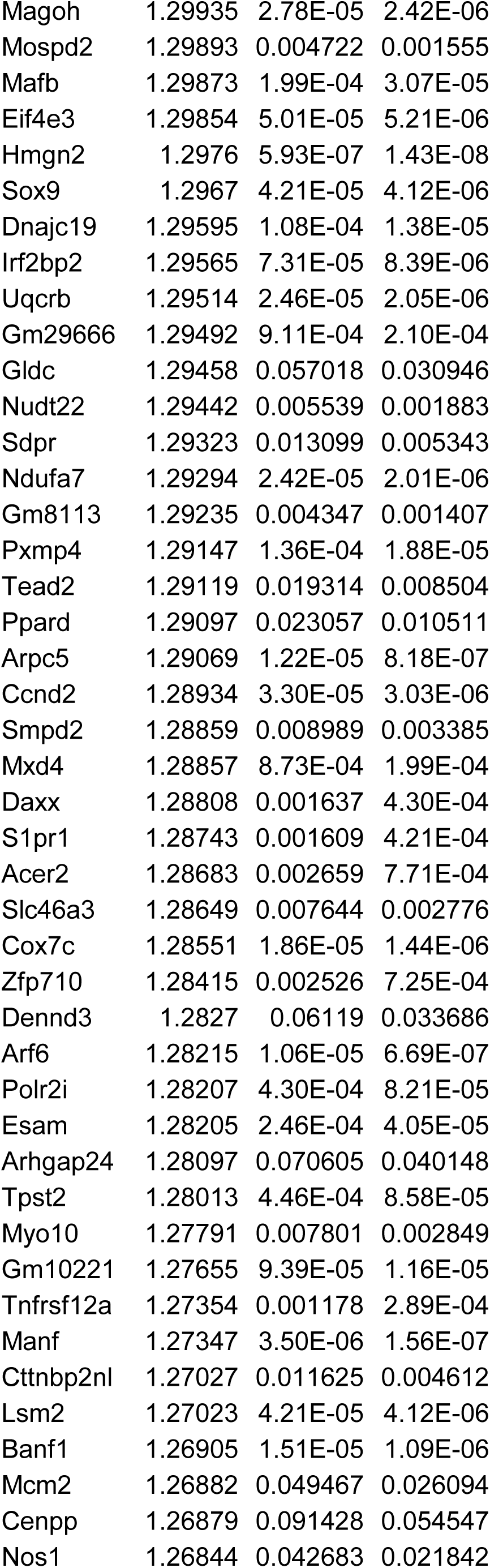
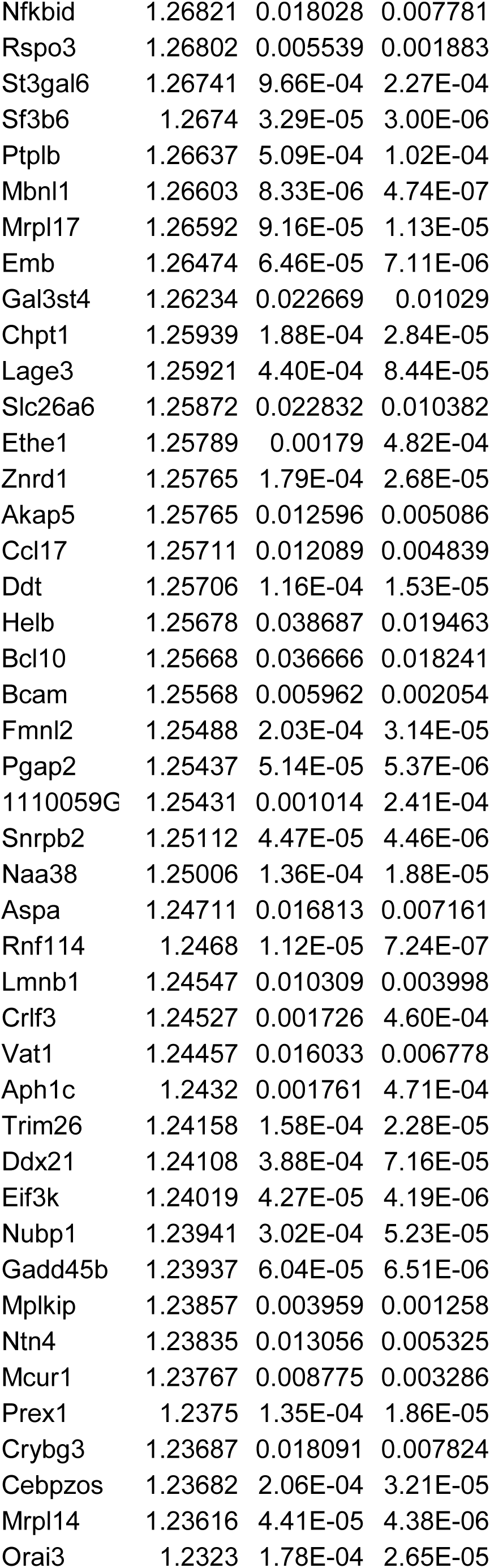
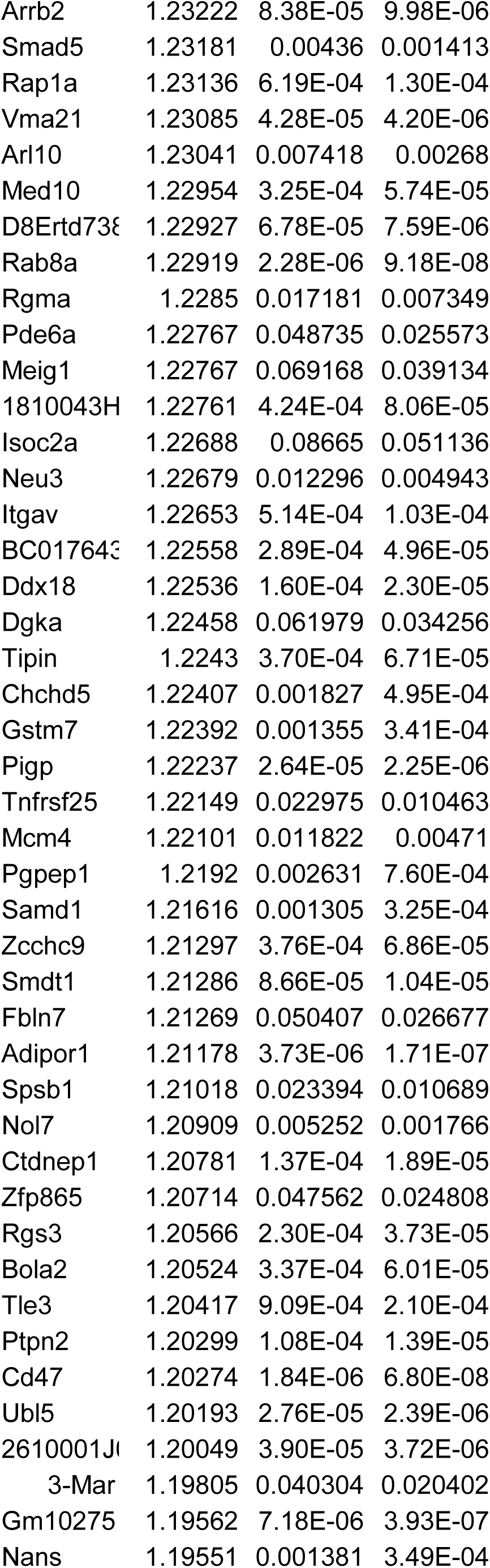
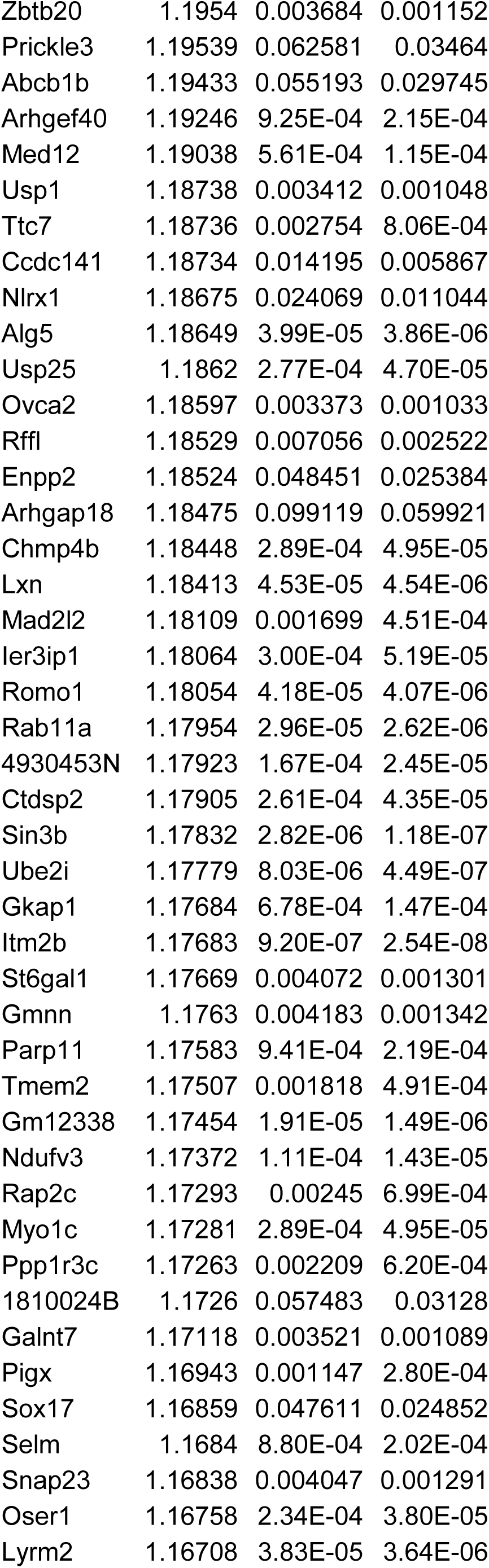
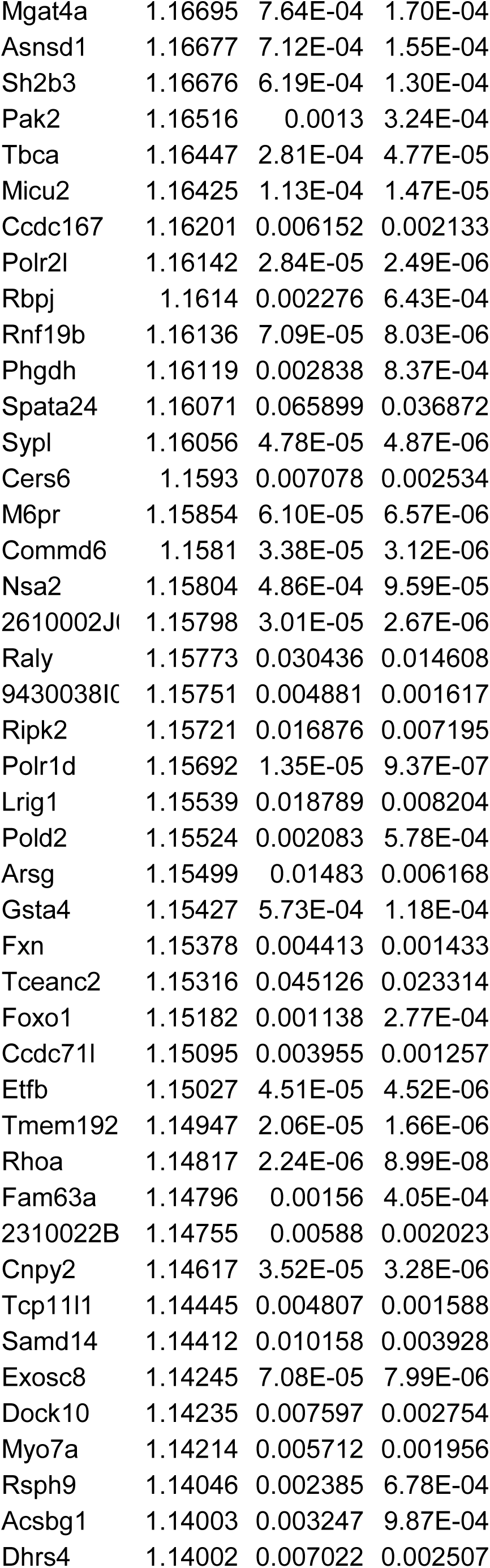
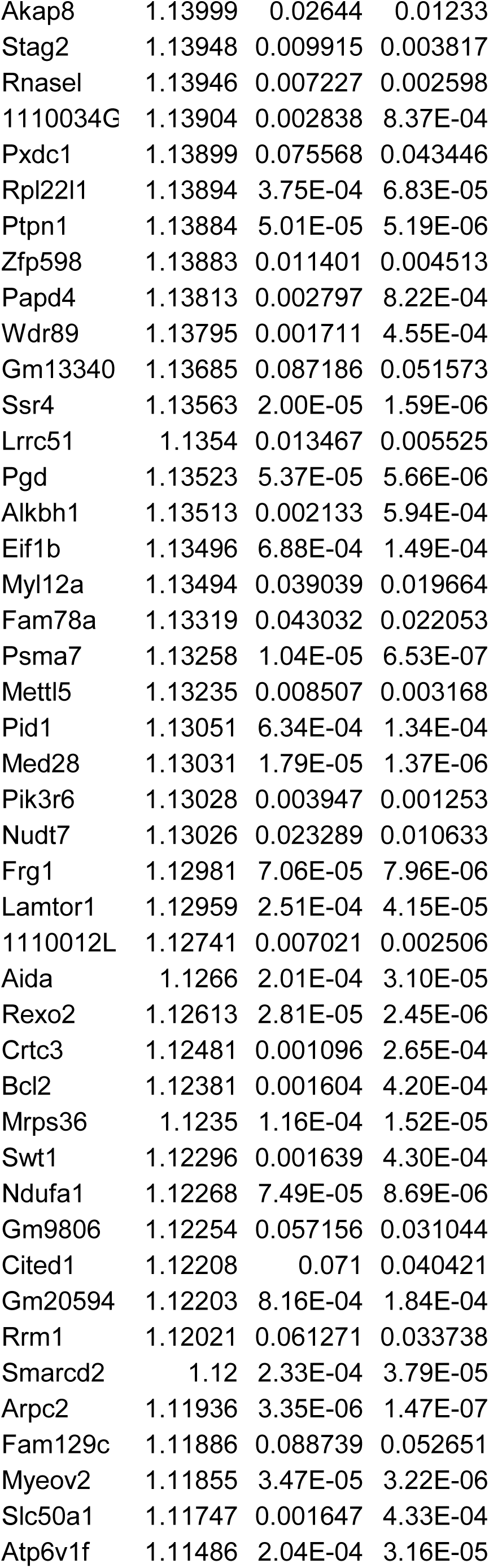
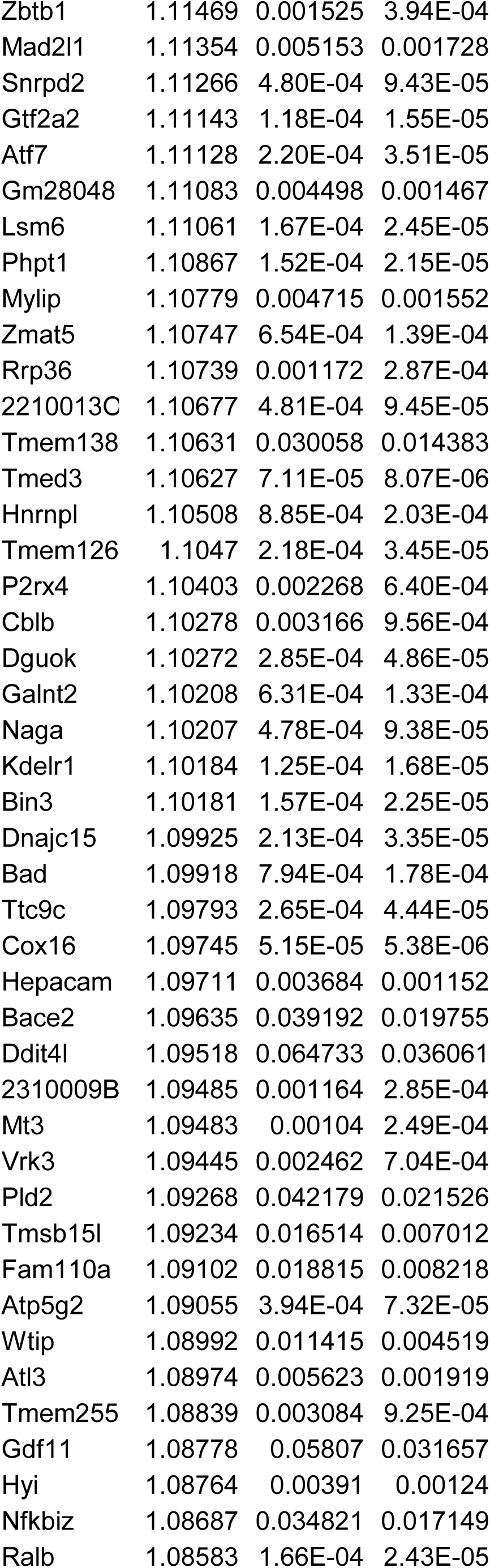
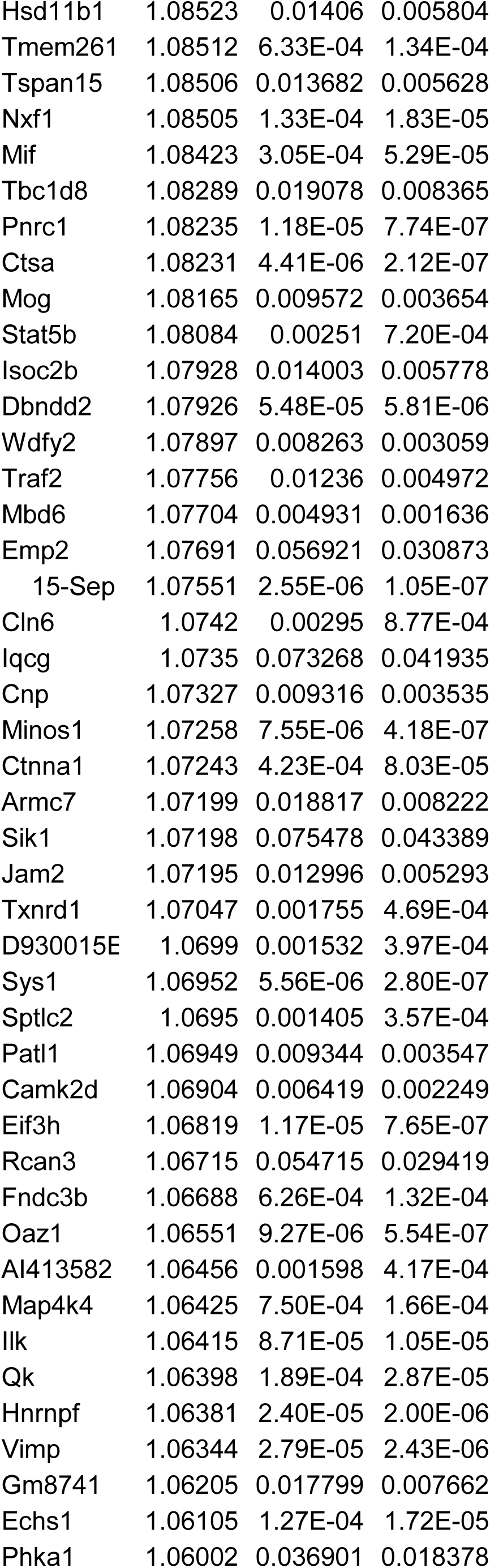
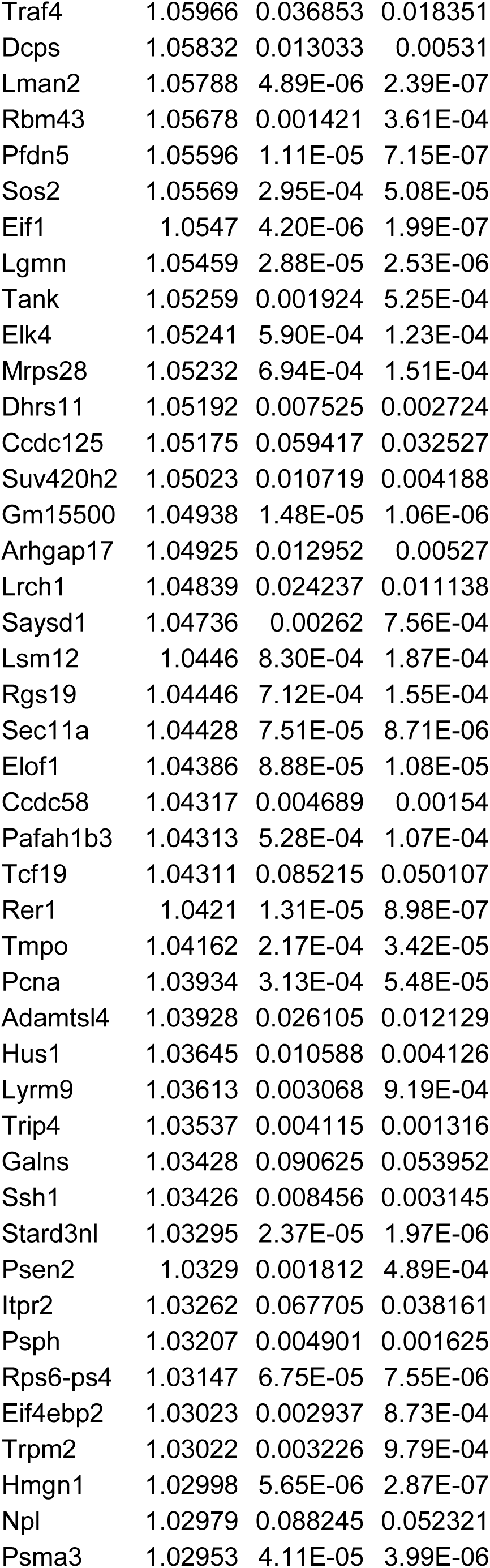
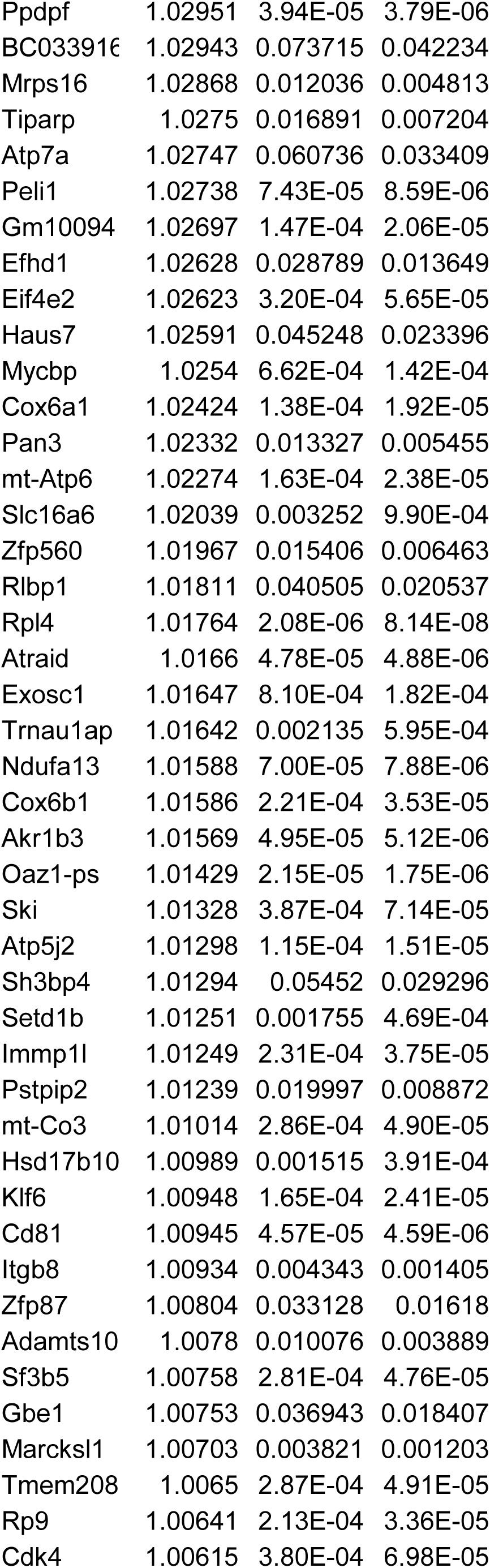
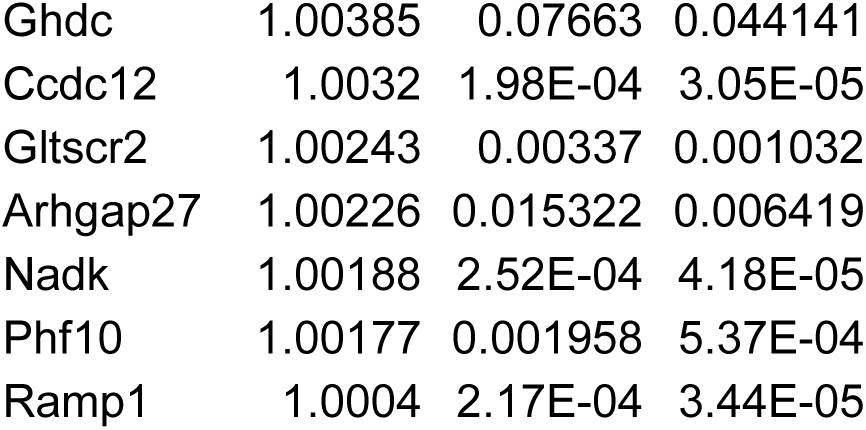
Differentially expressed genes from *T. gondii* infected or injected neurons from laser capture microdissection dataset. Values are Log2FC.

**Table 2.**
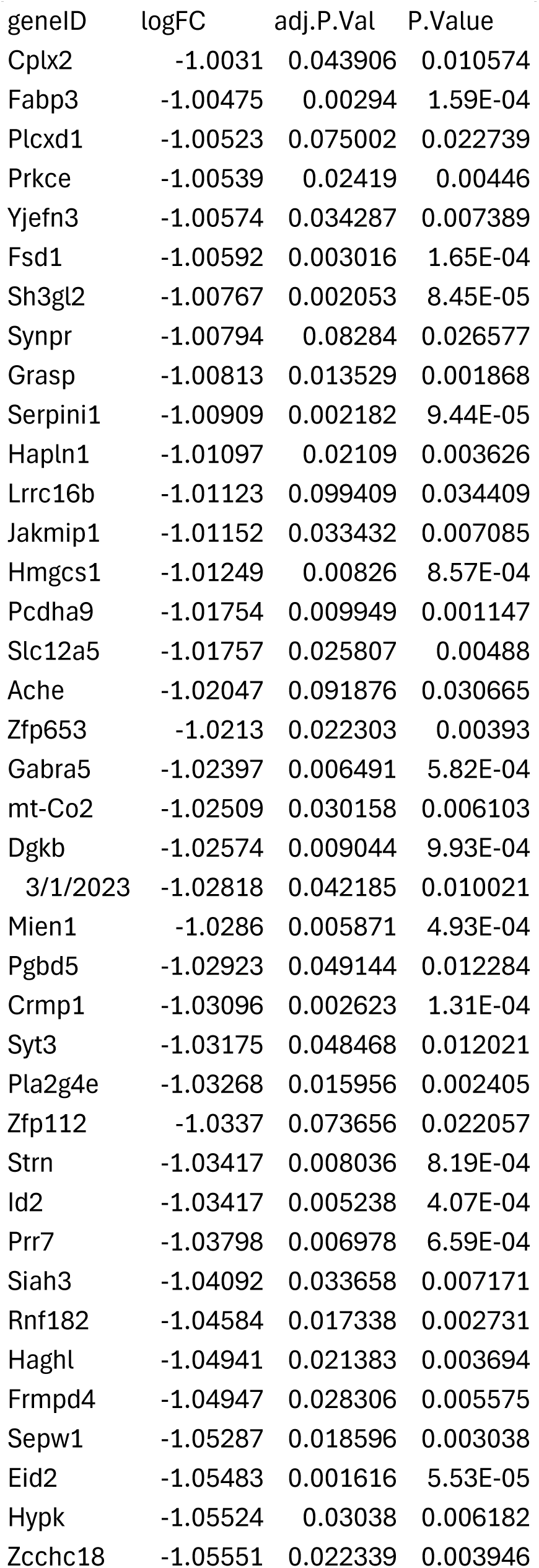
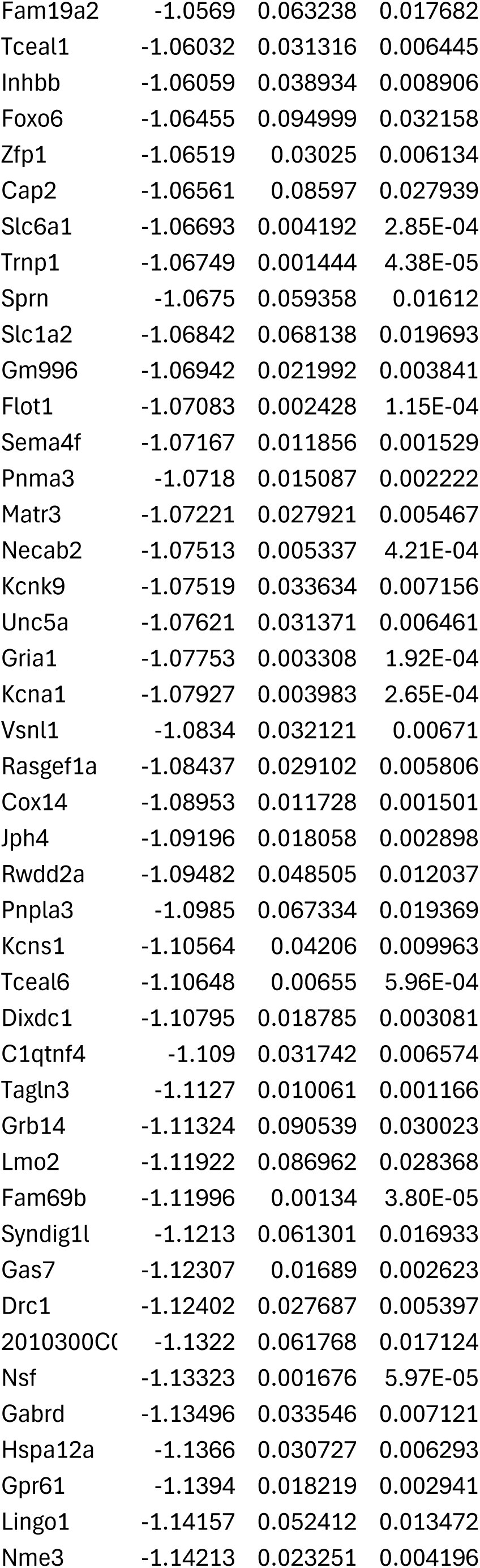
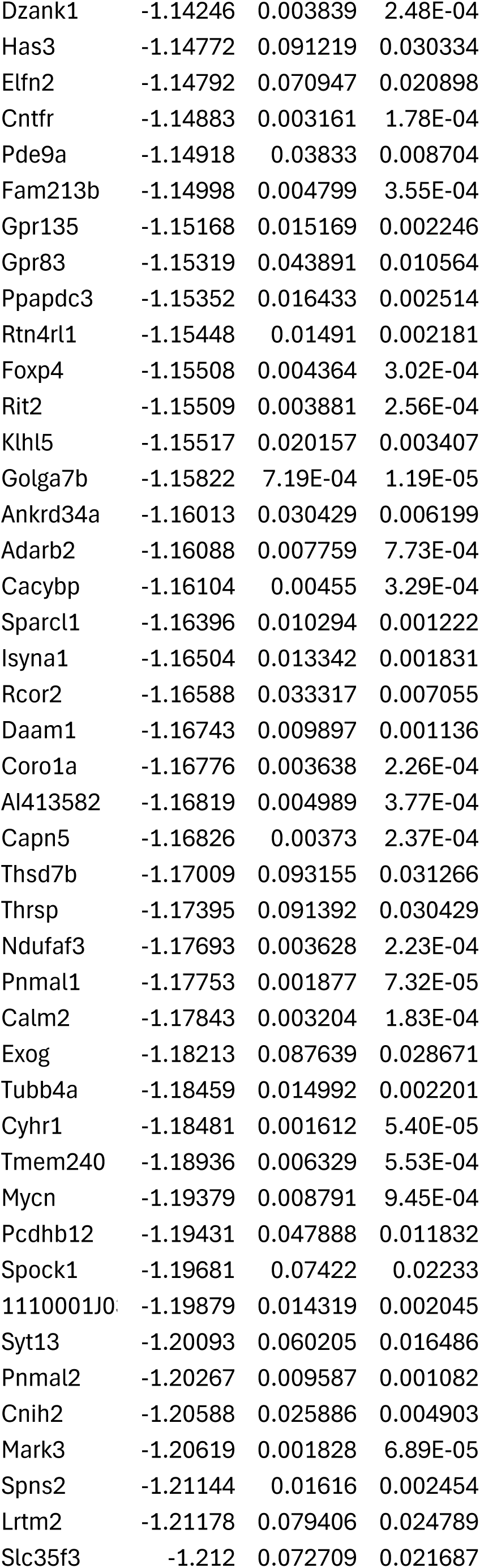
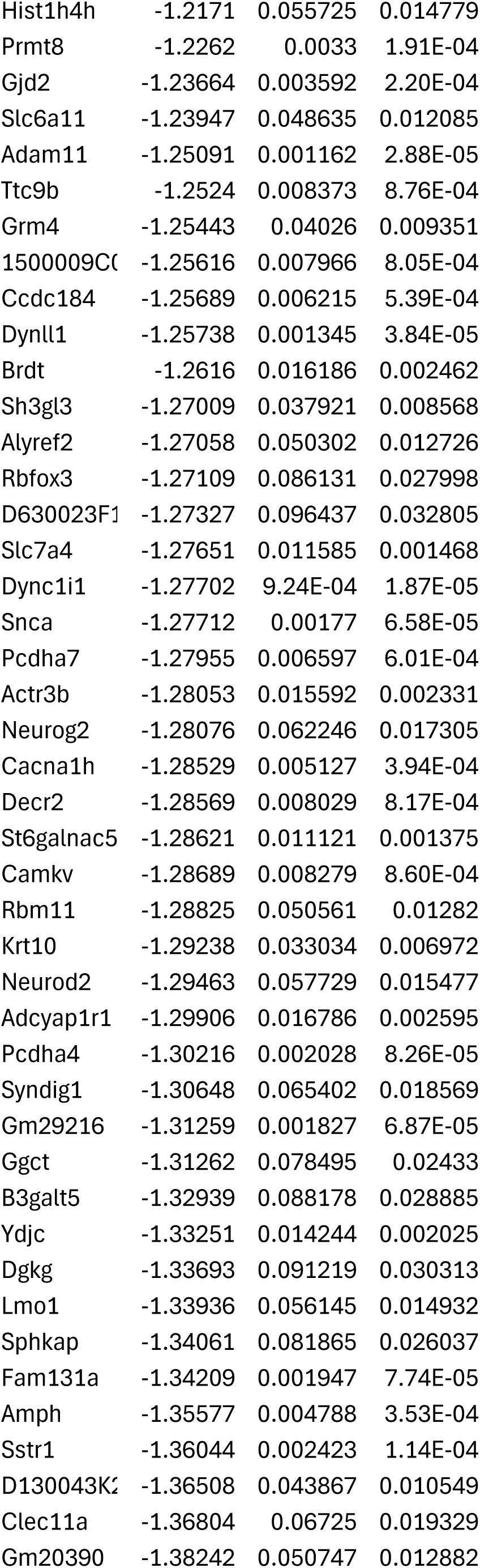
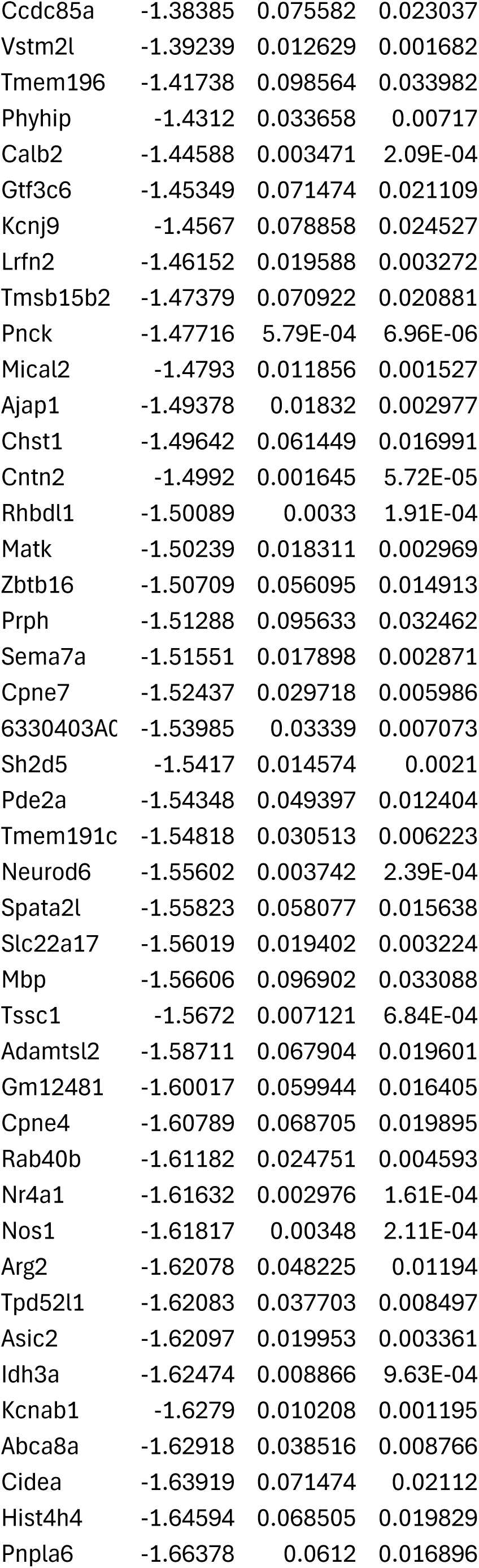
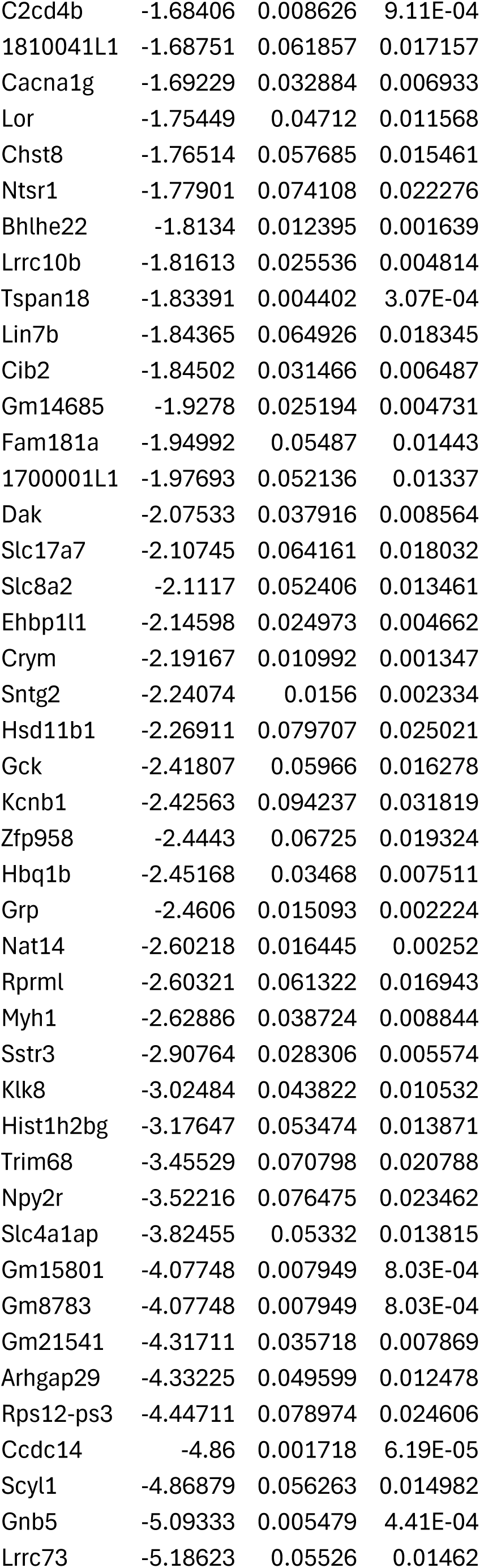
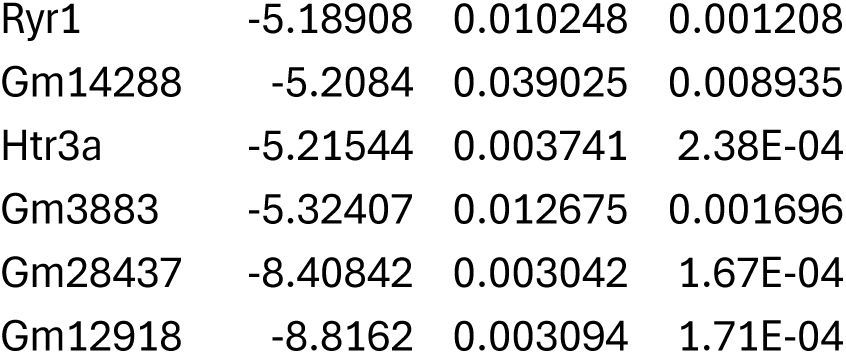
Differentially expressed genes from *T. gondii* infected or injected neurons from primary neuron cultures. Values are Log2FC.

As noted above, the transcriptomes from TINs contained transcripts classically associated with immune cells^24^. Such transcripts were not observed in PNCs (**Fig. S2**), except for *Cd80* and *Cd44* which are receptors that are expressed during neuron development^28,29^. Thus, the pathways identified in the PNCs data that are also identified in the LCM data likely represent neuron specific responses to *T. gondii*. Functional Enrichment Analysis of both sample types revealed 975 upregulated pathways shared between infected PNCs and neurons captured *in vivo* (**Fig. 2A**). A comparison of the top 50 enriched pathways of each condition (LCM or PNCs) identified 14 pathways in common (**Table 3**). These pathways were associated with responses to different microbial stimuli (e.g., LPS, COVID, RSV) and cytokine signaling. By analyzing the individual genes associated with these pathways, we identified a small group of genes that were consistently enriched in these 14 pathways. These genes included CXC motif chemokine ligand 10 (*Cxcl10*, 12 of the 14 pathways), chemokine (C-X-C motif) ligand 1 (*Cxcl1*, 10 of the 14), and chemokine (C-C motif) ligand 2 (*Ccl2*, 9 of the 14) (**Fig. 2B**). These pathways were functionally similar in that they primarily centered around chemokine/cytokine signaling and proinflammatory responses (**Fig. 2C**). While both PNCs and the LCM data showed an enrichment of these pathways, the LCM dataset showed a higher number of genes involved (set size) and increased Log_2_FC of the DEGs (**Fig. 2C**). Only *Ifih1*, *Plaur*, and *Tnfaip3* were highly represented genes that were equivalent or higher in PNCs versus LCM neurons.

**Figure 2.**
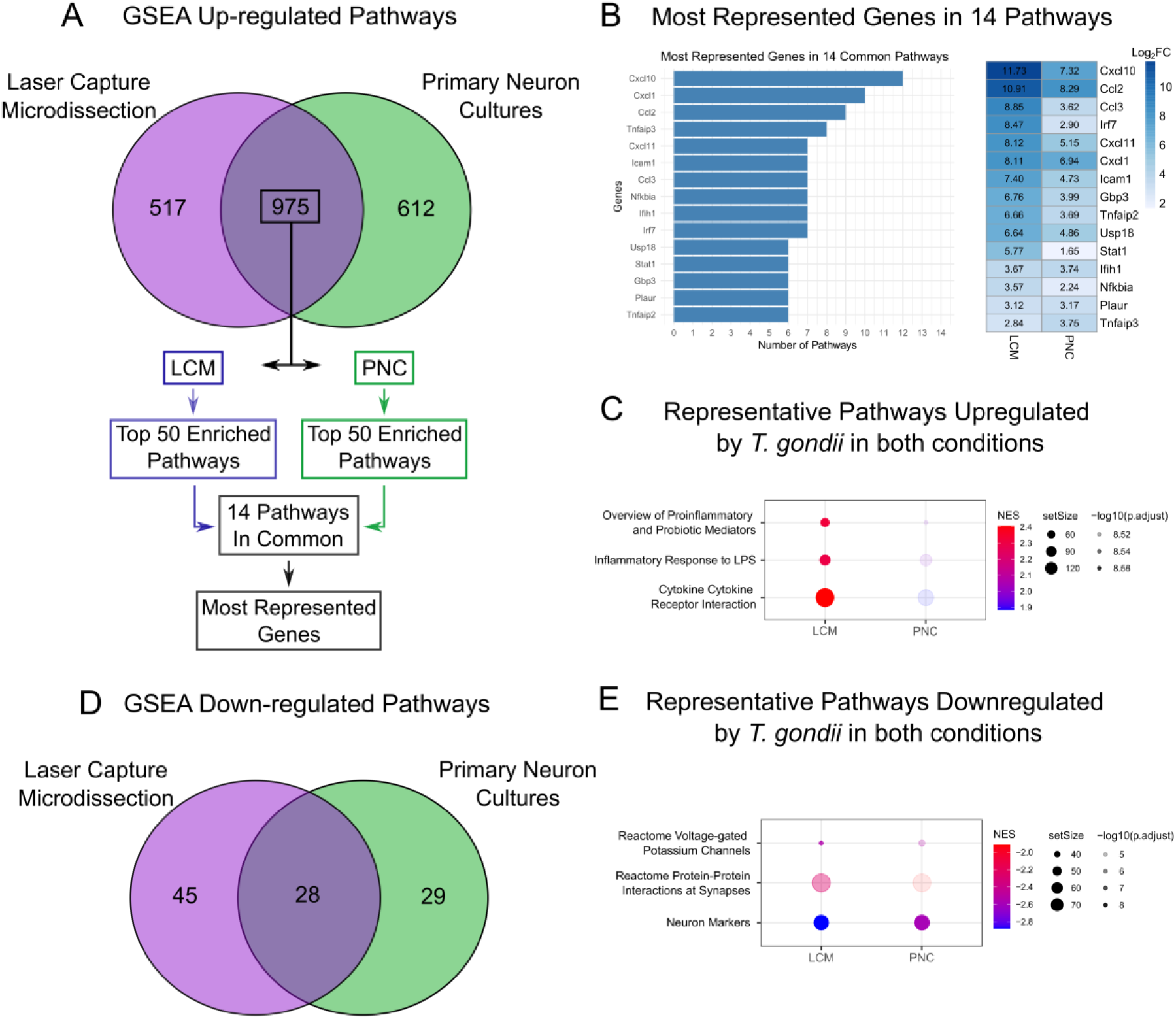
Pathway analysis reveals neuronal response to *T. gondii* infection involves an increase in pro-inflammatory cytokines and a decrease in neuron function. (**A**) The top 14 enriched pathways between LCM dataset and PNCs were selected out of 975 upregulated pathways. (**B**) Quantification of the most represented genes in the 14 most enriched pathways with a heatmap of their Log2FC. (**C**) Representative signature pathways between PNCs and LCM with normalized enrichment scores (NES). (**D**) Venn diagram of 28 downregulated pathways in LCM and PNCs datasets. (**E**) Enrichment scores of downregulated neuron pathways in *T. gondii* datasets.

**Table 3.**
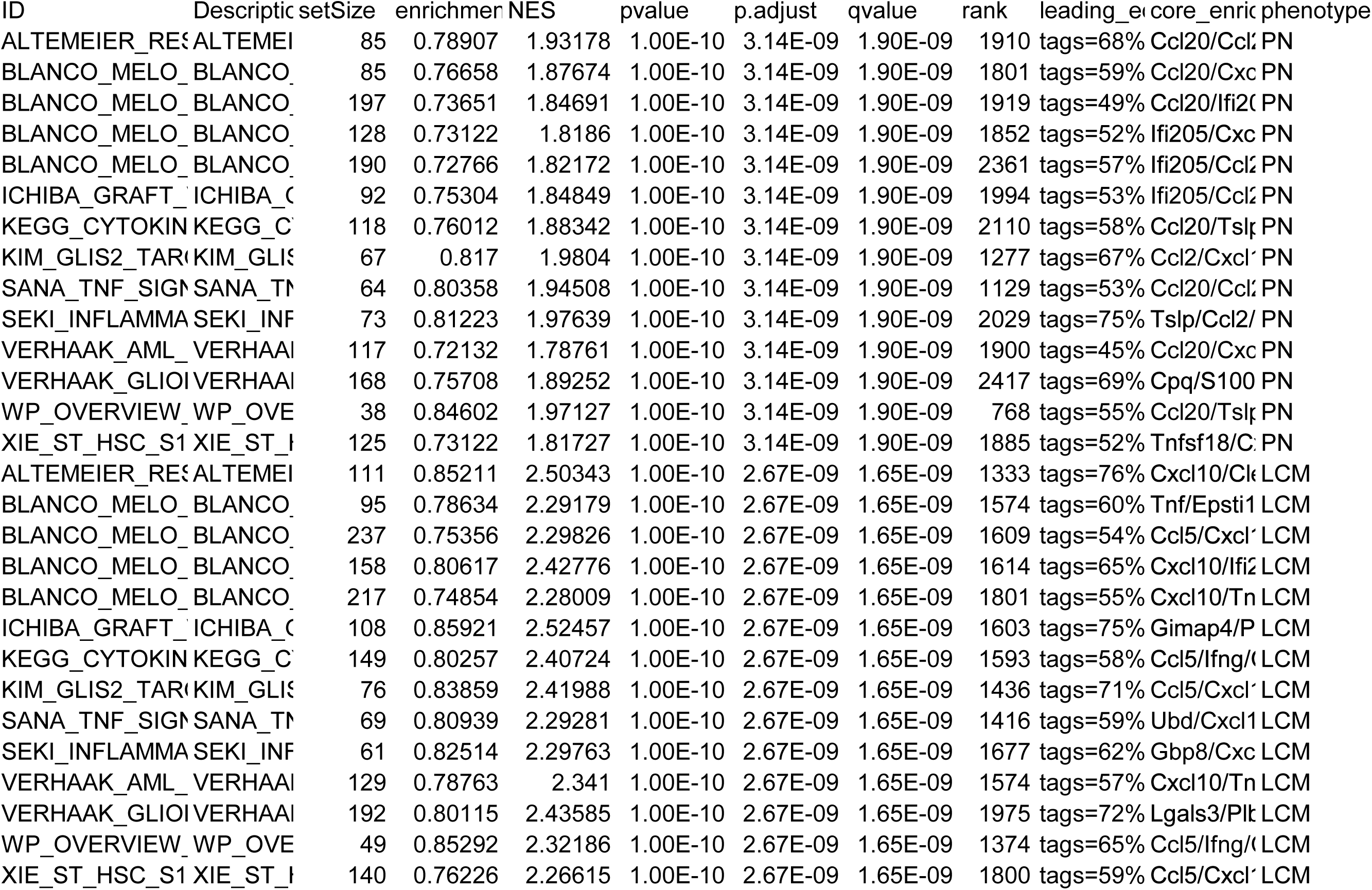
Fourteen pathways shared in the top 50 enriched pathways of each condition (LCM or PNC).

A comparison of the downregulated pathways between LCM and PNCs found 28 pathways in common (**Fig. 2D**). Many of these pathways were neuron-specific, including neuronal markers, protein-protein interactions at synapses (e.g., SNARE proteins), long-term potentiation, activation of NMDA receptors, and GABA synthesis and receptor signaling. We had previously noticed fewer neuron markers in our LCM dataset but could not determine if this decrease was due to an increase in contaminating immune cells comprising a higher proportion of our transcripts or a true decrease in neuronal transcription. However, in the PNCs—which lack immune cells—we still saw a decrease in these neuron-specific pathways and related genes (**Table 4**). In addition to synaptic and neuron marker pathways, multiple voltage-gated potassium channels were downregulated (**Fig. 2E**). Using the genes from the GO pathway GOMF_POTASSIUM_ION_LEAK_CHANNEL_ACTIVITY, we found that many were downregulated in both of our datasets (**Fig. S3**) except for Kcnk5/TASK-2, which was upregulated in both paradigms. In summary, the ability to compare the *in vivo* and *in vitro* datasets appears to be a feasible way to identify neuron-specific responses from complex *in vivo* transcriptional studies and suggests that *T. gondii* may directly modulate neuronal function.

**Table 4.**
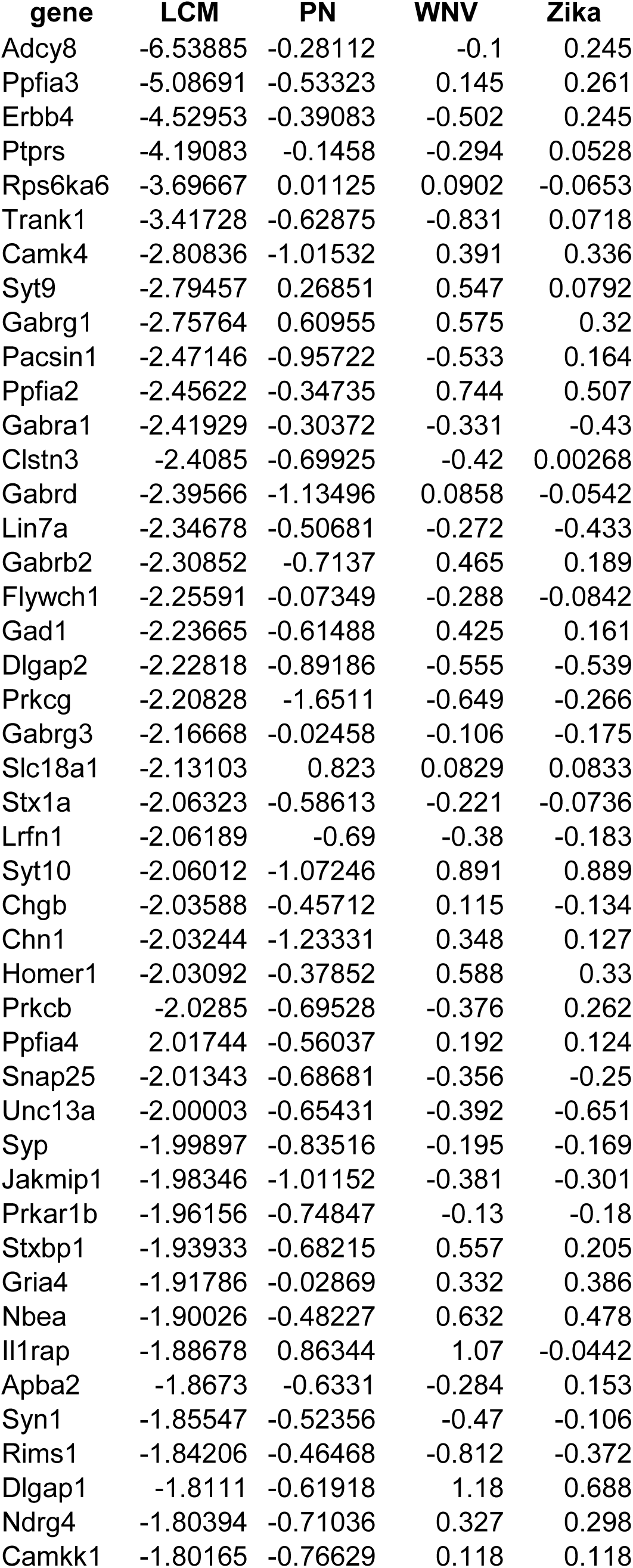
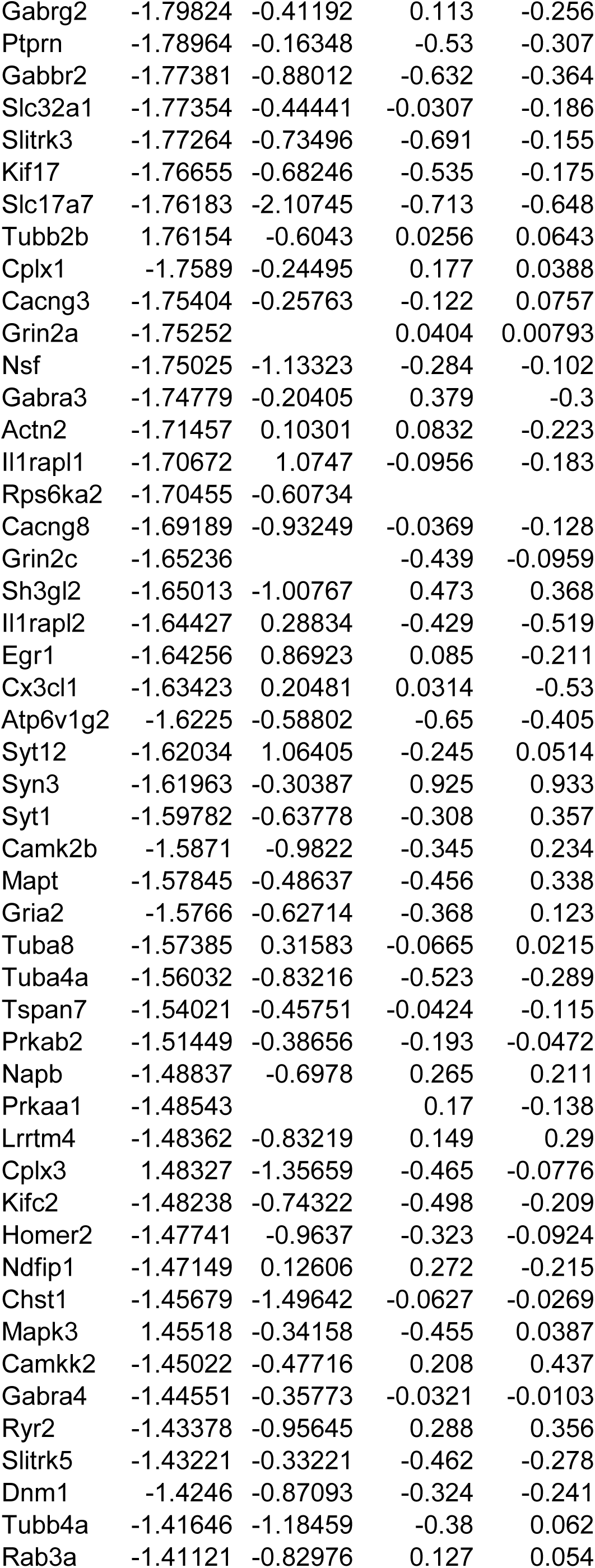
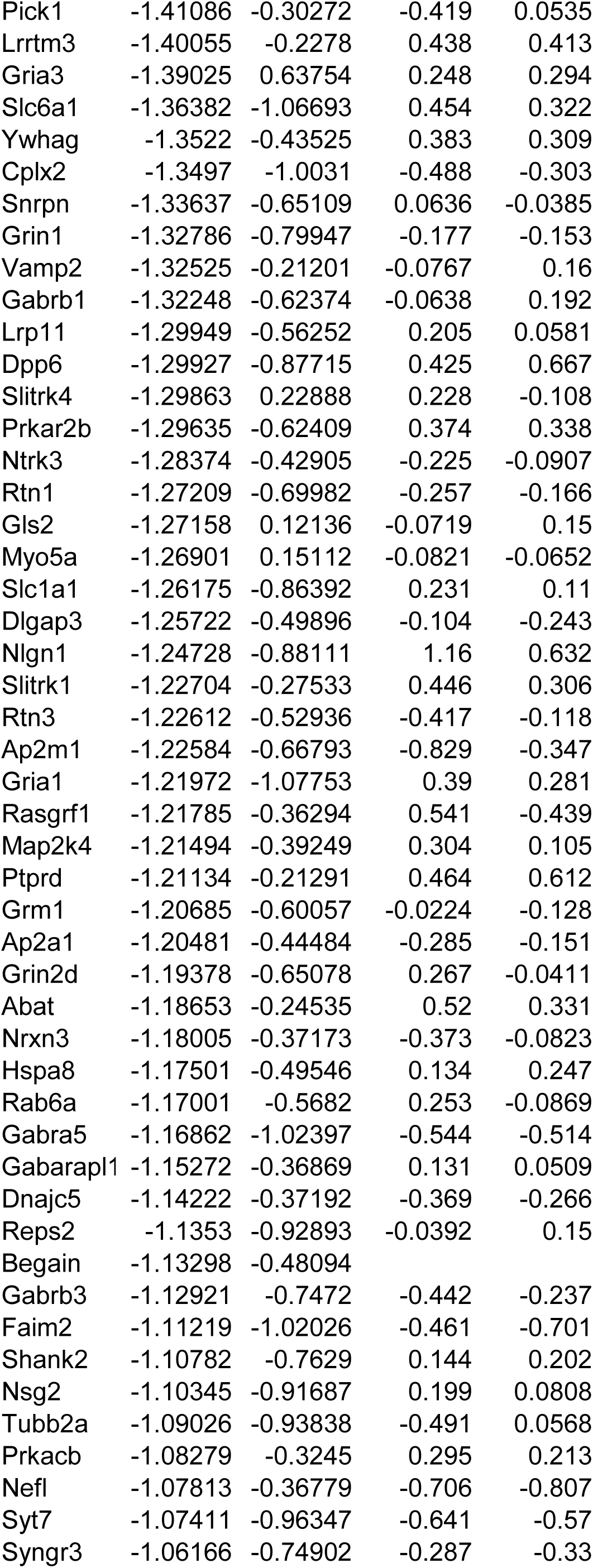
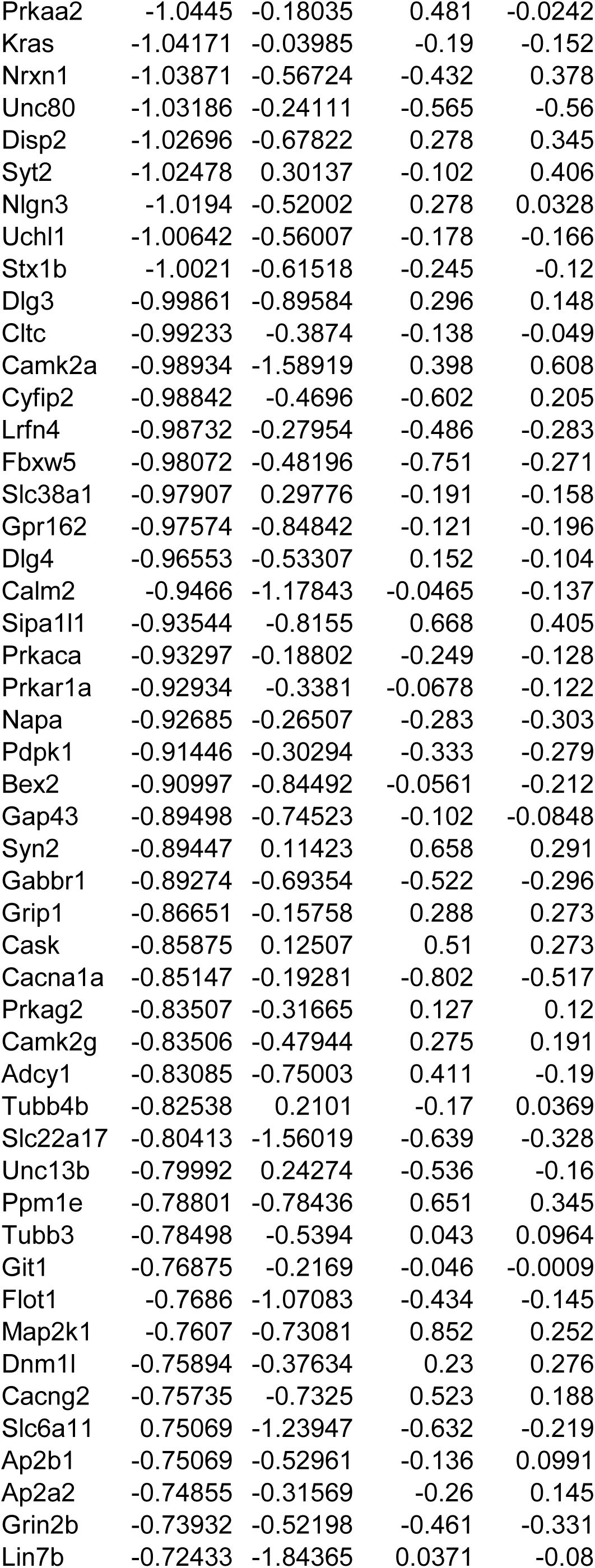
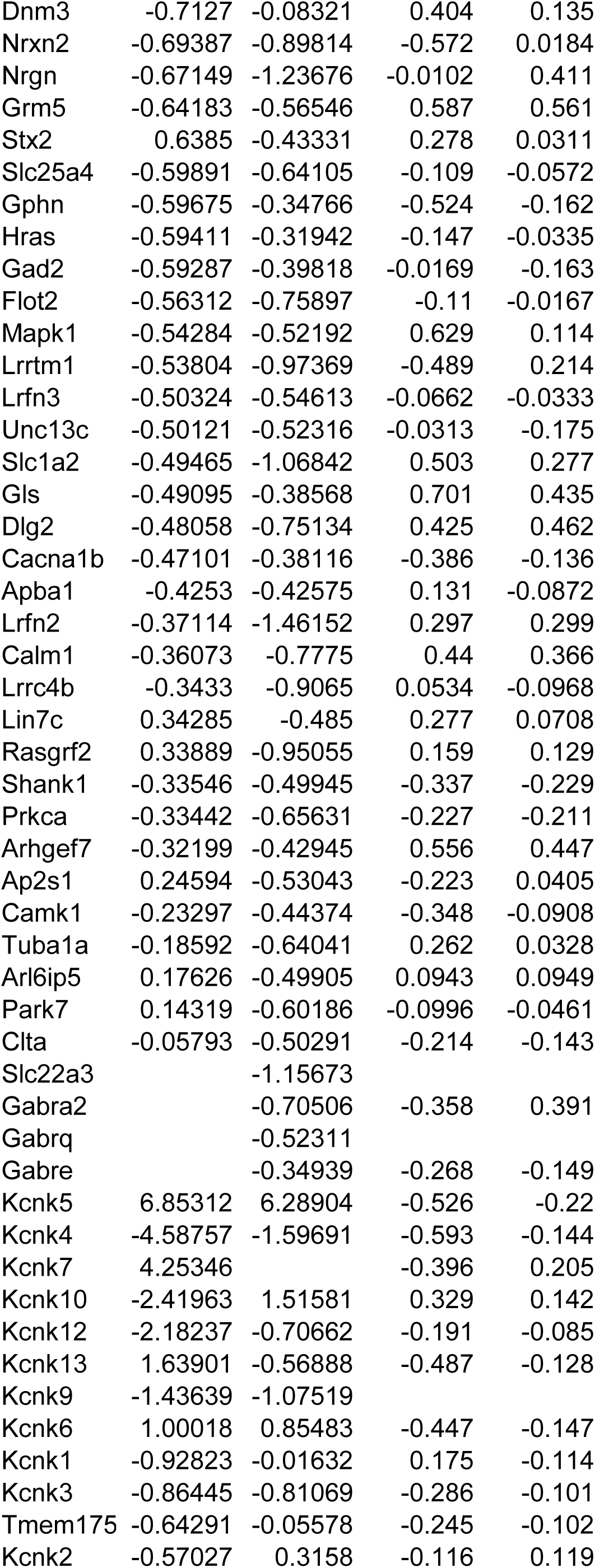
Neuron-associated genes from pathway analysis across four datasets. Values are Log2FC.

### Comparison of neuronal responses to *T. gondii* or viral infection

To determine if these neuron responses were specific to *T. gondii* or occurred with other relevant infections we wanted to compare the LCM and PNCs datasets with transcriptional studies from neurons infected with non-parasitic microbes. A search of the Gene Expression Omnibus (GEO) repository, the NIH’s publicly funded genomics data repository, identified several transcriptional studies on infected wildtype murine neurons^7,30–32^. From these studies, we analyzed 4 datasets: Zika Virus (ZKV) and West Nile Virus (WNV) infected PNCs profiled by microarray^7^ and two single cell RNA seq studies of cortical neurons infected with a circuit tracing, attenuated rabies virus^31,32^. The latter two datasets^31,32^ had very few genes that met our criteria for DEGs (FDR ≤ 0.1, Log_2_FC ≥ 1) and were excluded from further analysis. However, the datasets for ZKV and WNV^7^ contained 193 and 690 DEGs respectively (**Fig. 3A**), making them amenable for comparison with the *T. gondii* datasets.

**Figure 3.**
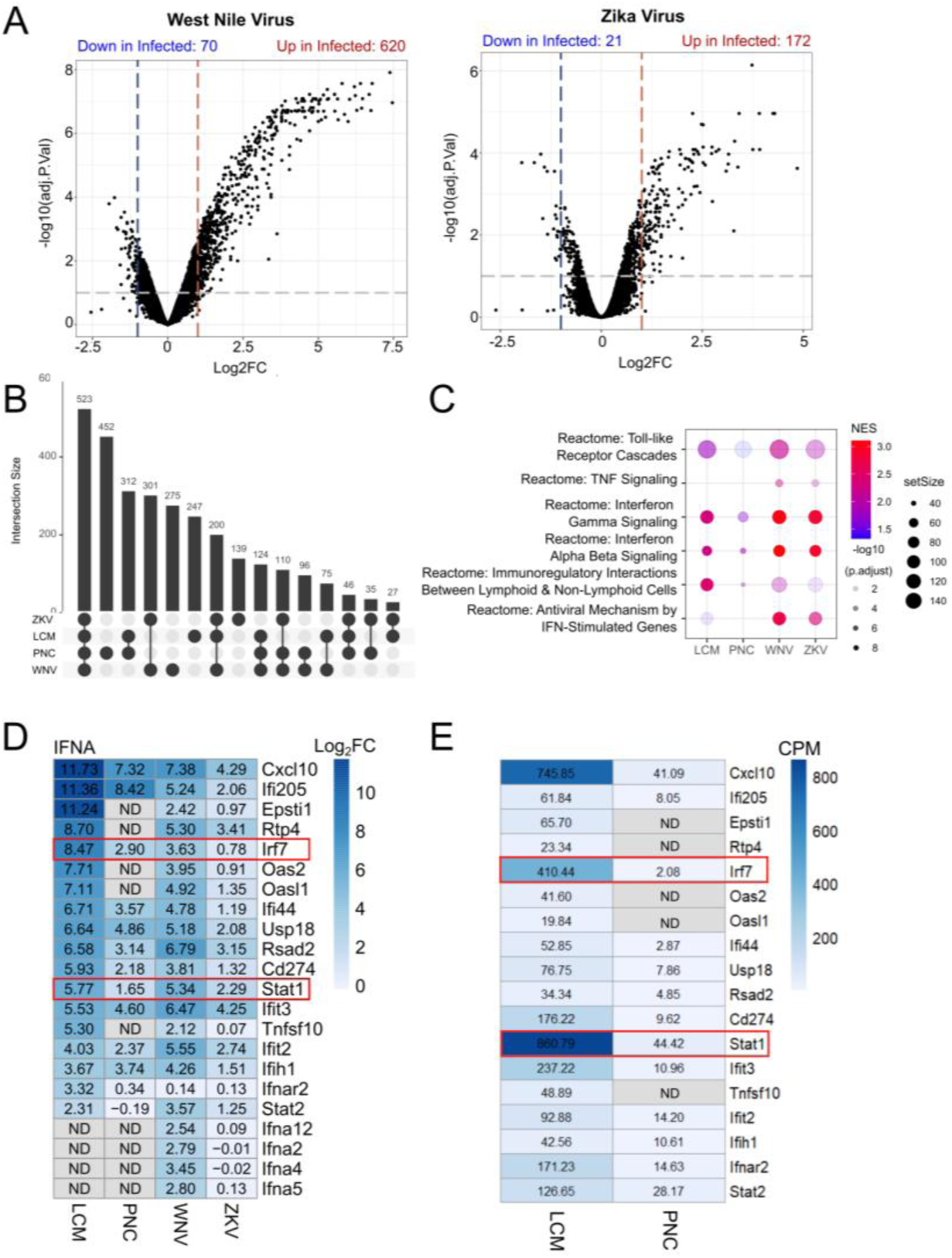
WNV and ZKV-infected PNCs and the LCM dataset show upregulation of IFN*-α* response genes, unlike *T. gondii*-infected PNCs. **(A)** Volcano plot of DEGs for WNV and ZKV. (**B**) Upset plot of upregulated GSEA pathways in *T. gondii-*infected PNCs, LCM data, WNV-infected PNCs, and Zika-infected PNCs (**C**) Relative expression of inflammatory pathways across datasets. (**D**) IFN-α response genes expressed in LCM, *T. gondii*, WNV, and ZKV-infected in Log2FC. (**E**) Count per million of IFN-α response genes in LCM and *T. gondii* PNC datasets with raw values shown. ND = Not Detected. ND genes in both LCM and PNCs in D are not included in E.

Between these datasets there were 532 pathways in common (**Fig. 3B**). We further narrowed our focus to pathways relating to IFN-γ, IFN-α/β, and TNF signaling to see if there were differences in these responses between infections (**Fig. 3C**). We found that all datasets had an enrichment for the IFN-γ signaling pathway and innate immunity pathways (represented by Toll-like Receptor Cascades) (**Fig. 3C**) but only the viral datasets showed enrichment for TNF signaling. As expected, the viral datasets showed an enrichment in anti-viral, type I IFN pathways, specifically in IFN-stimulated genes (ISG). Consistent with prior work^33^, *T. gondii* infected PNCs showed very little type I IFN response, while the LCM dataset showed some enrichment in the “Antiviral Mechanism By IFN-related antiviral mechanisms” (**Fig. 3C, Fig. S4**). To understand these differences, we compared the individual genes involved in type I IFN pathways and found that several key IFN-α genes were differentially expressed (**Fig 3D, E**). *T. gondii* PNCs showed upregulation in *Stat1* and *Irf7* but not in many downstream genes. These downstream genes fell into two clusters, with the first cluster (e.g., *Oas2*, *Oas1l*, and *Epsti1*) showing no baseline expression or upregulation and the second cluster (e.g., *Ifnar2*, *Stat2*) showing low baseline expression and no upregulation (**Fig. 3E**). WNV, ZKV, and, to a lesser extent, *in vivo* infection with *T. gondii* showed upregulation of many of the genes in this pathway, though differences could be seen even between these 3 experimental conditions (e.g., *Ifna2,4,5,12*) (**Fig 3D**). Collectively, these data suggest that *in vitro*, WNV and ZKV trigger neuron IFN-α responses but *T. gondii* does not, while *T. gondii* triggers a broader immune response *in vivo*.

## Discussion

Here we sought to define how neurons respond to *T. gondii* and determine how this response compares to infection with other neurotropic microbes. To accomplish this goal, we compared four transcriptional datasets: *T. gondii*-injected neurons (TINs) captured by LCM from infected murine brain tissue and primary murine cortical neuron cultures (PNCs) infected with *T. gondii*, WNV, or ZKV. These comparisons revealed that cortical neurons have conserved responses to these infections but also show key differences that distinguish virus versus eukaryotic parasite.

All four datasets had a pronounced increase in inflammatory pathways, including in type I and type II interferon signaling (**Fig. 3c**). Of the many cytokines/chemokines upregulated in these datasets, *Cxcl10* is highly represented in the pathways upregulated in the LCM and *T. gondii*-infected PNCs datasets and is also upregulated in WNV and ZKV-infected PNCs. (**Fig. 4**). These data suggest *Cxcl10* upregulation is a conserved feature of the neuron response to these infections. As *Cxcl10* is a chemokine that attracts innate and adaptive immune cells, its conserved upregulation is consistent with the need to attract immune cells to infected neurons, whether the infecting microbe is viral or parasitic (e.g., effector T cells for *T. gondii*^34^).

**Figure 4.**
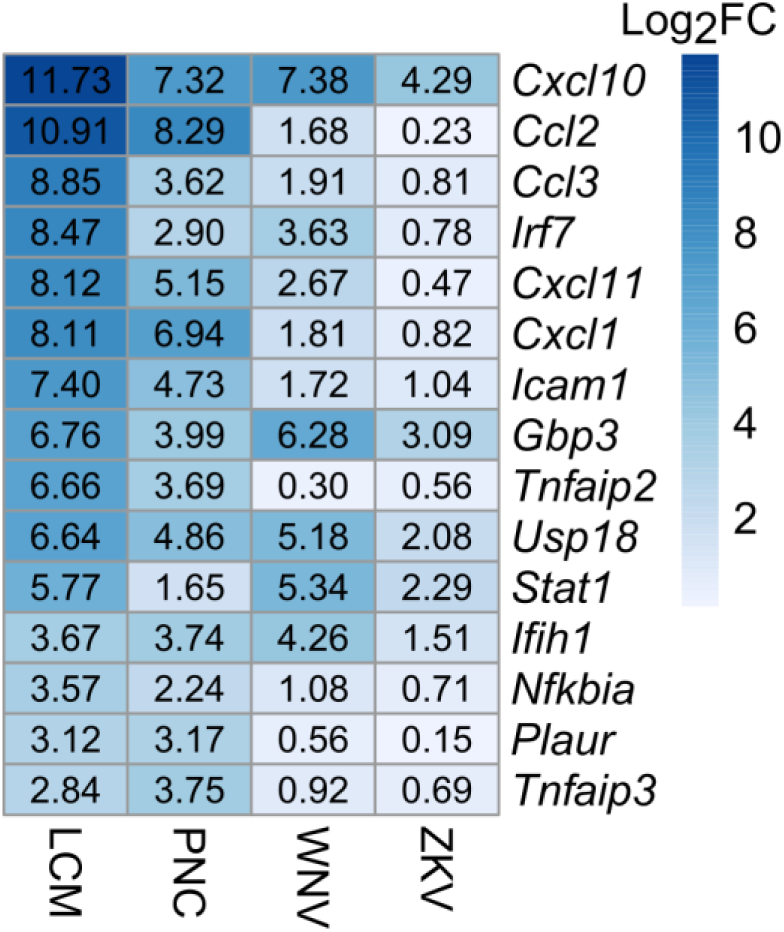
*Cxcl10* is upregulated across acute and subacute datasets. Other conserved genes include *Irf7, Icam1, Gbp3, Usp18, Stat1, and Ifih1*.

The differences between the datasets are also revealing. Only the *T. gondii* infection datasets showed a consistent downregulation of neuronal function pathways (markers, LTP, synapse function, potassium channels). The downregulation in potassium channels was of particular interest to us because it could explain our recent finding that TINs have a depolarized resting membrane potential when compared to non-injected neighboring neurons or neurons in uninfected mice^35^. Neuronal dysfunction associated with *T. gondii* infection has been identified previously^36–41^, but the experimental setups made it a challenge to distinguish the direct effect of *T. gondii* on neurons versus effects from infiltrating immune cells or microglia and astrocytes. The findings presented here suggest that *T. gondii* can directly induce neuronal dysfunction.

Another interesting example of infection-dependent effects is IFN-α signaling. Akin to other type I IFN responses, IFN-α signaling begins with activation of a host cell pattern recognition receptor (PRR) by a pathogen. This activation leads to IRF7 phosphorylation, resulting in the upregulation of IFN-α. Once released, IFN-α binds to the Interferon alpha and beta receptor (IFNAR), which allows IFN-α to act in an autocrine and paracrine fashion. IFNAR activation leads to the phosphorylation of STAT1 and STAT2, that mediate the transcriptional upregulation of a specific set of downstream IFN-α response genes^42^ (**Fig. 5**). As expected, the virus-infected PNCs showed upregulation in genes throughout this pathway, but the two *T. gondii* datasets were less consistent. Both *T. gondii* datasets showed an upregulation in IRF-7, but no upregulation of type I IFNs (alpha or beta) (**Fig S4C**). These findings are consistent with prior work in human fibroblasts that suggest that *T. gondii*-infected cells block type I IFN responses^33^ upstream of the *T. gondii* effector TgIST, which prevents IFN signaling by binding to pSTAT1/2 heterodimers and pSTAT1 homodimers^43,44^ (**Fig 5**). That the *in vivo T. gondii* dataset shows upregulation of some of the downstream IFN response genes suggests that autocrine and paracrine signaling from neuronal and non-neuronal cells may overcome this inhibition *in vivo* or these downstream genes are upregulated by other pathways. Collectively, these findings are consistent with a model in which neuronal responses to infection depend on the context, with conserved responses arising from pathogen sensors that converge on the same downstream pathways. Such sensors could detect microbes directly or through neuronal stress that, in turn, triggers cellular responses associated with pathogen recognition^45^.

**Figure 5.**
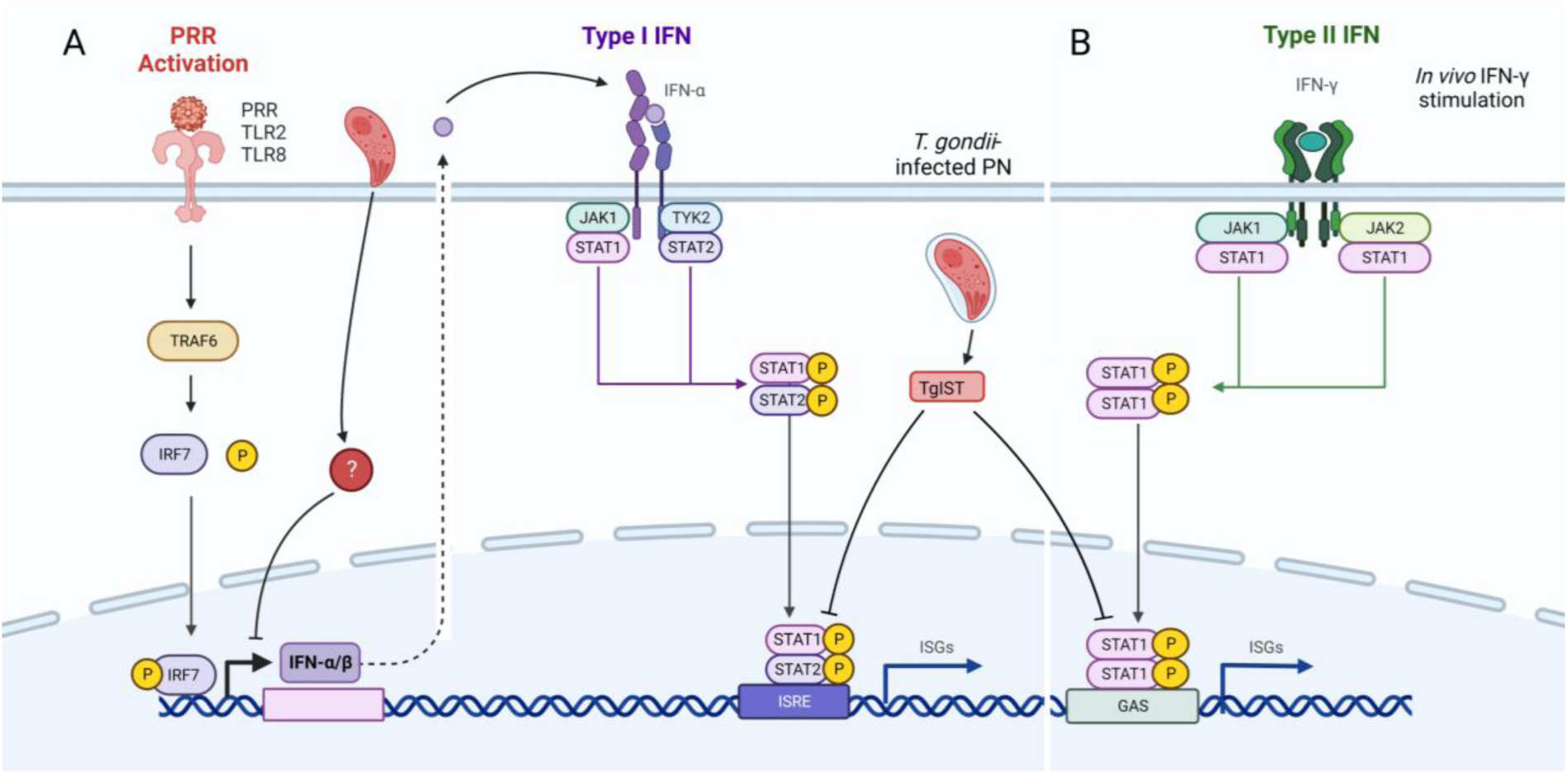
Despite *Irf7* upregulation in all datasets, *T. gondii-*infected neurons fail to upregulate and secrete IFN-α. (**A**) WNV and *T. gondii* activate PRR receptors leading to an intracellular cascade that results in IFN-α production through IRF7 phosphorylation. IFN-α binds IFNAR and upregulates *Stat1* and other subsequent IFN-α response genes in WNV-infected neurons. *T. gondii* inhibits *Stat1* in this pathway *in vitro* with *T. gondii* effector protein TgIST. The lack of *Ifn-α* upregulation in both *T. gondii* datasets indicate that the parasite may inhibit the action of phosphorylated IRF7 through an unknown mechanism; either before full invasion or after. (**B**) *In vivo, Stat1* may be upregulated through alternative stimulation pathways such as IFN-γ.

## Methods

### Parasite Maintenance

As previously described, type II *T. gondii* (Pruginaud) used in this study was maintained through serial passage in human foreskin fibroblasts (gift of John Boothroyd, Stanford University, Stanford, CA) using DMEM, supplemented with 10% fetal bovine serum, 2mM glutagro, and 100 IU/ml penicillin and 100 μg/ml streptomycin^27^.

### Mice

All procedures and experiments were carried out in accordance with the Public Health Service Policy on Human Care and Use of Laboratory Animals and approved by the University of Arizona’s Institutional Animal Care and Use Committee (#12-391). All mice were bred and housed in specific-pathogen-free University of Arizona Animal Care facilities. Cre reporter mice (ZsG mice) (mice stock no. #007906) were originally purchased from Jackson Laboratories. Mice were inoculated with 10,000 freshly syringe-lysed parasites diluted in 200 μl of UPS grade PBS.

### Primary murine neuron culturing

Mouse primary cortical neurons were harvested from E174 mouse embryos obtained from pregnant ZsG mice. Dissections of E174 cortical neurons were performed and primary neuronal cell cultures were generated by methods described previously with minor modifications^46^. The culturing plates were prepared by coating overnight with 0.001% poly-L-lysine solution (Millipore Sigma, P4707, diluted in water 1:10) for plastic surfaces and 100 μg/ml poly-L-lysine hydrobromide (Sigma, P6282, dissolved in borate buffer, pH 8.4) for glass surfaces. They were washed three times for ten minutes each with water and transferred to plating media (MEM, 0.6% D-glucose, 10% FBS). Neurons were seeded at 500,000 in 6-well plates for RNA-seq. Four hours after plating, full media exchange to neurobasal media (Neurobasal base media, 2% B27 supplement, 1% L-glutamine and 1% penicillin-streptomycin) was performed. On day *in vitro* (DIV) 4, neurons received a half volume media change of neurobasal media with 5 μM cytosine arabinoside (AraC, Millipore Sigma, C6645) to stop glial proliferation. One third media exchanges with neurobasal media occurred every 3-4 days thereafter. All the experiments were performed on 10 DIV neurons.

### RNA isolation, preparation of cDNA libraries and sequencing

Primary neuronal cell cultures were infected (MOI 5) for 24 hours prior to RNA isolation. Total RNA was extracted using Direct-zol RNA Miniprep Kit and protocol (Zymo Research, R2051). Samples were sent to Novogene for quality control, library preparation and sequencing. RNA quality was measured on an Agilent 2100. Only samples with an RNA Integrity (RIN) score of >7 were used. After the QC procedures, mRNA from eukaryotic organisms is enriched using oligo(dT) beads. For prokaryotic organisms or eukaryotic organisms’ long non-coding libraries, rRNA is removed using the Ribo-Zero kit. First, the mRNA is fragmented randomly by adding fragmentation buffer, then the cDNA is synthesized by using mRNA template and random hexamers primer, after which a custom second-strand synthesis buffer (Illumina), dNTPs, RNase H and DNA polymerase I are added to initiate the second-strand synthesis. Second, after a series of terminal repair, A ligation and sequencing adaptor ligation, the double-stranded cDNA library is completed through size selection and PCR enrichment. Paired end sequencing was performed on an Illumina NovaSeq 6000 at 20 million reads per sample. Initial QC and Adaptor trimming was performed by Novogene.

### RNA-seq analysis and data visualization

Analyses and visualizations were conducted as previously described^47^ using a combination of statistical computing environment R version 3, RStudio version 1.2.5042, Bioconductor version 3.1^48^, and Prism 9.4.1 per a previously published training tool^49^. Transcript-level count were summarized to genes using the TxImport package^50^ and mouse gene annotation package from biomaRt^51^. Data were filtered and normalized with the EdgeR package^52^ by the Trimmed Mean of M-values (TMM) method. Genes with less than one count per million in n of the samples (3 for PN, 5 for LCM) were filtered out. The VOOM function in Limma^53^ was used to variance stabilize the filtered, normalized data. DGE analysis was performed with Benjamini-Hochberg correction with Limma^53^. Gene Set Enrichment Analysis (GSEA) was done with the using GSEA software (Broad Institute, version 4.0.2)^54^ in R with the GSEABase package. Venn diagrams were generated with Venn Diagram tool from VIB / UGent Bioinformatics & Evolutionary Genomics at http://bioinformatics.psb.ugent.be/webtools/Venn/. Heat maps were generated in R with pheatmap. Microarray data from BioProject PRJNA503843^7^, GSE122121, WNV samples GSM3455732, GSM3455733, GSM3455734, ZKV samples GSM3455735, GSM3455736, GSM3455737, and saline samples GSM3455729, GSM3455730, GSM3455731 were retrieved from the Gene Expression Omnibus (GEO) with the GEOquery package. The rabies dataset was retrieved in the same manner, GSE38975, samples GSM953148, GSM953149, GSM953150, GSM953151. Microarray data was analyzed with code from NCBI’s GEO2R with the Limma^53^ and umap^55^ packages.

## Acknowledgements

The author(s) disclose receipt of the following financial support for the research, authorship, and/or publication of this article: Funding by the NIH [R01 AI157247 (A.A.K, C.A.H, D.P.B); the BIO5 Institute, University of Arizona (A.A.K.). The funders had no role in the study design; data collection, analysis, or interpretation; or the decision to submit the work for publication.

---

**Figure S1.**
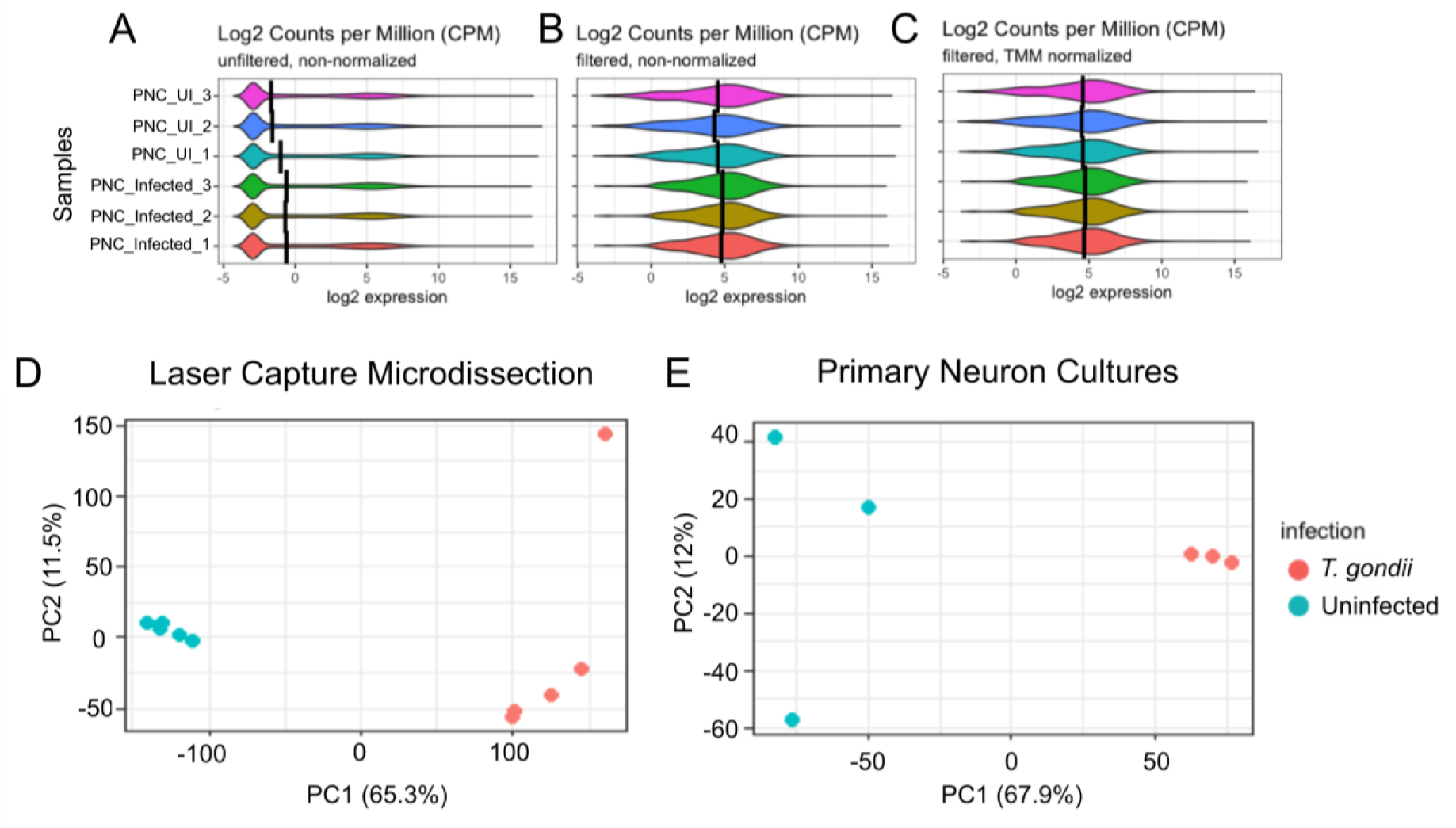
Normalization and data preparation in Primary Neuron Culture (PNC) analysis. (**A**) Unfiltered, non-normalized counts per million (CPM) in PNC samples. UI = uninfected. (**B**) Filtered, non-normalized CPM in PNC samples. (**C**) Filtered, TMM normalized CPM in PNC samples. (**D**) Principal component analysis of PNC. (**E**) Principal component analysis of LCM neurons.

**Figure S2.**
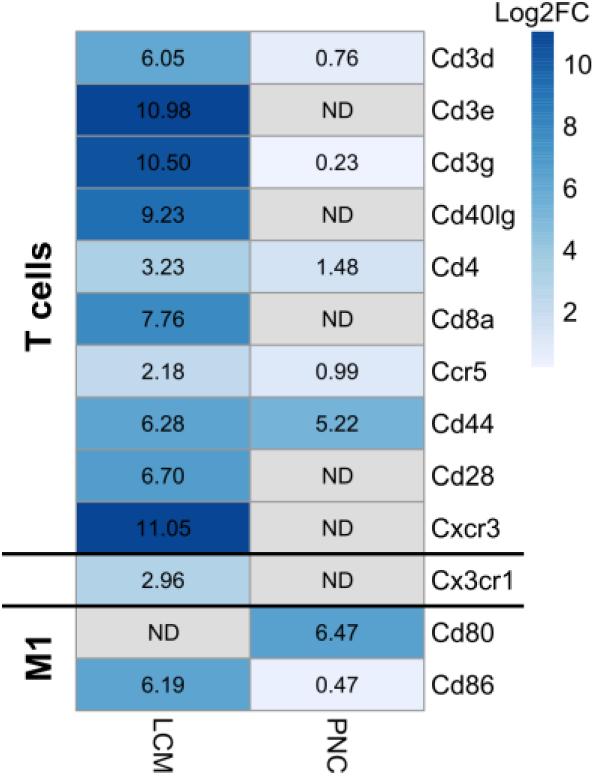
Heat map showing that immune cell transcripts are only observed in LCM dataset. The numbers denote the fold change between infected and uninfected samples. “ND” no differential expression between infected and uninfected datasets. *Cd44* and *Cd80* are immune cell receptors that are also expressed in neurons during development^28,29^.

**Figure S3.**
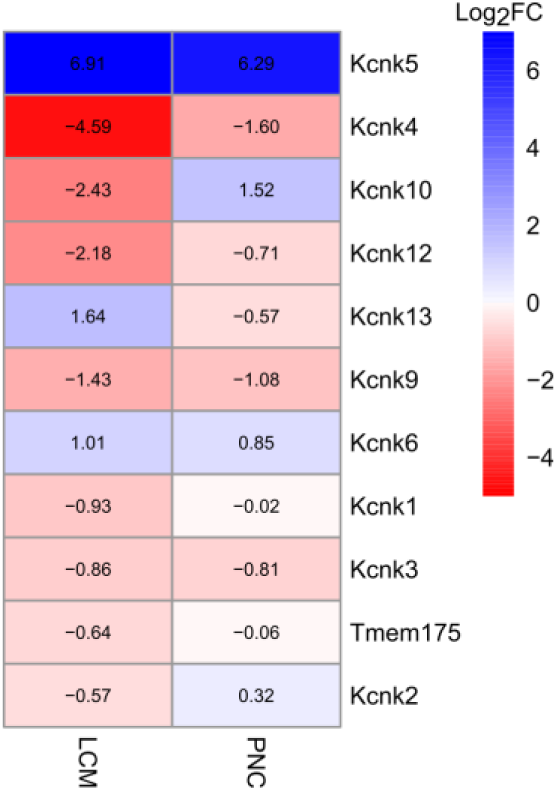
Potassium leak channels are downregulated in LCM and PNCs. Heatmap of genes encoding potassium leak channels shows a downregulation across *T. gondii* datasets. Previous data suggests decreased potassium leak channels result in a depolarized resting membrane potential in TINs^35^.

**Figure S4.**
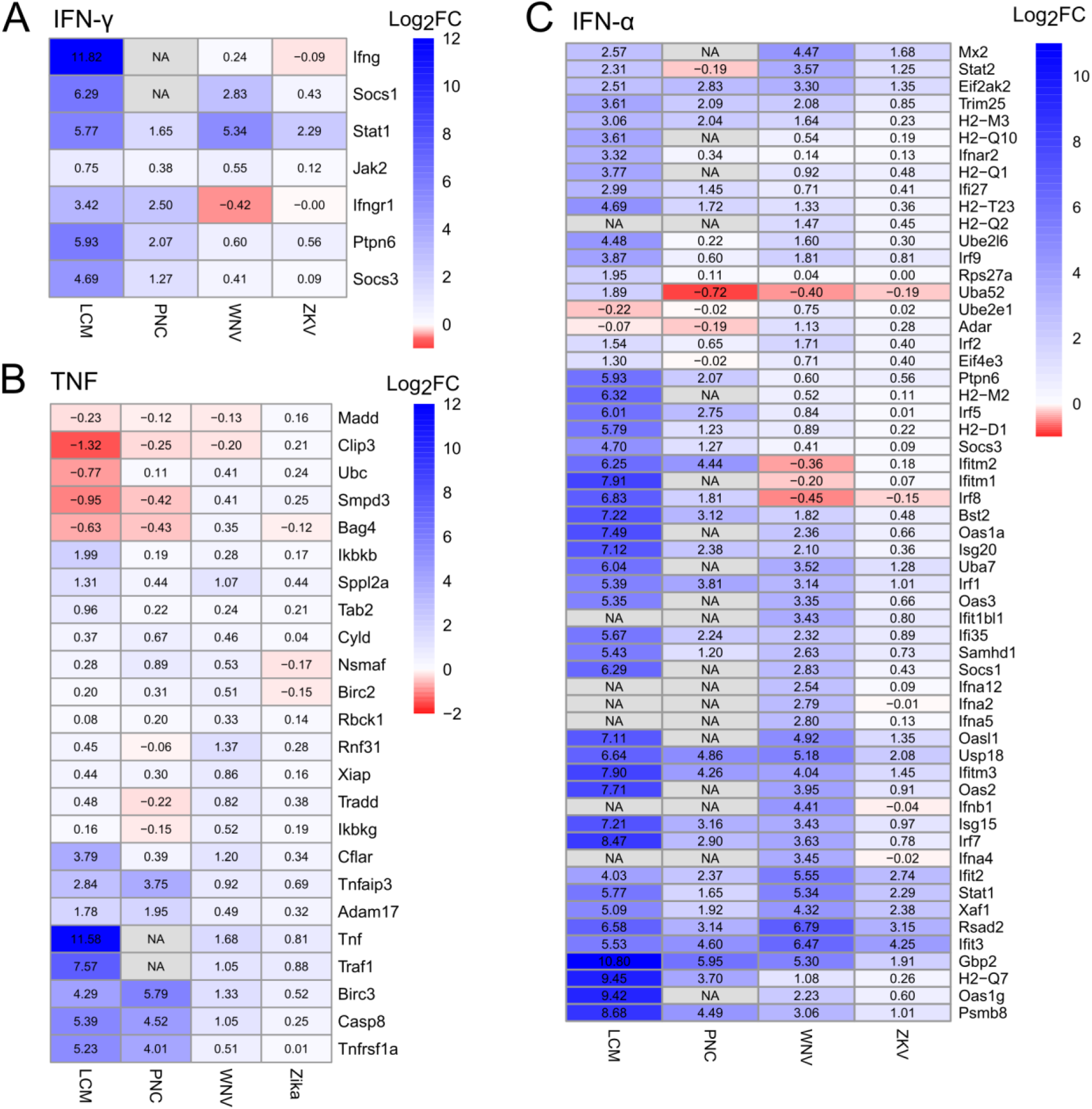
IFN-γ, TNF, and IFN-α pathways are differentially upregulated between datasets. (**A**) Heatmap of IFN-γ genes. (**B**) Heatmap of TNF genes. (**C**) Heatmap of IFN-α genes. Scale = Log2FC. NA = Not Applicable/Not Detected.

