## Supplementary figures and images for "Defining neuronal responses to the neurotropic parasite *Toxoplasma gondii*"

### Supplemental Figure 1

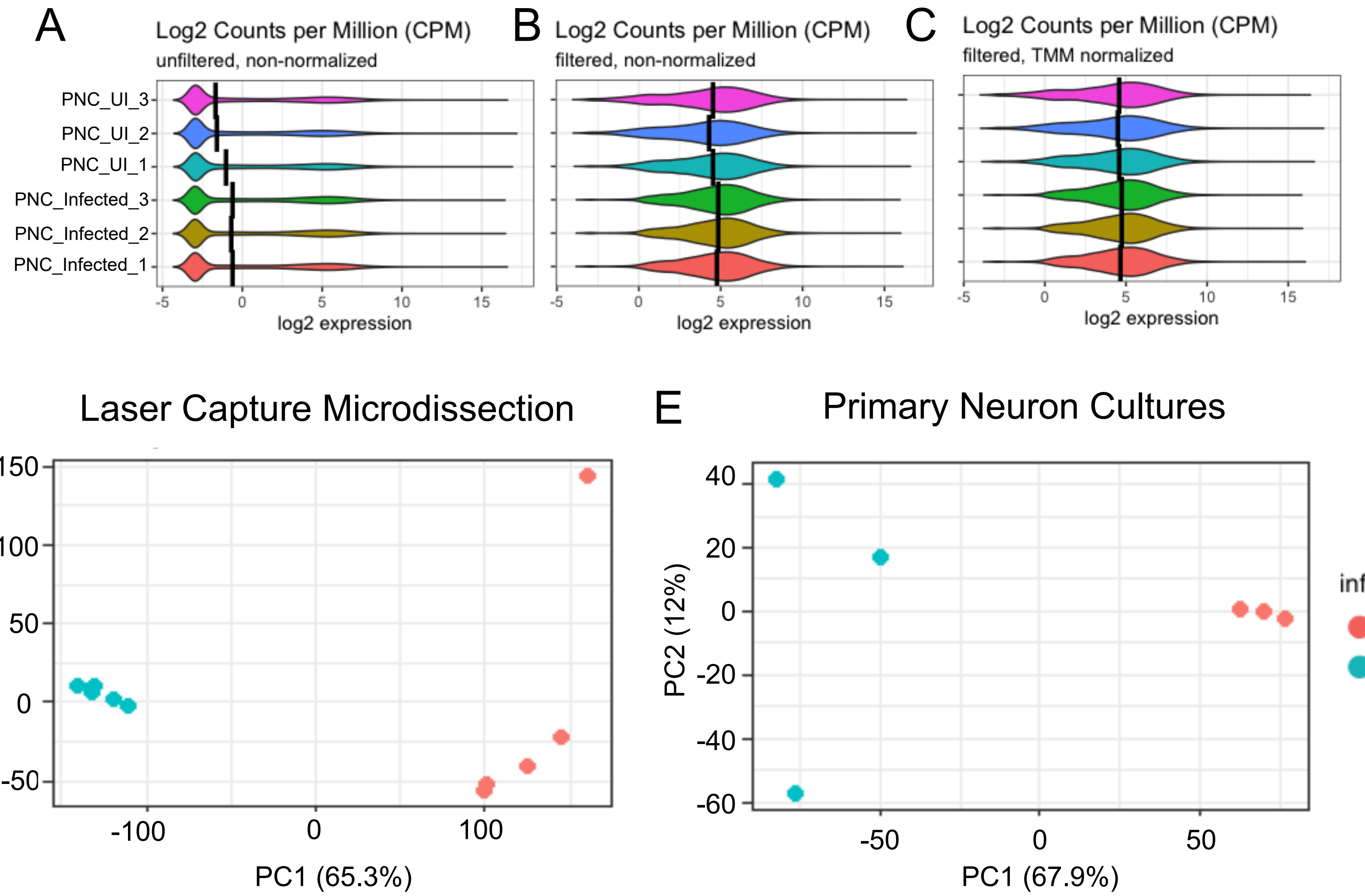
