## Supplemental Figure 2 for "Defining neuronal responses to the neurotropic parasite *Toxoplasma gondii*"

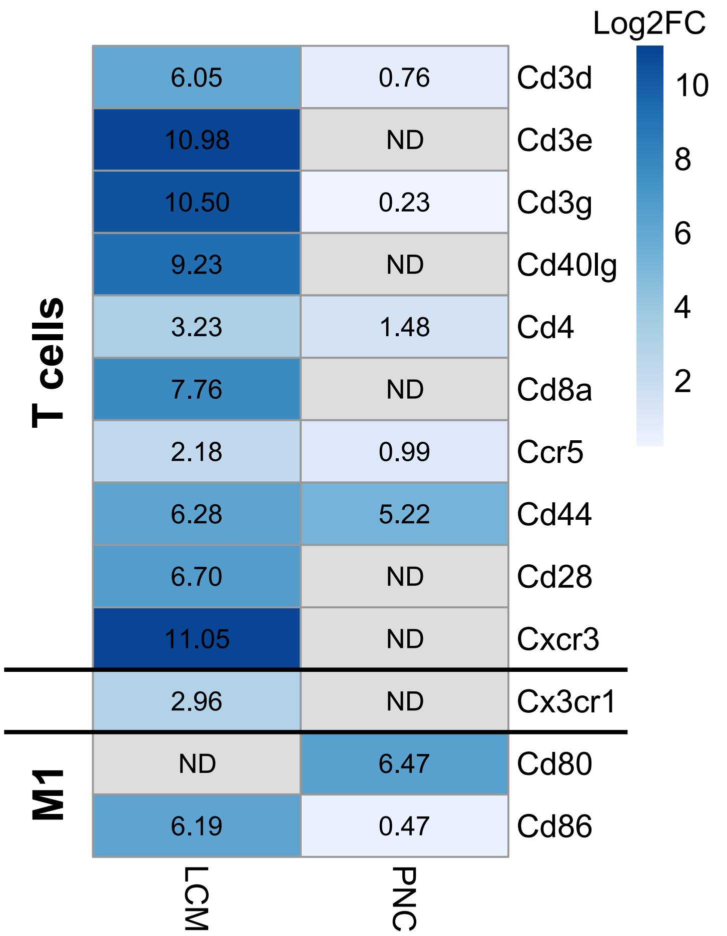


**Figure S2. Heat map showing that immune cell transcripts are only observed in LCM dataset.** The numbers denote the fold change between infected and uninfected samples. "ND" no differential expression between infected and uninfected datasets. Cd44 and Cd80 are immune cell receptors that are also expressed in neurons during development[28,29].
