## Supplemental Figure 3 for "Defining neuronal responses to the neurotropic parasite *Toxoplasma gondii*"

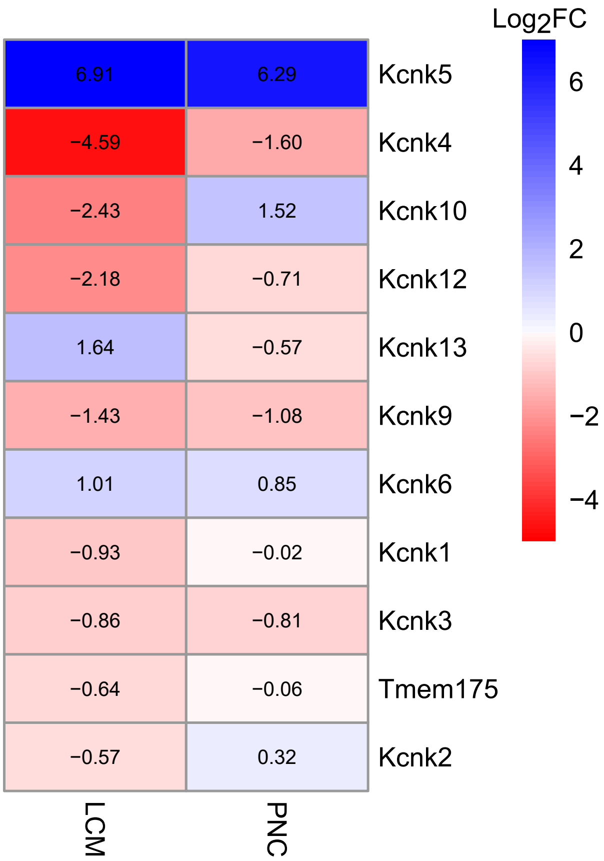


**Figure S3. Potassium leak channels are downregulated in LCM and PNCs.** Heatmap of genes encoding potassium leak channels shows a downregulation across T. gondii datasets. Previous data suggests decreased potassium leak channels result in a depolarized resting membrane potential in TINs[35].
