## Supplemental Figure 4 for "Defining neuronal responses to the neurotropic parasite *Toxoplasma gondii*"

### A IFN-γ

Log2FC

| Gene | LCM | PNC | WNV | ZKV |
|---|---|---|---|---|
| Ifng | 11.82 | NA | 0.24 | −0.09 |
| Socs1 | 6.29 | NA | 2.83 | 0.43 |
| Stat1 | 5.77 | 1.65 | 5.34 | 2.29 |
| Jak2 | 0.75 | 0.38 | 0.55 | 0.12 |
| Ifngr1 | 3.42 | 2.50 | −0.42 | −0.00 |
| Ptpn6 | 5.93 | 2.07 | 0.60 | 0.56 |
| Socs3 | 4.69 | 1.27 | 0.41 | 0.09 |

### B TNF

Log2FC

| Gene | LCM | PNC | WNV | Zika |
|---|---|---|---|---|
| Madd | −0.23 | −0.12 | −0.13 | 0.16 |
| Clip3 | −1.32 | −0.25 | −0.20 | 0.21 |
| Ubc | −0.77 | 0.11 | 0.41 | 0.24 |
| Smpd3 | −0.95 | −0.42 | 0.41 | 0.25 |
| Bag4 | −0.63 | −0.43 | 0.35 | −0.12 |
| Ikbkb | 1.99 | 0.19 | 0.28 | 0.17 |
| Sppl2a | 1.31 | 0.44 | 1.07 | 0.44 |
| Tab2 | 0.96 | 0.22 | 0.24 | 0.21 |
| Cyld | 0.37 | 0.67 | 0.46 | 0.04 |
| Nsmaf | 0.28 | 0.89 | 0.53 | −0.17 |
| Birc2 | 0.20 | 0.31 | 0.51 | −0.15 |
| Rbck1 | 0.08 | 0.20 | 0.33 | 0.14 |
| Rnf31 | 0.45 | −0.06 | 1.37 | 0.28 |
| Xiap | 0.44 | 0.30 | 0.86 | 0.16 |
| Tradd | 0.48 | −0.22 | 0.82 | 0.38 |
| Ikbkg | 0.16 | −0.15 | 0.52 | 0.19 |
| Cflar | 3.79 | 0.39 | 1.20 | 0.34 |
| Tnfaip3 | 2.84 | 3.75 | 0.92 | 0.69 |
| Adam17 | 1.78 | 1.95 | 0.49 | 0.32 |
| Tnf | 11.58 | NA | 1.68 | 0.81 |
| Traf1 | 7.57 | NA | 1.05 | 0.88 |
| Birc3 | 4.29 | 5.79 | 1.33 | 0.52 |
| Casp8 | 5.39 | 4.52 | 1.05 | 0.25 |
| Tnfrsf1a | 5.23 | 4.01 | 0.51 | 0.01 |

### C IFN-α

Log2FC

| Gene | LCM | PNC | WNV | ZKV |
|---|---|---|---|---|
| Mx2 | 2.57 | NA | 4.47 | 1.68 |
| Stat2 | 2.31 | −0.19 | 3.57 | 1.25 |
| Eif2ak2 | 2.51 | 2.83 | 3.30 | 1.35 |
| Trim25 | 3.61 | 2.09 | 2.08 | 0.85 |
| H2−M3 | 3.06 | 2.04 | 1.64 | 0.23 |
| H2−Q10 | 3.61 | NA | 0.54 | 0.19 |
| Ifnar2 | 3.32 | 0.34 | 0.14 | 0.13 |
| H2−Q1 | 3.77 | NA | 0.92 | 0.48 |
| Ifi27 | 2.99 | 1.45 | 0.71 | 0.41 |
| H2−T23 | 4.69 | 1.72 | 1.33 | 0.36 |
| H2−Q2 | NA | NA | 1.47 | 0.45 |
| Ube2l6 | 4.48 | 0.22 | 1.60 | 0.30 |
| Irf9 | 3.87 | 0.60 | 1.81 | 0.81 |
| Rps27a | 1.95 | 0.11 | 0.04 | 0.00 |
| Uba52 | 1.89 | −0.72 | −0.40 | −0.19 |
| Ube2e1 | −0.22 | −0.02 | 0.75 | 0.02 |
| Adar | −0.07 | −0.19 | 1.13 | 0.28 |
| Irf2 | 1.54 | 0.65 | 1.71 | 0.40 |
| Eif4e3 | 1.30 | −0.02 | 0.71 | 0.40 |
| Ptpn6 | 5.93 | 2.07 | 0.60 | 0.56 |
| H2−M2 | 6.32 | NA | 0.52 | 0.11 |
| Irf5 | 6.01 | 2.75 | 0.84 | 0.01 |
| H2−D1 | 5.79 | 1.23 | 0.89 | 0.22 |
| Socs3 | 4.70 | 1.27 | 0.41 | 0.09 |
| Ifitm2 | 6.25 | 4.44 | −0.36 | 0.18 |
| Ifitm1 | 7.91 | NA | −0.20 | 0.07 |
| Irf8 | 6.83 | 1.81 | −0.45 | −0.15 |
| Bst2 | 7.22 | 3.12 | 1.82 | 0.48 |
| Oas1a | 7.49 | NA | 2.36 | 0.66 |
| Isg20 | 7.12 | 2.38 | 2.10 | 0.36 |
| Uba7 | 6.04 | NA | 3.52 | 1.28 |
| Irf1 | 5.39 | 3.81 | 3.14 | 1.01 |
| Oas3 | 5.35 | NA | 3.35 | 0.66 |
| Ifit1bl1 | NA | NA | 3.43 | 0.80 |
| Ifi35 | 5.67 | 2.24 | 2.32 | 0.89 |
| Samhd1 | 5.43 | 1.20 | 2.63 | 0.73 |
| Socs1 | 6.29 | NA | 2.83 | 0.43 |
| Ifna12 | NA | NA | 2.54 | 0.09 |
| Ifna2 | NA | NA | 2.79 | −0.01 |
| Ifna5 | NA | NA | 2.80 | 0.13 |
| Oasl1 | 7.11 | NA | 4.92 | 1.35 |
| Usp18 | 6.64 | 4.86 | 5.18 | 2.08 |
| Ifitm3 | 7.90 | 4.26 | 4.04 | 1.45 |
| Oas2 | 7.71 | NA | 3.95 | 0.91 |
| Ifnb1 | NA | NA | 4.41 | −0.04 |
| Isg15 | 7.21 | 3.16 | 3.43 | 0.97 |
| Irf7 | 8.47 | 2.90 | 3.63 | 0.78 |
| Ifna4 | NA | NA | 3.45 | −0.02 |
| Ifit2 | 4.03 | 2.37 | 5.55 | 2.74 |
| Stat1 | 5.77 | 1.65 | 5.34 | 2.29 |
| Xaf1 | 5.09 | 1.92 | 4.32 | 2.38 |
| Rsad2 | 6.58 | 3.14 | 6.79 | 3.15 |
| Ifit3 | 5.53 | 4.60 | 6.47 | 4.25 |
| Gbp2 | 10.80 | 5.95 | 5.30 | 1.91 |
| H2−Q7 | 9.45 | 3.70 | 1.08 | 0.26 |
| Oas1g | 9.42 | NA | 2.23 | 0.60 |
| Psmb8 | 8.68 | 4.49 | 3.06 | 1.01 |

**Figure S4. IFN-γ, TNF, and IFN-α pathways are differentially upregulated between datasets.** (A) Heatmap of IFN-γ genes. (B) Heatmap of TNF genes. (C) Heatmap of IFN-α genes. Scale = Log2FC. NA = Not Applicable/Not Detected.
